# General and specific patterns of cortical gene expression as spatial correlates of complex cognitive functioning

**DOI:** 10.1101/2023.03.16.532915

**Authors:** Joanna E. Moodie, Sarah E. Harris, Mathew A. Harris, Colin R. Buchanan, Gail Davies, Adele Taylor, Paul Redmond, David Liewald, Maria del C Valdés Hernández, Susan Shenkin, Tom C. Russ, Susana Muñoz Maniega, Michelle Luciano, Janie Corley, Aleks Stolicyn, Xueyi Shen, Douglas Steele, Gordon Waiter, Anca Sandu-Giuraniuc, Mark E. Bastin, Joanna M. Wardlaw, Andrew McIntosh, Heather Whalley, Elliot M. Tucker-Drob, Ian J. Deary, Simon R. Cox

## Abstract

Gene expression varies across the brain. This spatial patterning denotes specialised support for particular brain functions. However, the way that a given gene’s expression fluctuates across the brain may be governed by general rules. Quantifying patterns of spatial covariation across genes would offer insights into the molecular characteristics of brain areas supporting, for example, complex cognitive functions. Here, we use principal component analysis to separate general and unique gene regulatory associations with cortical substrates of cognition. We find that the region-to-region variation in cortical expression profiles of 8235 genes covaries across two major principal components : gene ontology analysis suggests these dimensions are characterised by downregulation and upregulation of cell-signalling/modification and transcription factors. We validate these patterns out-of-sample and across different data processing choices. Brain regions more strongly implicated in general cognitive functioning (*g*; 3 cohorts, total meta-analytic *N =* 39,519) tend to be more balanced between downregulation and upregulation of both major components (indicated by regional component scores). We then identify a further 41 genes as candidate cortical spatial correlates of *g*, beyond the patterning of the two major components (|β| range = 0.15 to 0.53). Many of these genes have been previously associated with clinical neurodegenerative and psychiatric disorders, or with other health-related phenotypes. The results provide insights into the cortical organisation of gene expression and its association with individual differences in cognitive functioning.

## Introduction

In any given cell, genes that are required for that cell’s function are expressed. Therefore, it is tenable that observed regional variations in the expression of genes across the brain reflect location-pertinent cellular processes critical for functioning. Information about regional gene expression profiles across the cerebral cortex has been recently used to infer substrates of brain development, maintenance, and health (^1, 2, 3, 4^). This is achieved by comparing the spatial agreement between the brain regional expression profiles of individual genes or gene sets with the brain regional associations with a phenotype of interest. For example, which specific genes or gene sets are more highly expressed in brain regions that are most strongly related to a particular phenotype of interest (^5^)? This approach, while powerful, potentially suffers from confounding by association. That is, for example, the expression of an individual gene might show a correlation with a phenotype because it reflects general rules that govern the spatial variation in the expression of very many genes over the brain’s cortex, rather than something unique to the specific gene in question. There are general dimensions of spatial variation in gene expression covariance, demonstrating shared covariation in expression patterns across multiple genes, across the human body (^6^), and within multiple organs (^7^). This includes across the human cortex (^8^), where general dimensions of gene expression have previously been linked to in-vivo MRI estimates of cortical structural anatomy (^9^), and to functional MRI-derived neurocognitive associations (^10, 11^). It is therefore critical to control for general dimensions along which regional variation in gene expression covary when seeking gene-specific associations. Because much of our information on gene expression patterns in the brain (with sufficient regional fidelity for these questions) comes from relatively few donors, it is also critical to seek out-of-sample replication. Here, we unite micro-(gene expression) and macro-level (morphometry) information about the brain, to inform the underlying molecular neurobiology of complex cognitive functioning.

General cognitive functioning, or ‘*g*’, is a robust and well-replicated index of individual differences in cognitive functioning, capturing variance in reasoning, planning, problem-solving, some aspects of memory, processing speed and abstract thinking (^12, 13^). It is associated with educational attainments (^14^), life achievements (^15^), health (^16^). And lifespan(^17, 18^). Regions of the brain proposed to support general cognitive functioning, or ‘*g*’ (and which relate to individual differences therein), have been identified via an array of methods including resting state fMRI (^19^), structural and functional connectivity (^20^), lesion studies (^21^), post mortem brain studies (^22^), and genetic information (^23^). These brain regions overlap substantially with those associated with other summary cognitive constructs, such as executive functioning (^24^). Macrostructural cortical measures provide some convergent evidence for a specific patterning of brain regional *g-* correlates, particularly highlighting parieto-frontal regions (^25, 26^). However, debate remains about the loci of *g*’s cortical correlates, for which large multi-cohort analyses are required (^27^). Specifically, there is uncertainty in how much overlap there is in the spatial patterns of *g* associations with cortical thickness and surface area, measures which are largely phenotypically and genetically distinct (^28, 29, 30, 31^).

Here, we combine i) *post mortem* gene expression data and ii) the largest meta-analysis of the cortico-macrostructural correlates (*in vivo* MRI; cortical volume, surface area and thickness) of individual differences in cognitive functioning to-date. Both are available at the same level of granularity with respect to brain regions, allowing us to quantitatively assess spatial associations between cortical gene expression and general cognitive function. Therefore, we can ask this new question: is there an association between variation in gene expression across different brain areas and how strongly brain structural measures are associated with cognitive functioning in those same brain regions? I.e. does the brain regional map of gene expression resemble the brain regional map of brain structure-cognitive function correlations?

In contrast to prior work looking at the cortical expression patterns of single candidate genes or gene types, we discover that the expression of 8235 genes varies together in a synchronised fashion across the cerebral cortex. Two major components account for the majority (49.4%) of the variance in regional gene expression profiles, representing a cell-signalling/modifications axis and a transcription factors axis. We address the potential limitations of having only *N* = 6 tissue donors and one regional sampling approach: the dimensions of gene expression are validated in two independent gene expression atlases (*N* = 5, *N* = 11 tissue donors), and are not driven by a small number of individual outlier regions. Similarly, our meta-analysis of associations between *g* and regional cortical morphometry (volume, surface area, thickness), across 3 cohorts (total *N* = 39,519), shows good cross-cohort consistency in regional mapping.

The patterning of *g*-associations with brain structural measures across the cortex are associated with both of the identified gene expression components, with medium-to-large effect sizes for *g-*volume and *g-*surface area associations but weaker ones for *g-* thickness. We further identify 41 single genes whose expression patterns are individually associated with *g-*cortical profiles beyond the two major dimensions of cortical gene expression. Thus, this study provides clarity on the patterning and replicability of the brain-macrostructural correlates of cognitive functioning differences, and identifies novel regional global and specific gene expression patterns that might govern them.

## Methods and materials

### Gene expression method

The Allen Human Brain Atlas is a high-resolution mapping of cortical gene expression for *N =* 6 donors (5 male, 1 female, age *M =* 42.50 years, *SD =* 13.38 years, range = 24-57 years). The complete microarray data from a custom-designed Agilent array for all 6 donors are openly available for download. French and Paus(^54^) summarised these data to the Desikan-Killiany cortical atlas. To briefly summarise their method (for more information, refer to the original paper and *Table S6*), gene expression values were averaged across multiple probes. Each of the 3702 brain samples was assigned to one of the 68 Desikan-Killany regions based on their MNI coordinates, and then gene expression values were averaged per region, resulting in an expression value for each gene for each region. These between-donor median expression values are publicly available (^32^). French and Paus also provide a method of quality control for between-donor consistency in regional gene expression profiles (^54^). In this method, profiles with Spearman’s ρ > 0.446 (equivalent to one-sided *p <* .05) between the average of donor-to-median left hemisphere profile correlations are considered to have high between-donor consistency. This method results in the retention of 8325 out of 20737 genes.

The right hemisphere expression data are based on a maximum of *N =* 2 donors, compared to a maximum of *N =* 6 donors for the left hemisphere. The number of samples per region is lower in the in right hemisphere (*M =* 12.59, *SD =* 8.90, range 2-34) than the left hemisphere (*M =* 37.32, SD = 24.37, range = 6-100). Further details of the number of samples and donors per region are in *Table S2*. The donor-level expression data are not available, so in the present study the 8235 genes that passed the quality control protocol in the left hemisphere were also analysed for the right. There is a strong correlation between the expression values of individual genes between hemispheres (*r =* 0.997, *p <* 2.2e-16) suggesting that, at the hemisphere level, the relative expression values for the 8235 genes were not affected by the sampling differences between the two hemispheres.

We conducted PCA on the median gene expression values of 8235 genes across 68 cortical regions – rows = cortical regions (in place of participants in a traditional PCA) and columns = genes. We performed extensive checks for the validity of the first two components – these are detailed in the Results section.

In the raw data, the right hemisphere has lower average expression values than the left hemisphere (right: *M =* 6.035, *SD =* 2.343, left: *M =* 6.091, *SD =* 2.368), *t*(58.345) = 6.490, *p =* 2.051e-08), an artefact of there being a maximum of *N =* 2 donors for the right hemisphere compared to a maximum of *N =* 6 donors for the left. This artefact creates a clear hemisphere difference centred around zero in component scores for both components: Component 1: *t*(65.931 = 7.794), *p =* 6.218e-11, *M left* = −0.687 (*SD =* 0.715), *M right* = 0.687 (*SD =* 0.739), Component 2: *t*(65.931) = −5.315, *p =* 1.388e-06, *M left* = −0.543 (*SD =* 0.798), *M right* = 0.543 (*SD =* 0.886). However, there was a strong interhemispheric correlation in scores between the 34 paired regions for both Component 1 (*r =* 0.815, *p =* 4.411e-09) and Component 2 (*r =* 0.725, *p =* 1.25e-06).

To confirm it was appropriate to treat these hemispheric differences as an artefact of the data, and thus scale the component scores in each hemisphere, we looked to the Kang et al. (^33^) dataset. In this dataset, there was a more even number of donors per hemisphere (left hemisphere *M =* 9.55, *SD =* 1.04 donors per region, right hemisphere region *M =* 7.19, *SD =* 0.60 donors per region), and there was no difference in means expression values per hemisphere *t*(19.344) = −0.852, *p =* .405, *M left* = 7.521 (*SD =* 1.944), *M right* = 7.535 (*SD =* 1.943). For the rotated scores of PC1 (which had a factor congruence of 0.96 with the French and Paus expression matrix), scores were comparable between hemispheres *t*(19.997) = −0.265, *p =* .794. Therefore, we deemed it appropriate to scale the component scores separately for each hemisphere in the current dataset (see Figure S2).

#### Statistical overrepresentation analysis

To assist with interpretation of the two identified major components of gene expression, PANTHER’s protein analysis and GO-Slim molecular, biological and cellular (version 16.0, released 2020-12-01) terms were analysed. All genes included in the PCA were submitted as a reference set for the statistical overrepresentation analysis and 7389 out of 8235 (89%) genes were available in PANTHER, and so were used as the background set. Fisher’s exact test and FDR correction were used, and four subsets of genes were tested for statistical overrepresentation: Component 1 loadings < −0.3 (total *N =* 3371, available *N =* 3099, 92%) and loadings > 0.3 (total *N =* 2093, available *N =* 2000, 96%); and Component 2 loadings < −0.3 (total *N =* 3477, available *N =* 3234, 93%) and loadings > 0.3 (total *N =* 1706, available *N =* 1551, 91%).

The statistical overrepresentation results are provided in full in a supplementary data file. Some genes have absolute loadings > 0.3 on both components (*N =* 3026, 36.75%). There are also a number of genes that had absolute loadings > 0.3 only on either Component 1 (*N =* 2438, 29.61%) or Component 2 (*N =* 2157, 26.19%). *N =* 614 genes (7.46%) did not load with an absolute > 0.3 on either component, and all statistical overrepresentation tests for this set were null.

For the two components, a gene set enrichment analysis was run in FUMA (Functional Mapping and Annotation of Genome-Wide Association Studies, https://fuma.ctglab.nl/). Hypergeometric tests were performed to test if genes of interest were overrepresented in any of the pre-defined gene sets (those with absolute loadings > 0.3 on each component), with the 8235 genes as a background set. No significant (α < .05) gene sets were obtained from the reported genes in the GWAS catalog.

### *g* ∼ cortical morphometry associations meta-analysis method

#### Cohorts

The UK Biobank (UKB, http://www.ukbiobank.ac.uk, ^34^) holds data from ∼500,000 participants, and for ∼40,000 at wave 2 of data collection, data includes head MRI scans and cognitive test data. In the current study, we did not include participants if their medical history, taken by a nurse at the data collection appointment, recorded a diagnosis of e.g. dementia, Parkinson’s disease, stroke, other chronic degenerative neurological problems or other demyelinating conditions, including multiple sclerosis and Guillain– Barré syndrome, and brain cancer or injury (a full list of exclusion criteria is listed in the *Supplementary tabular data file*, and see Figure S3 for *N* by exclusion condition). After these exclusions, the final study included *N =* 37,840 participants (53% female), age *M =* 63.81 years (*SD =*7.64 years), range = 44-83 years. The UKB was given ethical approval by the NHS National Research Ethics Service North West (reference 11/NW/0382). The current analyses were conducted under UKB application number 10279. All participants provided informed consent. More information on the consent procedure can be found at https://biobank.ctsu.ox.ac.uk/crystal/label.cgi?id=100023.

STRADL is a population-based study, developed from the Generation Scotland Scottish Family Health Study. Participants who had taken part in the Generation Scotland Scottish Family Health Study were invited back to take part in this additional study, which was initially designed to study major depressive disorder, although participants were not selected based on the presence of depression (^35^) https://www.research.ed.ac.uk/en/datasets/stratifying-resilience-and-depression-longitudinally-stradl-a-dep. Data are available for *N =* 1188 participants. The current sample includes *N =* 1043 participants, for whom both MRI head scans and cognitive data are available (60% female), age *M =* 59.29 years (*SD =* 10.12 years), range = 26-84 years. STRADL received ethical approval from the NHS Tayside Research ethics committee (reference 14/SS/0039), and all participants provided written informed consent.

The LBC1936 is a longitudinal study of a sample of community-dwelling older adults most of whom took part in the Scottish Mental Survey of 1947 at ∼11 years old, and who volunteered to participate in this cohort study at ∼70 years old (^36, 37^) https://www.ed.ac.uk/lothian-birth-cohorts. The current analysis includes data from the second wave of data collection, which is the first wave at which head MRI scans are available, in addition to cognitive tests. In total, 731 participants agreed to MRI scanning. After image processing, data were available from *N =* 636 participants (47% female), age *M =* 72.67 years, *SD =* 0.41 years, range = 70–74 years. The LBC1936 study was given ethical approval by the Multi-Centre Research Ethics Committee for Scotland, (MREC/01/0/56), the Lothian Research Ethics Committee (LREC/2003/2/29) and the Scotland A Research Ethics Committee (07/MRE00/58). All participants gave written consent before cognitive and MRI measurements were collected.

#### MRI protocols

Detailed information for MRI protocols in all three cohorts are reported elsewhere: UKB (^38^), LBC1936 (^39^) and STRADL ^40^, but are briefly summarised here. In the present sample, UKB participants attended one of four testing sites: Cheadle (*N =* 22,636, 60%), Reading (*N =* 5463, 14%), Newcastle (*N =* 9526, 25%), and Bristol (*N =* 51, 0.14%).The same type of scanner was used in all four testing sites, a 3T Siemens Skyra, with a 32-channel Siemens head radiofrequency coil. The UKB MRI protocol includes various MRI acquisitions (more details available here https://www.fmrib.ox.ac.uk/ukbiobank/protocol/V4_23092014.pdf) but relevant to this work are the T1-weighted MPRAGE and T2-FLAIR volumes. For T1-weighted images, 208 sagittal slices were acquired with a field view of 256 mm and a matrix size of 256 x 256 pixels, giving a resolution of 1 x 1 x 1 mm^3^. The repetition time was 3.15 ms and the echo time was 1.37 ms.

STRADL had 2 testing sites: Aberdeen (in the present sample, *N =* 528, 51%) and Dundee (*N =* 515, 49%). Detailed information about the STRADL structural image acquisitions are available here https://wellcomeopenresearch.org/articles/4-185. For the current analysis, we used the T1-weighted fast gradient echo with magnetisation preparation volume sequence. The Aberdeen site used a 3T Philips Achieva TX-series MRI system (Philips Healthcare, Best, Netherlands) with a 32-channel phased-array head coil and a back facing mirror (software version 5.1.7; gradients with maximum amplitude 80 mT/m and maximum slew rate 100 T/m/s). For T1-weighted images, 160 sagittal slices were acquired with a field of view of 240 mm and a matrix size of 240 x 240 pixels, giving a resolution of 1 x 1 x 1 mm^3^. Repetition time was 8.2 ms, echo time was 3.8 ms and inversion time was 1031 ms. In Dundee, the scanner was a Siemens 3T Prisma-FIT (Siemens, Erlangen, Germany) with 20 channel head and neck phased array coil and a back facing mirror (Syngo E11, gradient with max amplitude 80 mT/m and maximum slew rate 200 T/m/s). For T1-weighted images 208 sagittal slices were acquired with a field of view of 256 mm and matrix size 256 x 256 pixels giving a resolution of 1 x 1 x 1 mm^3^. Repetition time was 6.80 ms, echo time was 2.62 ms, and inversion time was 900 ms.

All LBC1936 participants were scanned in the same scanner at the Brain Research Imaging Centre, Western General Hospital, Edinburgh, using a GE Signa LX 1.5T Horizon HDx clinical scanner (General Electric, Milwaukee, WI) with a manufacturer supplied 8-channel phased array head coil. More information on the structural image acquisitions for the LBC1936 cohort is available in (^39^). For T1-weighted images (3D IR-Prep FSPGR), 160 coronal slices were acquired, with a field of view of 256 mm and a matrix size of 192 x 192 pixels giving a resolution of 1 x 1 x 1.3 mm^3^. The repetition time was 10 ms, echo time was 4 ms and inversion time was 500 ms.

For all cohorts, the FreeSurfer image analysis suite (http://surfer.nmr.mgh.harvard.edu/) was used for cortical reconstruction and volumetric segmentation. The Desikan-Killany atlas parcellation yields 34 paired regional measures in left and right cortical hemispheres (^41^). Different versions of FreeSurfer were used in the three cohorts (UKB = v6.0, STRADL = v5.3, LBC1936 = v5.1), and only for UKB were T2-FLAIR volumes used to improve the pial surface reconstruction. The LBC1936 and STRADL parcellations have previously undergone thorough quality control, with manual editing to rectify any issues. Manual edits were performed to ensure correct skull stripping, tissue identification and positioning of cortical regional boundary lines. The UKB regional data were extracted from the aparc.stats files and these parcellations have not been manually or automatically edited. For the current study, UKB values more than 4 standard deviations from the mean for any individual regional measure were excluded (UKB *M =* 24.28, *SD =* 19.41, range = 0–104 participants per region). For UKB and STRADL cohorts, cognitive and MRI data were collected on the same day, but in LBC1936, there was a slight delay between the two testing sessions (*M =*65.08, *SD =* 37.77 days). Raw values are plotted for mean volume, surface area and thickness by age and cohort in *Figure S6*, and for each region in *Figures S7-13*.

#### Cognitive Tests

All three cohorts have collected data across several cognitive tests, covering several cognitive domains, which enables the estimation of a latent factor of general cognitive functioning *(g)*. The cognitive tests in each cohort have been described in detail elsewhere: UKB (^42^), STRADL (^43^), LBC1936 (^36, 44, 45^). The measures used in the present study are summarised in *Tables* S11-S11. In STRADL and LBC1936, the cognitive data was used as provided, as this data has been pre-cleaned. For UKB, we coded prospective memory from 0 to 1, as suggested in ^46^, for numeric memory, values at −1 were removed (abandoned test) and for Trail B, values at 0 were removed (trail not completed). Reaction time, trail B and pairs matching scores were log transformed.

A latent factor of *g* was estimated for each cohort, using all available cognitive tests, using confirmatory factor analysis in a structural equation modelling framework. Each individual test was corrected for age and sex. Latent *g* model fits were assessed using the following fit indices: Comparative Fit Index (CFI), Tucker Lewis Index (TLI), Root Mean Square Error of Approximation (RMSEA), and the Root Mean Square Residual (SRMR) (for model fits, see *Table S18*). For the LBC1936, *g* has previously been modelled with a hierarchical confirmatory factor analysis approach, to incorporate defined cognitive domains (^47, 48^). Here, in keeping with these previous models, within-domain residual covariances were added for four cognitive domains (Visuospatial skills, Crystalised ability, Verbal memory and Processing speed). Results of the *g* measurement models are summarised in *Tables S14-S17,* and *Figure S5*. For all cohorts, all estimated paths to latent *g* were statistically significant with all *p <* .001.

The latent *g* scores were extracted for all participants. Those for UKB were multiplied by −1 so a higher score reflected better cognitive performance, to match scores from STRADL and LBC1936. Then, for each cohort, a standardised β was estimated between *g* and three measures of cortical morphometry (volume, surface area and thickness) for each of the 68 regions. Cortical measures were controlled by age, sex head position in the scanner (X, Y and Z coordinates), testing site (for UKB and STRADL) and lag between cognitive and MRI appointments (for LBC1936). The resulting standardised β estimates for each region and each measure were meta-analysed between the three cohorts (68 regions x 3 measures = 204 random effects meta-analyses). The full results of these meta-analyses are in *Tables S14-17*.

Although we controlled for age in the *g-*cortical morphometry association models within each cohort, each cohort had different age ranges (with the LBC1936 having a notably narrow age-range of 70-74 years old), and it is possible this might affect the associations. Therefore, we also tested for mean age moderation effects on meta-analytic estimates, and none were significant after FDR correction (all *FDR Q* > .27), see *Tables S22-S24*.

#### Additional analyses

In addition to the main analyses, which focus on *g-*associations with general and specific gene expression profiles, we also ran a parallel supplementary analysis simply on the regional morphometry means (see *Supplementary Text 1*).

### Analysis software

Most analyses were conducted in R 4.0.2. (R Core Team, 2020). The psych package was used for PCAs (^49^), the core R stats package was used for the Kruskal-Wallis tests, the FSA package (^50^) was used for Dunn’s Kruskal-Wallis multiple comparisons, and the metafor (^51^) package was used for the meta-analyses. All structural equation models were estimated in lavaan (^52^) with the full information maximum likelihood method. GO term analyses were conducted at http://geneontology.org/, which is powered by PANTHER 1F^53^. FUMA https://fuma.ctglab.nl/ was used for gene set enrichment analysis for the two components, and previous GWAS associations with allelic status of the specific individual genes-*g* associations were looked up in the GWAS catalog.

### Data Availability

All UKB data analysed herein (including IDPs) were provided under project reference 10279. A guide to access UKB data is available from http://www.ukbiobank.ac.uk/register-apply/. To access data from the STratifying Resilience and Depression Longitudinally (STRADL) study, which is part of the Generation Scotland study, see https://www.research.ed.ac.uk/en/datasets/stratifying-resilience-and-depression-longitudinally-stradl-a-dep, and to access the Lothian Birth Cohort data, see https://www.ed.ac.uk/lothian-birth-cohorts/data-access-collaboration.

## Results

### Two major dimensions of cortical gene expression

The Allen Human Brain Atlas consists of a high-resolution mapping of gene expression to the cerebral cortex for *N =* 6 donors (5 male, 1 female, Age *M =* 42.50 years, *SD =* 13.38 years, range = 24-57 years). French and Paus (^54^) summarised these data across donors to find the average gene expression values for each region in the Desikan-Killiany atlas and provide a method of quality control for between-donor consistency in regional gene expression profiles, which results in retention of 8325 genes (out of 20,737 originally available from the atlas). These retained genes are associated with neural gene ontology (GO) terms, and those not retained tend to have low expression across the cortex or are associated with other GO terms e.g. olfactory receptor and keratin genes (^54^). This results in a gene expression matrix (rows = 68 cortical regions, columns = 8235 genes) of median gene expression values for each region for each gene across donors. Initial results of a principal component analysis (PCA) on these data indicated that regional variation in gene expression across the cortex occurs across very few biological dimensions (see *Figure 1B*); that is, there was much similarity across genes in the patterning of their expression across brain regions. Mindful of the potential limitations of basing a new discovery in fundamental neuroscience on a modest *post mortem* dataset (*N* = 6 donors), we performed extensive checks.

**Figure 1.**
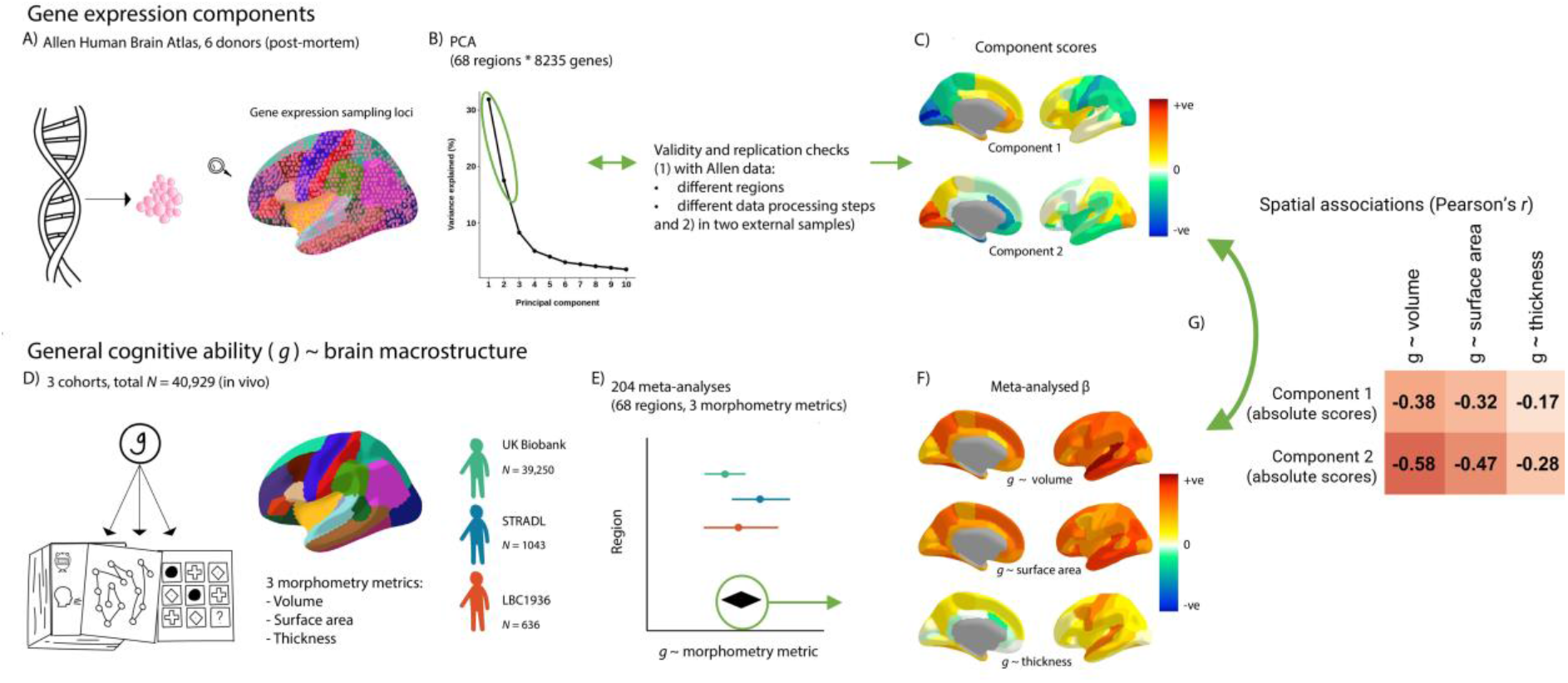
An illustration of the analytic framework. *Figure 1 note* A) Gene expression data from the Allen Human Brain Atlas was summarised to the Desikan-Killiany Atlas. B) We conducted PCA on the gene expression matrices (68 regions * 8235 genes) and two components were justified with validity checks. C) We rotated these two components, and the component scores show the relative positions of the 68 Desikan-Killany regions on these components. D) *g* ∼ brain cortical morphometry associations were calculated for three cohorts. E) The *g* ∼ brain cortical morphometry associations were meta-analysed with random effects models. F) The meta-analysed standardised β values of each regional morphometry metric (cortical volume, surface area and thickness) show their associations with *g.* G) Spatial associations were tested between the brain-regional component scores for gene expression and the regional *g* ∼ brain cortical morphometry associations. Then, controlling for the regional component scores, *g*-associations for individual genes were calculated.

We tested the factor congruence of the resulting principal components in terms of: over-reliance on specific cortical regions, congruence with nine different gene expression data processing pipelines, in two independent samples, and different brain parcellation choices. First, to test the regional dependence of the principal components, we used cross-validation to create 5 random partitions of the 68 regions 50 times without replacement. Each time, the partitions were arranged into two sets, one with ∼54-55 regions (4 of 5 partitions) and the other with ∼13-14 regions (1 of 5 partitions). The PCA was repeated for each iteration (a total of 250 tests). Absolute coefficients of factor congruence between the two sets tended to be high for the first two components (PC1: *M =* 0.926, *SD =* 0.064; PC2: *M =* 0.830, *SD =* 0.092), and were notably weaker, with higher variability, from the third component onwards, see *Figure 2c*. Therefore, the first two components do not rely heavily on individual regions, and so were taken forward in the current analysis. Unrotated, PC1 accounted for 31.9% of the variance, and PC2 for 17.5%, (after varimax rotation PC1 accounts for 25.8% of the variance, and PC2 for 23.6%).

**Figure 2.**
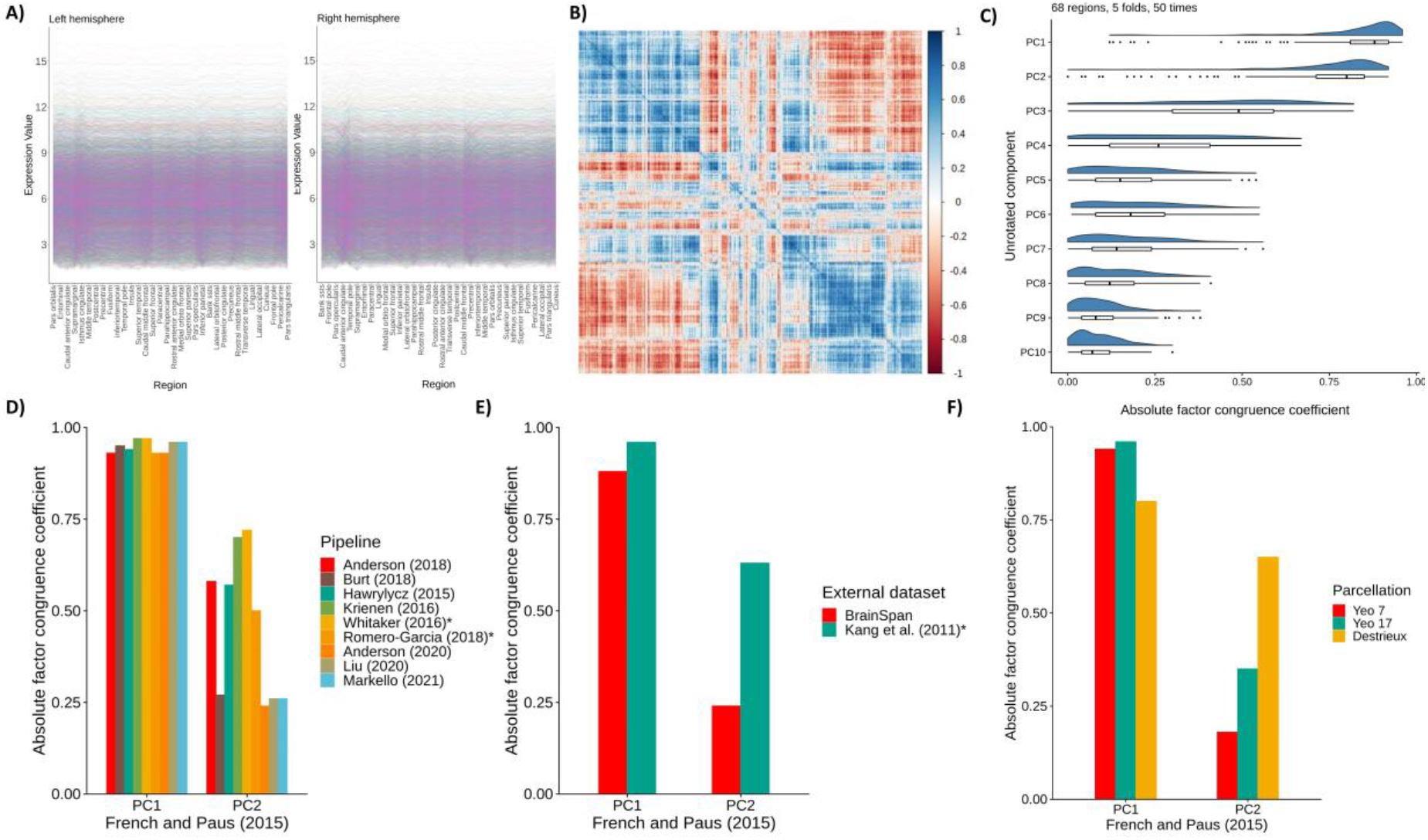
Validating gene expression components. *Figure 2 note* A) Raw gene expression values for the 34 regions for the left and right hemispheres, for the 8235 consistent genes. B) Correlation plot of the 8235 genes across the 68 cortical regions (8235 * 8235). C) Absolute factor congruence coefficients for the first 10 components between “train” and “test” folds (∼54-55 regions, and ∼12-13 regions), over 50 repetitions. D) Absolute factor congruence coefficients from different pipelines with PC1 and PC2 of the current dataset of interest, using the Desikan-Killiany atlas. * denotes that PC3 from that pipeline is compared with PC2. E) Absolute factor congruence coefficients for two external datasets with PC1 and PC2 of the current dataset of interest. * denotes that PC3 from is compared with PC2. F) Absolute factor congruence coefficients for three alternative parcellations with the PC1 and PC2 of the current dataset (which uses the Desikan-Killiany atlas).

Although there are efforts towards developing standardised processing of gene expression data (^55^), there remains no consensus. There have been several proposed pipelines for summarising the Allen Human Brain Atlas data, and so we sought to test whether the gene expression components derived using PCA are valid when the data are summarised with different processing choices. We applied the scripts provided by Markello et al. (55), that reproduce the pipelines for several studies (see *Table S6*). The initial number of retained genes, and the number of genes matched to French and Paus’ post-consistency check genes are in *Table S8*. We investigated whether the two identified components are similar when different methods of summarising the Allen Human Brain Atlas to Desikan-Killiany space (see Figure 2D). To test this, we replicated the gene expression matrix from French and Paus using Markello et al.’s scripts (^55^) and the abagen toolbox (^56^). This replication is not exact, but very close – 8108 genes were retained and the factor congruence coefficient for PC1 = 0.99, and for PC2 = 0.98. We then ran PCAs on the resulting gene expression matrices obtained from nine gene expression data processing pipelines^55^, see *Table S3*. For each pipeline, we calculated factor congruence coefficients with French and Paus’ method based on matched genes. Absolute coefficients ranged from 0.93-0.97 for PC1 and 0.24-0.72 for PC2, see *Figure 2.* There are notable PC2 inconsistencies with particular pipelines - Burt (2018), Anderson (2020), Liu (2020) and Markello (2021) - which are likely due to less common choices such as donor-specific probe selection, stringent interareal similarity filtering thresholds, and other choices that impact the number of genes retained.

We limited the donor age from these additional datasets to be in proximity to the age range of the Allen Human Brain Atlas (24-57 years of age). See *Table S1* for descriptive statistics of the validation samples. The test for between-donor consistency provided by French and Paus (59) was applied to these datasets. Then, PCAs were conducted on the gene expression matrices (rows = cortical regions, columns = genes), and genes were matched with those in the Allen dataset to test for factor congruence. Summaries of the number of retained genes at each step are in *Table S7*.

To test whether the first two components were generalizable beyond the 6 donors from which the Allen Human Brain Atlas data were derived, we sought external validation with two independent datasets, the BrainSpan Atlas https://www.brainspan.org/ and an atlas provided by Kang et al. connected to the Human Brain Transcriptome Project https://hbatlas.org/55F^57^. Both external datasets used the Affymetrix GeneChip Human Exon 1.0 ST Array Platform to summarise gene expression data. and include 11 cortical regions, which have previously been roughly matched to 14 regions in the Desikan Killiany atlas ^58^ (see Figure S1). For BrainSpan (donor *N =* 5), these are collapsed across hemispheres, but for the Kang et al. dataset (donor *N =* 11), they are available for each hemisphere separately (a total of 22 regions). Genes with consistent between-donor profiles were identified, using French and Paus’ procedure ^59^. To test for factor congruence, these were then matched with the 8235 genes that were consistent between donors in the French and Paus dataset, resulting in 2250 genes for the BrainSpan comparison and 908 for Kang et al.. The relatively small numbers of retained genes could be due to different cortical boundaries, extent of cortical coverage or the gene expression measurement and sampling methods used. There was high factor congruence for PC1_Allen_ in both datasets (the coefficient for PC1_BrainSpan =_ 0.88 and PC1_Kang et al._ = 0.96) and low-moderate factor congruency for PC2_Allen_ (with PC2_BrainSpan =_ 0.24, and PC3_Kang et al._ = 0.63 (see *Figure 2E*). PC2_Kang et al._ did not have high factor congruence with any Allen component (the maximum absolute value was 0.19, which was with PC6_Allen_).

Lastly, we tested whether the positioning of regional boundaries affected the consistency of the components (see *Figure 2F*). Three open source atlases were tested: Yeo’s Functional Connectivity *7* and 17 Network atlases (with 7 and 17 regions, respectively) ^60^ and the Destrieux atlas (134 regions, 67 per hemisphere) ^61^. For all three, as with the Desikan-Killiany atlas, 8108 genes matched with the 8235 from the main working dataset. Again, factor congruence coefficients tended to be higher for PC1_Allen_ than PC2_Allen_, (PC1_Yeo7_ = 0.94, PC1_Yeo17_ = 0.96, PC1_Destrieux_ = 0.80; PC2_Yeo7_ = 0.18, PC2_Yeo17_ = 0.35, PC2_Destrieux_ = 0.65). Notably, factor congruence coefficients tended to increase for PC2 with increasing granularity. These results may partially explain why PC2_BrainSpan_ was less with PC2_Allen_ (11 regions, less granular) compared to PC2_Kang et al._ (22 regions, more granular).

In summary, just two components explain the majority of gene expression variation across the human cerebral cortex. Unrotated, PC1 accounted for 31.9% of the variance, and PC2 for 17.5%, (after varimax rotation PC1 accounts for 25.8% of the variance, and PC2 for 23.6%). These two components are not heavily reliant on individual regions, nor are they donor-specific (see Figure 2). The first component is robust across all validation tests but, for the second, we note some effects of cortical boundary positioning, sampling differences, and the number of retained genes – factors which can partly be attributed to technical confounds. With these results in mind, we extracted two components, which together account for 49.4% of the variance, with varimax rotation for further analysis.

### Interpretation of gene expression components

To aid interpretation of the two components, we conducted statistical overrepresentation analyses, at http://geneontology.org/, which is powered by PANTHER (^62^). The results suggest that Component 1 represents cell-signalling and post-translational modification processes (with loadings < −0.3 providing upregulation and those > 0.3 providing downregulation) (see *Figure 3* and the supplementary data file for full GO results and component loadings). Prominent GO terms include i) amino acids and organic compounds, which provide energy to cells and hasten chemical reactions necessary for post-translational modifications, and ii) signalling terms, which convey information about nutrients in the environment and support coordination between cells. Component 2 is a transcription factors axis (with loadings < −0.3 providing downregulation and those > 0.3 providing upregulation). The GO terms implicate biosynthesis, binding and RNA polymerase II, defining characteristics of transcription factors.

**Figure 3.**
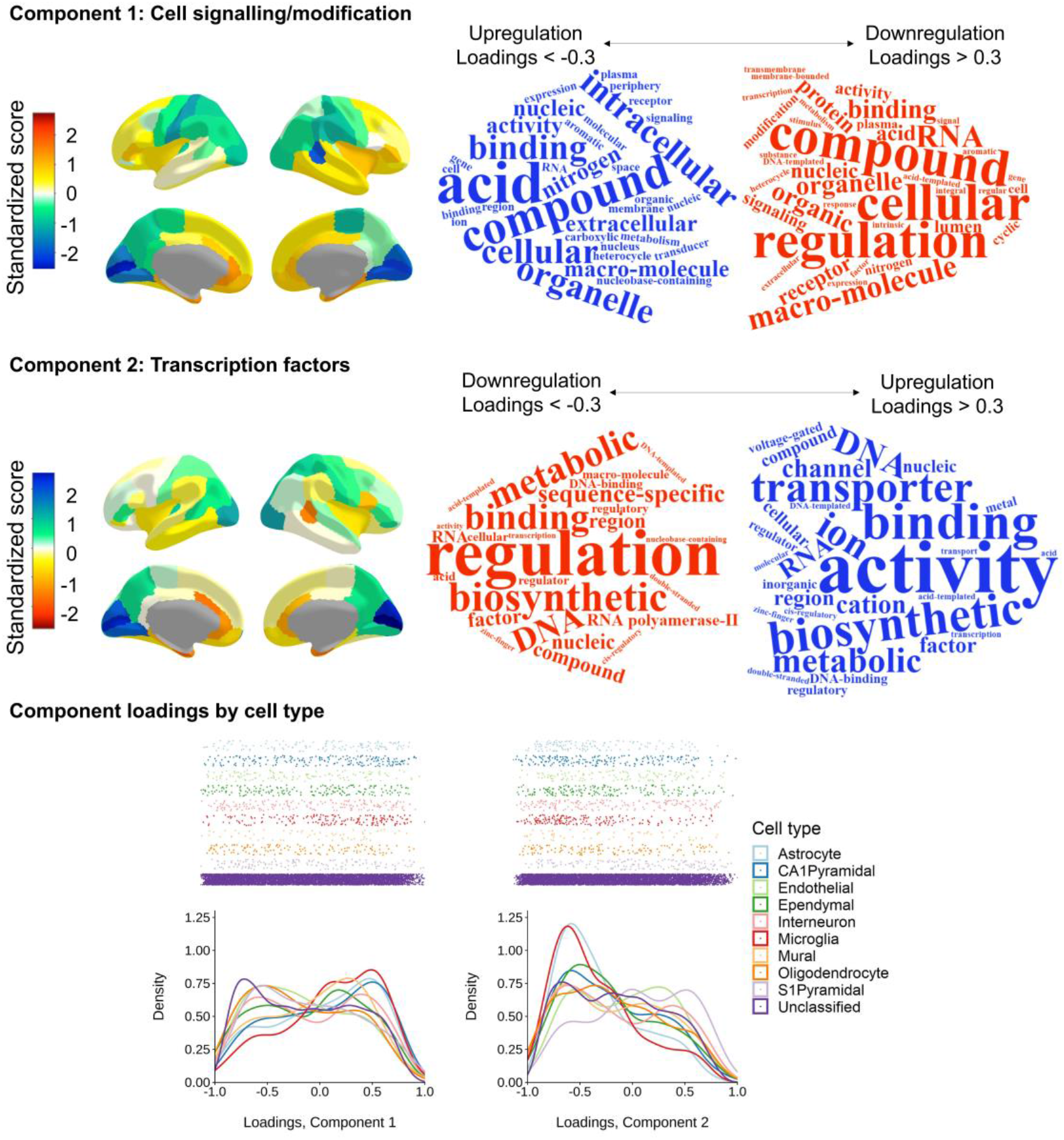
Two major components of cortical gene expression. *Figure 3 note* Top and middle panels (Component 1 and Component 2, respectively) left: Regional *z* scores mapped to the cerebral cortex (scaled for each hemisphere) and right: word clouds of the statistical over-representation results. The relative direction of component scores is arbitrary (dictated by the PCA), and here, the colour scale is flipped between components so that the directions of upregulation/downregulation match. Bottom panel: Density distribution plots of loadings on Component 1 and Component 2 coloured by cell type.

Additionally, we tested whether the distribution of component loadings differed by cell-type. Zeisel et al. (^63^) identified proteins expressed in 9 specific cell-types, from single-cell transcriptomes of 3005 cells in the mouse somatosensory cortex and hippocampus. Shin et al. (^64^) converted these genes to human gene symbols, with the HologoGene database (^65^). We matched these genes with those available in our current dataset (*N =* 8235 genes). Out of the initial set of cell-specific genes, in our dataset there were 129/214 astrocytes, 204/357 CA1 pyramidal neurons, 127/321 endothelial cells, 191/415 ependymal cells, 181/293 interneurons, 185/374 microglia, 60/133 mural cells, 139/393 oligodendrocytes, 155/236 S1 pyramidal neurons, and the remaining 6864 proteins were “unclassified”, and treated as a baseline group.

Descriptive statistics of component loadings for each cell-type are in *Table S3* and the results of the Dunn posthoc tests are in *Tables S4 and S5.* Generally, the loadings of different cell types tend to be skewed. For Component 1, loadings for all but ependymal and interneuron cell types have an absolute skewness value > 0.228. For Component 2, all but endothelial cells have absolute skewness > 0.187. This skewness in loadings suggests that specific cell types might play a particular roles in the regulation of the two components. We investigated whether specific cell types load on the two major components in ways that deviate from the average distribution of “unclassified” loadings. Loading distributions by cell type are shown in *Figure 3* (bottom panel) and descriptive statistics and full results of Dunn posfthoc tests, with *p*-values adjusted with the Holm method, are in *Tables S3-5*. There are main effects of cell classification for both components (Component 1: *H*(9) = 88.986, *p =* 2.6e-15, Component 2: *H*(9) = 81.046, *p =* 1.001e-13).

For Component 1, the unclassified set’s distribution tends towards the expression side of the axis (*M =* 0.14, *SD =* 0.49, skewness = −0.145). This contrasts with astrocytes (*z =* −4.05, *p =* .002; *M =* −0.04, *SD =* 0.49, skewness = 0.375), CA1 pyramidal neurons (*z =* 05.12, *p =* 1.36-05; *M =* −0.03, *SD =* 0.50, skewness = 0.254) and microglia (*z =* −5.60, *p =* 1.86e-09; *M =* −0.11, *SD =* 0.46, skewness = 0.506), which are skewed towards the regulation side. For the second component, the unclassified set of genes tend towards regulating transcription factors (*M =* 0.15, *SD =* 0.46, skewness = −0.183). Whereas, S1 pyramidal cells oppose this direction (*z =* −5.03, *p =* 1.98e-08; *M =* −0.05, *SD =* 0.47, skewness = 0.200), astrocytes and microglia fall more sharply on the regulation side than the unclassified set of genes (unclassified kurtosis: 1.925; astrocytes: *z =* 3.89, *p =* .003, kurtosis = 2.582; microglia: *z =* 5.41, *p =* 2.66e-06, kurtosis = 2.846). For all other comparisons between the unclassified cells and individual cell types, *p =* 1.

Additionally, through FUMA https://fuma.ctglab.nl/ we tested whether the genes with absolute loadings > 0.3 on each component were significantly related to gene sets in the GWAS catalog. There were no significant (α = .05) associations for either component, demonstrating the highly general nature of the two components of cortical gene expression.

### Regional distribution of component scores across the cerebral cortex

The component scores were scaled to correct for interhemispheric differences in gene expression values (see Methods for details). These component scores were then mapped to the 68 Desikan Killany regions (see *Figure 4* and *Table* S9). There was a strong interhemispheric correlation in scores between the 34 paired regions for both Component 1 (*r =* 0.815, *p =* 4.411e-09) and Component 2 (*r =* 0.725, *p =* 1.25e-06). The direction of upregulation and downregulation of the loadings (as determined by GO analysis) informed whether the regional component scores suggested upregulation or downregulation of the two components. For Component 1, negative loadings suggest upregulation, and positive suggest downregulation; conversely for Component 2, positive loadings suggest upregulation and negative suggest downregulation. Parietal and occipital regions are on the upregulation side of Component 1 (cell signalling/modification), with frontal and temporal regions indicating downregulation. For Component 2 (transcription factors), lateral frontal areas tend towards balance between upregulation and downregulation, whereas medial frontal regions tend towards downregulation and parietal and occipital regions towards upregulation. For both components and in both hemispheres, the highest absolute scores are observed in the medial occipital regions (pericalcarine, cuneus and lingual), which fall strongly on the upregulation side of both components.

**Figure 4.**
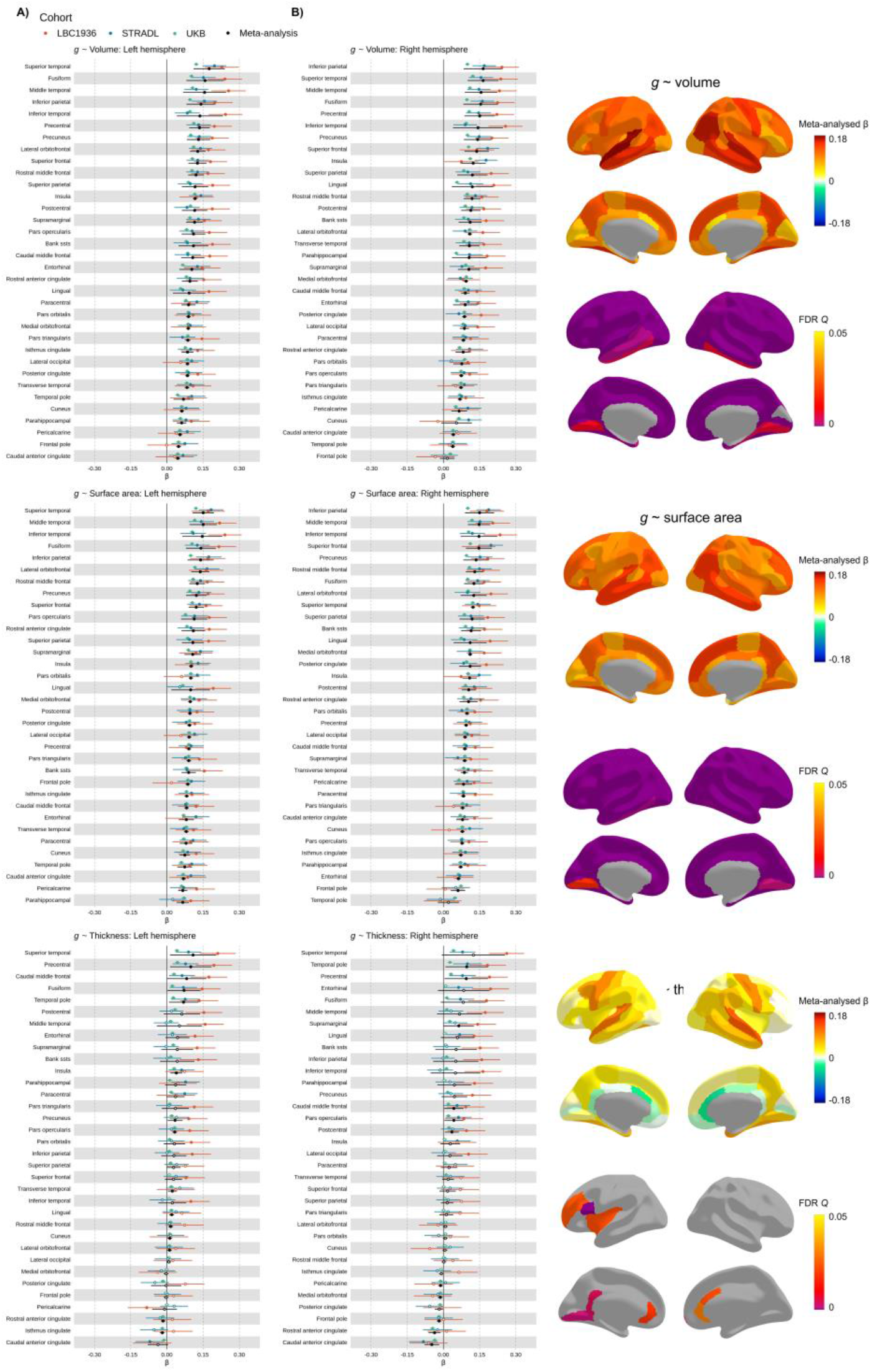
Meta-analysed brain regional associations with *g.* *Figure 4 note* A) Standardised β estimates mapped to the cerebral cortex. B) Meta-analysed standardised β estimates for *g-*volume, *g-*surface area and *g-*thickness. Those for which *p* < .05 are filled in, and those for which *p* > .05 are outlined. The y-axis is ordered by the meta-analysed β values (decreasing).

We tested whether the regional mean morphometry profiles (*see Supplementary Text 1*) are associated with regional gene expression component score patterning. In general, the thicker a region is, the more strongly it falls on the downregulation side of both gene expression components (Component 1: *r =* 0.764, *p =* 3.67e-14 and Component 2: *r =* - 0.799, *p =* 3.132e-16). Associations between mean regional surface area patterns and both components were small-to-moderate (Component 1: *r =* −0.230, *p* = .059, Component 2: *r =* 0.245, *p* = .044), and between mean regional volume and both components were small and not statistically significant at the α < .05 level (Component 1: *r =* −0.082, *p* = .504, Component 2: *r =* 0.111, *p* = .368).

### Cortical morphometric associations with general cognitive functioning *(g)* – Meta-analyses (*N =* 39,519)

We first used raw data from three cohorts to estimate regional associations between three MRI-derived morphometry measures (cortical volume, surface area and thickness) and *g* (total *N =* 39,519; three cohorts – the UK Biobank (UKB, _6_^66^, http://www.ukbiobank.ac.uk): *N =* 37,840 participants (53% female), age *M =* 63.81 years (*SD =* 7.64 years), range = 44-83 years; STRADL (^67^, a Generation Scotland imaging sample): *N =* 1043 participants (60% female), age *M =* 59.29 years (*SD =* 10.12 years), range = 26-84 years; and the Lothian Birth Cohort 1936 (LBC1936, ^68, 69^ https://www.ed.ac.uk/lothian-birth-cohorts): *N =* 636 participants, (47% female), age *M =* 72.67 years, *SD =* 0.41 years, range = 70–74 years). General cognitive function (*g*) scores were derived using confirmatory factor analysis (in a structural equation modelling framework) in each of the three cohorts using multi-domain cognitive test batteries, and each individual test score was corrected for age and sex. As one of the most replicated phenomena in psychological science*, g* is based upon the tendency for performance on all cognitive tests to be correlated, and is generally invariant to cognitive test content, provided that multiple domains are captured (^70^). These properties lend it well to cross-cohort genetic analyses (^23^), for example, and we leverage them here.

Latent *g* scores were extracted for all participants, and associations with three measures of cortical morphometry (volume, surface area and thickness) were estimated for each of the 68 regions in each cohort. Cortical measures were controlled by age, sex, head position in the scanner (X, Y and Z coordinates), testing site (for UKB and STRADL) and lag between cognitive and MRI testing appointments (for LBC1936). For UKB, X, Y and Z co-ordinates were calculated relative to one target participant, and for LBC1936 and STRADL, they were taken from the mri_info -cras flag output. We computed standardised β estimates of the association in each brain region between *g* and each brain morphometric property (volume, surface area, thickness) for each cohort. There were strong cross-cohort correlations for *g-*associations between the 68 regions for each measure of morphometry (see *Table 1*).

**Table 1.**
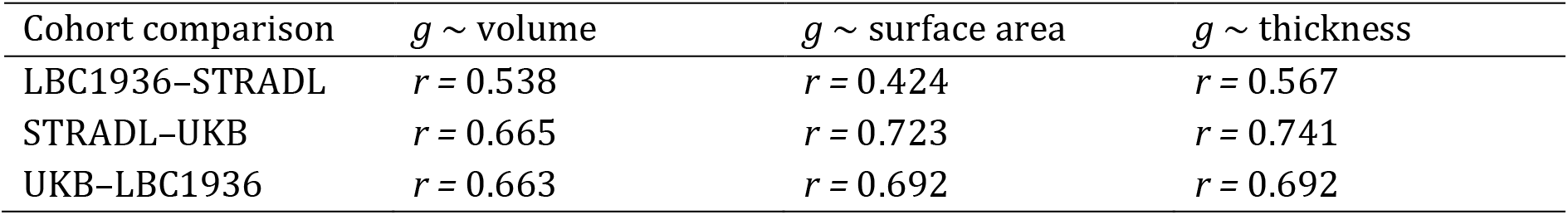
Cross-cohort correlations of regional *g-*associations. *Table 1 note* (all *p <* .001).

| Cohort comparison | $g \sim$ volume | $g \sim$ surface area | $g \sim$ thickness |
| --- | --- | --- | --- |
| LBC1936–STRADL | $r = 0.538$ | $r = 0.424$ | $r = 0.567$ |
| STRADL–UKB | $r = 0.665$ | $r = 0.723$ | $r = 0.741$ |
| UKB–LBC1936 | $r = 0.663$ | $r = 0.692$ | $r = 0.692$ |

We then ran a random effects meta-analysis on the standardized β values. The meta-analytic results of the three cohorts’ associations between *g* and brain morphometry data (68 regions x 3 measures = 204 meta-analyses) are summarised in *Figure 4* and reported in detail in *Tables S19-S21*. Meta-analysed standardised βs for *g*-volume associations *M* β = 0.103 (SD = 0.034, range from 0.015 to 0.175), for *g-*surface area, *M* β = 0.102 (SD = 0.027, range from 0.020 to 0.150), and for *g*-thickness associations *M* β = 0.031 (SD = 0.035, range = −0.048 to 0.124).

The current results provide support for theories (^25,26^) regarding which regions are key in brain morphometry-*g* associations (e.g. the parieto-frontal integration theory, P-FIT, see Figure S15). Parietal and frontal regions generally have relatively strong *g-* associations with volume and surface area, though not with cortical thickness. For all three morphometry measures, the superior temporal region had relatively high *g-* associations mean β (between hemispheres) = 0.163, 0.143 and 0.116, for volume, surface area and thickness respectively. Some of the highest *g-*volume and *g-*surface area associations are for the fusiform (mean β (between hemispheres) = 0.154 and 0.126, for volume and surface area, respectively) and inferior parietal region(mean β (between hemispheres) = 0.153 and 0.145, respectively). The precuneus regions also have among the overall highest associations (mean β = 0.136, and β = 0.129), in line with updated reports of regional *g-*associations (^71, 26^).

There is high inter-hemispheric consistency for each of the meta-analytic *g-*morphometry associations: volume (*r =* 0.887, *p =* 2.988e-12), surface area (*r =* 0.807, *p =* 8.105e-09), and thickness (*r =* 0.878, *p =* 9.578e-12), see *Figure S14*. In addition, estimates were strongly correlated between *g-*volume and *g-*surface area associations (*r =* 0.831, *p* = 1.66e-18), and moderately correlated between *g-*volume and *g-*thickness (*r =* 0.579, *p =* 2.365e-07). As anticipated, based on previous work showing phenotypic and genetic distinctions between surface area and thickness (^28,29,30,31^), the correlation between *g-* surface area and *g-*thickness estimates was small and not statistically significant at the α *< .*05 level (*r =* 0.150, *p =* .222).

To help interpret why some regions might have higher associations with *g* than others, we tested correlations between regional *g-*associations and the regional mean profiles of volume, surface area and thickness (reported in *Supplementary Text 1*). The regional *g-* associations were positively associated with the corresponding regional mean profiles for all three morphometry measures. Volume had the strongest correlation (*r =* 0.709, *p* 1.35e-11), followed by surface area (*r =* 0.614, *p =* 2.58e-08), and then thickness (*r =* 0.313, *p =* .009). In other words, regions with stronger *g-*associations tend to be larger in terms of volume and surface area, and also moderately tend to be thicker.

We then tested whether larger and thicker brain regions are more strongly associated with *g* because they tend to be better proxies for whole-brain measures (as they contribute more to the total measure). The maginitude of total brain *g*-volume and *g-* surface area associations are in line with the maximum of the individual regions – *g-* whole cortex volume (β = 0.180, SE = 0.036, p = 5.93e-07), *g-*whole cortex surface area (β = 0.160, SE = 0.021, p 3.93e-14). Perhaps as there are some negative *g-*thickness associations, the *g-*whole cortex mean thickness associaiton was not significant (β = 0.065, SE = 0.043, p = 0.13). We then corrected regional *g-*associations for these total brain measures, and correlated regional profiles with and without correction. These correlations are moderate-to-strong: *g-*volume (*r =* 0.613, p = 2.76e-08), *g-*surface area (*r =* 0.556, *p* = 8.78e-07), and *g-*thickness (*r =* 0.945, *p* < 2.2e-16), suggesting that it is not simply because larger/thicker brain regions are a better proxy for the whole brain measure that they are more strongly associated with *g*.

### Interregional variation in gene expression corresponds to interregional variation in cognitive function

Next, we tested whether brain regions’ differences in gene expression (as measured using the two components we described earlier) are correlated with *g-*morphometry associations. That is, we asked whether brain regions for which morphometric measures (volume, surface area and thickness) are more strongly related to *g* were also more strongly related to general dimensions of gene expression.

We tested linear correlations between the absolute component scores of gene expression and the meta-analysed standardised β scores for cortical morphometry associations with *g*, and also report the comparable quadratic regression results with non-absolute scores (see *Table 2* and *Figure 5*). There were negative associations for all analyses, and those for *g-*volume and *g-*surface area were moderate-to-strong and statistically significant at the α < .05 level. These results suggest that, generally, regions more strongly associated with *g* tend to be more balanced between the downregulation and upregulation sides of both cell-signalling/modification and transcription factors components.

**Figure 5.**
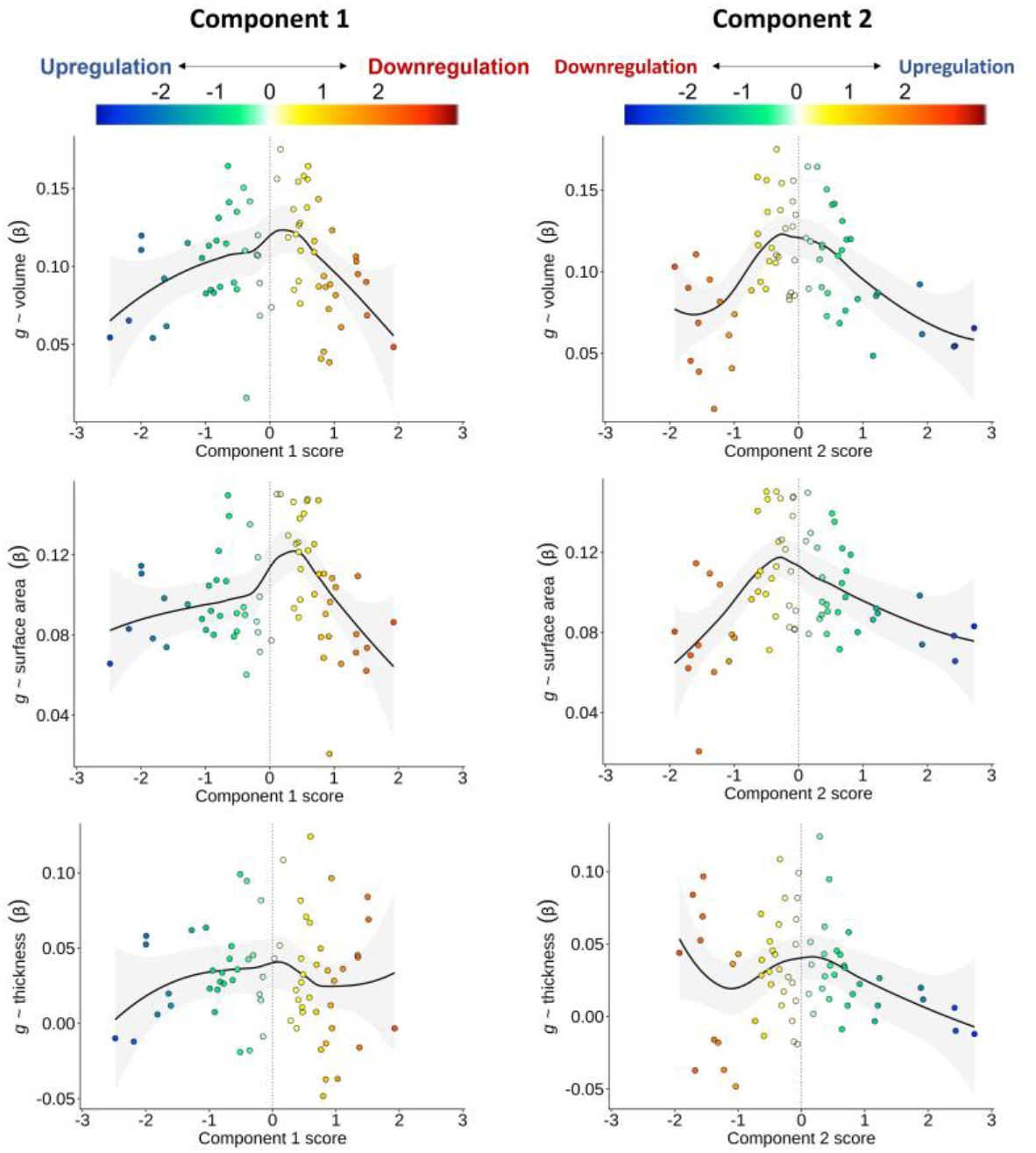
Associations between regional-*g* profiles and the two gene expression components. *Figure 5 note* LOESS functions are plotted (the quadratic model results are comparable to the absolute score correlations and are presented in *Table S25*). A vertical line at component scores of 0 represents a balance between upregulation and downregulation ends of each component. The colour scale is flipped between Components, so that the direction of downregulation and upregulation match.

**Table 2.** Correlations of correlations: Meta-analysed *g*-cortex associations with two major gene expression components. *Table 2 note.* Pearson’s *r* values for the correlation between brain-g associations and brain-gene expression component profiles. Note that these are for linear associations using the absolute gene-expression component scores. Results for the equivalent associations using the non-absolute components scores (quadratic component) are presented in *Table S25* and *Figure 5*, which illustrates the balance between downregulation and upregulation.

| *g* ~ | Component 1 | Component 2 |
| --- | --- | --- |
| Volume | $r = -0.385, p = .001$ | $r = -0.582, p = 1.96\text{e-}07$ |
| Surface area | $r = -0.345, p = .004$ | $r = -0.500, p = 1.45\text{e-}05$ |
| Thickness | $r = -0.145, p = .237$ | $r = -0.255, p = .036$ |

There were no correlations between regional mean expression across genes and *g-*brain morphometry association profiles for which *p <* .05 (*g-*volume *r =* −0.023, *p =* .853; *g-* surface area *r =* −0.058, *p =* .640; *g-*thickness *r =* 0.103, *p =* .403), demonstrating the value of the PCA approach as associations between genome-wide dimensions of expression and *g* are not reducible to an average brain-wide pattern of gene expression.

### Associations between *g* and individual genes

Lastly, we tested for individual gene-*g* expression pattern associations, controlling for the two general components of gene expression. For all 8235 individual genes, the median expression scores per region were scaled separately for left and right hemisphere regions, to account for sample-based artifacts in hemisphere differences in expression values, in line with the method for the component scores. After FDR correction (threshold = *Q* < .05), there were 522 individual genes whose cortical patterning was correlated with *g-*volume patterning, 609 with *g-*surface area and 516 with *g-*thickness (these results are available in detail in the supplementary data file, and Figures S18-S20). 268 genes were shared between *g-*volume and *g-*surface area, 253 between *g-*volume and *g-*thickness and 42 between *g-*surface area and *g-*thickness. 41 genes with *FDR Q* < .05 overlapped for all three morphometry measures (|β| range = 0.15 to 0.53, see *Figure 6*). These genes are particularly likely candidate substrates of cognition. Some regional expression profiles have positive associations with *g*-cortical measure profiles, while others have negative associations. For discussion, genes with negative *g-*associations in the present study are marked with an asterisk. We also ran a protein network analysis through STRING^72^ (Search Tool for the Retrieval of Interacting Genes/Proteins), with a minimum required interaction score of “medium confidence” (0.400), and the background set of 8235 genes. Four proteins were not available in STRING, so there were 37 proteins involved in the analysis (out of a possible total of 41). The results are shown in Figure 6b. The PPI enrichment *p-*value is .909, with the expected number of edges being 4, and the observed number of edges, 2, showing that the network does not have significantly more interactions than expected. The only two edges that meet the threshold are between GABRQ and GABARAP, and between MYL3 and SMPX. These results suggest that there are not widespread interactions between these top genes, which further validates the aim of establishing unique signals beyond general gene expression patterns.

**Figure 6.**
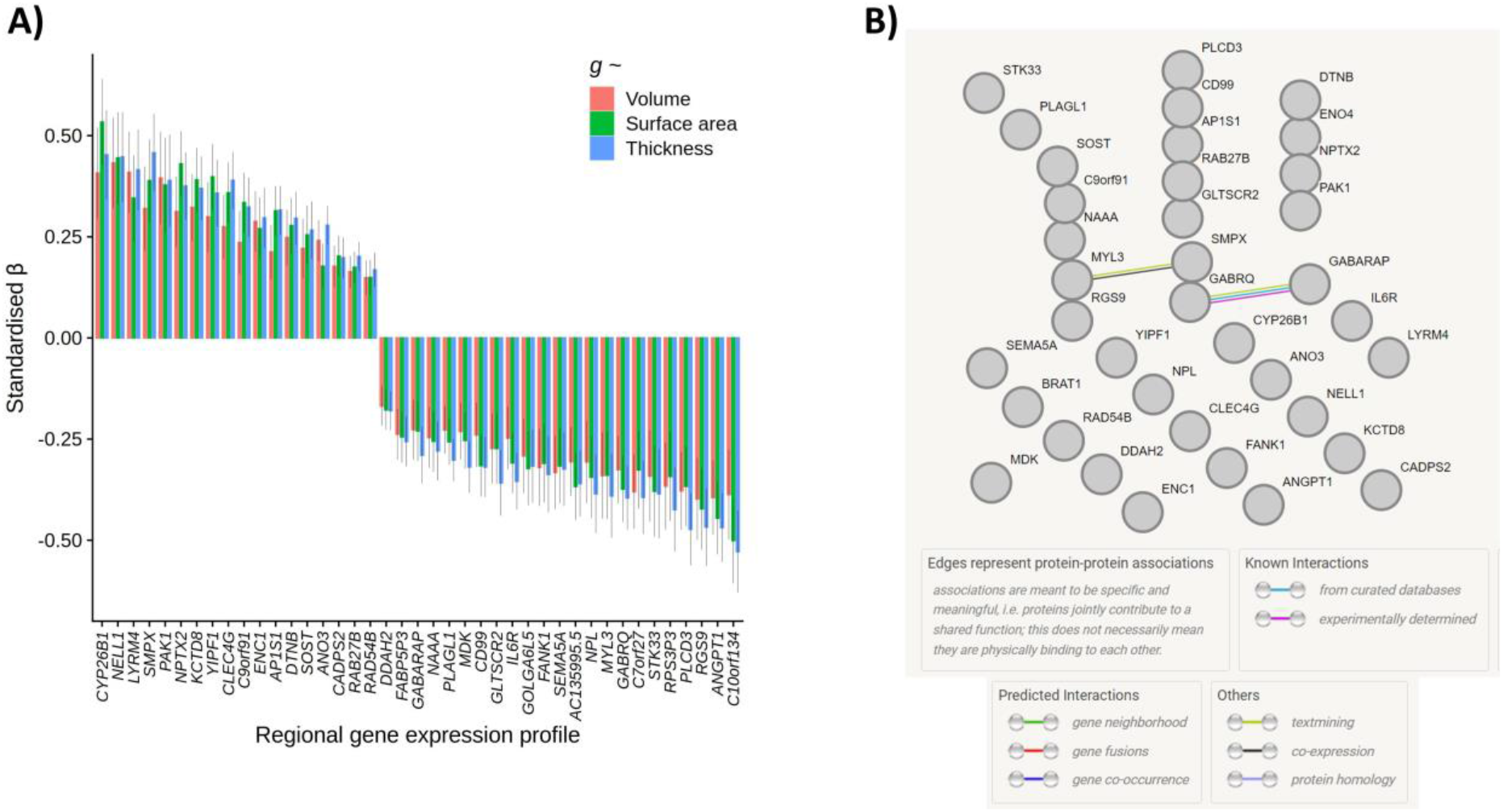
41 specific associations between regional g-morphometry profiles and individual gene expression profiles. A) β values for volume, surface area and thickness regional associations with the individual gene expression profiles. B) STRING results, showing paths of evidence-based interactions between them. *Note:* only 37 out of 41 proteins were available in STRING. *Figure 6 Note* Standardised β for specific individual gene profiles (i.e. corrected for general components of gene expression) for which FDR Q > .05 for all three cortical morphometry associations with g.

Several of these 41 genes have been previously reported to be associated with Alzheimer’s Disease: overexpression of *ANGPT1*\* has been found to increase amyloid beta secretion (^73^) whilst *CLEC4G* suppresses amyloid beta (^74^), and increased levels of *ENC1* (^75, 76^) and *NPTX2* (^77, 78, 79, 80^) are consistently demonstrated to have protective effects against cognitive decline in Alzheimer’s disease. Other genes from this list have been associated with other neurodegenerative disorders – for example. Loss of *GABRQ*\*-containing neurons is an indicator of early social-emotional cognitive decline in frontotemporal dementia (^81^), IL6R* has been associated with memory domain scores, and Alzheimer’s disease pathology (cerebrospinal fluid pTau and Aβ42/40 ratio) (^82^), *RAB27B* regulates α-Synuclein, which is a primary indicator of Parkinson’s disease and dementia with Lewy bodies (^83^), and *DTNB* is an indicator of extent of neuronal injury and inflammation in Alzheimer’s disease (^84^). Additionally, *SEMA5A*\* has previously been associated with hippocampal volume and performance on cognitive tests (^85^). *CYP26B1* is upregulated in the prefrontal cortex (^86^), and there are links between its catabolism in the hippocampus and poor cognitive outcomes in mice (^87^).

Other individual genes associated with *g-*cortical profiles have been associated with cognitive functioning more generally, for example, *RGS9*\* has been implicated in motor coordination and working memory (^88^), *CADPS2* is associated with cognitive functioning and memory in healthy adults (^89^), and *FANK1\** has been found to have genome-wide significant associations with *g* in CHARGE-COGENT and UKB cohorts (^23^). Some others have been linked to cognitive disorders, for example *LYRM4* with schizophrenia (^90^), and *DDAH2*\* with multiple neurological conditions and psychiatric disorders (^91^). Previous significant GWAS associations with these 41 genes were identified in the GWAS catalog and are available in the supplementary data file. Recurrent associations include educational attainment, body mass index (BMI), brain measurements, coronary artery disease, schizophrenia, and depression (see the supplementary data file). Potential novel, or less-studied, individual gene substrates of complex cognitive processing identified in the present study include: *AC135995.5*\**, ANO3*, *AP1s1, C7orf27*\*, *C9orf91, CD99*, FABP5P3*\*, GABARAP*, *GLTSCR2\**, *GOLGA6L5*, KCTD8*, *MYL3*, NAAA*, NPL*, PAK1*, *PLAGL1*\*, *PLCD3*\**, RPS3P3*, SMPX, SOST, STK33\** and *YIPF1*.

### Associations between *g* and cell types

Then, we used our discovery of what is common about regional cortical gene expression profiles to identify specific cell type-cognitive relationships. The mean profiles of the 9 specific cell types were scaled in each hemisphere, and we controlled for the two major components of gene expression (detailed regression results are in *Supplementary Table S26*). For two cell types, there were FDR Q < .05: ependymal cells with *g-*volume (β = - 0.200, SE = 0.054, FDR Q = .007) and with *g-*thickness (β = −0.244, SE = 0.053, FDR Q = .001), and for microglia, with volume (β =-0.155, SE = 0.054, FDR Q = .035) and surface area (β = −0.175, SE = 0.053, FDR Q = .013).

## Discussion

This study reveals and validates a fundamental organisation principle of cortical gene expression patterns across the human brain. We then use this information to identify the shared and specific aspects of regional cortical gene expression and show that they are associated with regional brain-structure correlates of complex thinking skills. We also show that this information is not obtainable by simply considering aggregate/mean levels of gene expression across regions.

We validated our discovery of two major components of interregional variation in gene expression: one indicating cell-signalling/modifications and the other, transcription factors. Using the largest meta-analysis of the cortical loci of general cognitive functioning (*g*) to-date, we find that regions that are more balanced between downregulation and upregulation of these two gene expression components are most strongly associated with *g.* Controlling for these established patterns of gene expression covariation allowed us to identify which individual genes had spatial expression patterns that specifically reflect cortical correlates of *g*, beyond the major dimensions of gene expression. Critically, without this approach, one is likely to miss or erroneously ascribe an interpretation to an individual gene, as its profile is confounded by major components of shared spatial covariation across multiple gene expression patterns.

We conducted one of the largest analyses of *g*-cortical morphometry associations to-date. These associations are generally in line with the parieto-frontal integration theory (P-FIT,^25^) and strengthens support for the involvement of regions (e.g., temporal, precuneus) that were not included in earlier iterations of the model. There was strong agreement across the three cohorts in the magnitudes and spatial patterning of associations, which speaks to the validity of *g* as a measure of cognitive functioning (indicated by different cognitive tests included by each cohort). The consistency of results also indicates that careful harmonisation of image processing alongside careful attention to phenotype measurement may partially offset the apparent need for many thousands of participants to obtain replicable brain-behaviour association results (^27^).

Turning to the gene expression components-*g* correlations, the more strongly regions were associated with *g*, the more they tended towards the balance between the downregulation and upregulation sides of the cell-signalling/modification and transcription factors components. Complex cognitive functions therefore may be facilitated at a midpoint of downregulation and upregulation of each of these components. Other regions that fall on either the downregulation or upregulation sides of each of the two major dimensions perhaps specialise in less general functions. An important question this raises, but we cannot answer here, is whether individual differences in the balance of gene expression in these cortical dimensions might partly explain why people differ from each other in their general cognitive functioning.

Using this newly-gleaned information about what is common among gene expression profiles, we then identified 41 individual genes whose spatial expression was correlated with *g* cortical patterning, independent of the two major dimensions. Whereas some of the genes strongly indicated in the two major dimensions themselves could also pertain, causally, to mechanisms and processes underpinning *g,* the nature of shared expression patterns as presented here disallows that direct inference for individual genes. In contrast, these 41 genes with specific associations are particularly strong candidates for playing a role in facilitating complex cognitive processes. Several of these genes have been previously identified as associated with various cognitive outcomes, whilst others are potentially less well-researched substrates of cognition.

After testing the g-associations of the major components of gene expression, we turned to specific cell type–g associations. Microglia and ependymal cells both had negative associations with cognitive morphometry measures - ependymal cells with volume and thickness, and microglia with volume and surface area. These two cell types both play key roles in waste removal from the brain, which might explain these negative associations – it could be that some regions specialise in fundamental brain maintenance processes, such as waste management, thus enabling others to specialise in cognitive processes.

This study has several strengths and limitations. As we quantitatively demonstrate, the present approach surpasses candidate gene and median expression information, clarifying our understanding of the molecular substrates of complex cognitive abilities in the human brain. We extensively validate the discovered components of gene expression, mitigating concerns that this finding might be an artefact of a small number of donors in a single sample. The first component is highly consistent across different datasets, gene expression data processing and summary choices, and brain regional parcellation choices. Although the second component does not depend heavily on individual regions, it is somewhat affected by the granularity and boundaries of the parcellation, gene expression data sampling and processing choices, and the number of genes retained. While efforts continue to standardize gene expression processing pipelines (^92^), the effects of different choices on dimensions of between-gene covariances should continue to be considered. As donor contributions to gene expression databases continue to increase, brain regional summaries of gene expression will become more precise. Several genes were excluded from the current analysis due to low between-donor consistency. Although this is partially due to some of these genes having generally low expression across the cortex, there also appears to be an effect of the sampling methods of gene expression data. Gene expression sampling methods consistent with clear cortical boundaries and full cortical coverage will increase between-donor consistency in regional gene expression profiles and enable stronger tests of external validity. Additionally, future research should consider whether major dimensions of regional cortical gene expression, such as those reported in the current paper, are consistent between postmortem and in vivo data (^93^).

We leveraged the fact that the UKB, STRADL and LBC1936 cohorts have adopted comparable methods, including similar MRI processing pipelines with FreeSurfer http://surfer.nmr.mgh.harvard.edu/, and collection of various cognitive test scores, which enabled us to harmonise the processing and approach to the calculation of *g.* Consistency in the applied methods between cohorts allows for direct quantitative comparison. Despite these advantages, there were also some differences between MRI data and processing the three cohorts, which might differentially affect the cortical surface results: 1) each of the three cohorts used different scanners for MRI acquisition and, although T1-weighted data provides consistent between-scanner measures (^94^), we cannot rule out scanner-specific differences, 2) Desikan-Killiany parcellations were visually inspected and manually edited for LBC1936 and STRADL, but not for UKB (outliers *SD* > 4 were excluded), and 3) different FreeSurfer versions were used for each cohort, which is likely to have contributed to some differences in estimations, alongside different types and quantity of cognitive tests. However, high between-cohort correlations suggest that these differences may not meaningfully affect the current results and provide evidence in support the use of *g* in meta-analytic studies to reach reproducible brain-cognition associations (^95^).

A separate limitation of this study is that all included participants were in relatively good health, as we chose to exclude participants with declared neurological conditions. It is therefore not clear that the reported regional *g-*associations would generalise to clinical populations. Additionally, whereas the cognitive-MRI data do not include childhood and adolescence (and therefore the results may not relate directly to those parts of the life span), the good adulthood age coverage, absence of age moderation of the meta-analytic estimates within-cohort, and clear agreement across cohorts suggests that the well-powered results reliably capture adulthood brain-*g* correlations.

In summary, this newly possible study uses robust methods to advance our understanding of how gene expression is associated with complex cognitive functioning. We discovered and interpreted two general components of cortical gene expression, and identified general and specific patterns of gene expression that are candidate substrates that may contribute to some of the association between brain structure and complex cognitive functioning.

## Supporting information

Supplementary tabular data

## Acknowledgements

We thank the participants of the three cohorts (UKB, Generation Scotland (STRADL) and LBC1936) for their participation and the research teams for their work in collecting, processing and giving access to these data for analysis. We are also thankful the brain donors to the Allen Human Brain Atlas, BrainSpan Atlas and Human Brain Transcriptome Project, and to the people who collected and processed the data and made it openly available.

SRC and JEM were supported by a Sir Henry Dale Fellowship, jointly funded by the Wellcome Trust and the Royal Society [221890/Z/20/Z]. For the purpose of open access, the author has applied a CC-BY public copyright licence to any Author Accepted Manuscript version arising from this submission. The LBC1936, supported by the BBSRC & ESRC [BB/W008793/1], Age UK [Disconnected Mind project], the Medical Research Council (MR/M01311/1; MR/K026992/1), the US National Institutes of Health [R01AG054628] and the University of Edinburgh. CRB, MEB, EMT-D, IJD and SRC were supported by a National Institutes of Health (NIH) research grant R01AG054628. TCR is a member of the Alzheimer Scotland Dementia Research Centre funded by Alzheimer Scotland. MCVH is funded by The Row Fogo Charitable Trust Centre for Research into Aging and the Brain (BRO-D.FID3668413).AS was funded as part of the STRADL study (Wellcome Trust reference 104036/Z/14/Z) and indirectly through the Lister Institute of Preventive Medicine research award for Prof. Daniel Smith (ref. 173096).

