## Supplementary figures, tables and text for "General and specific patterns of cortical gene expression as spatial correlates of complex cognitive functioning"

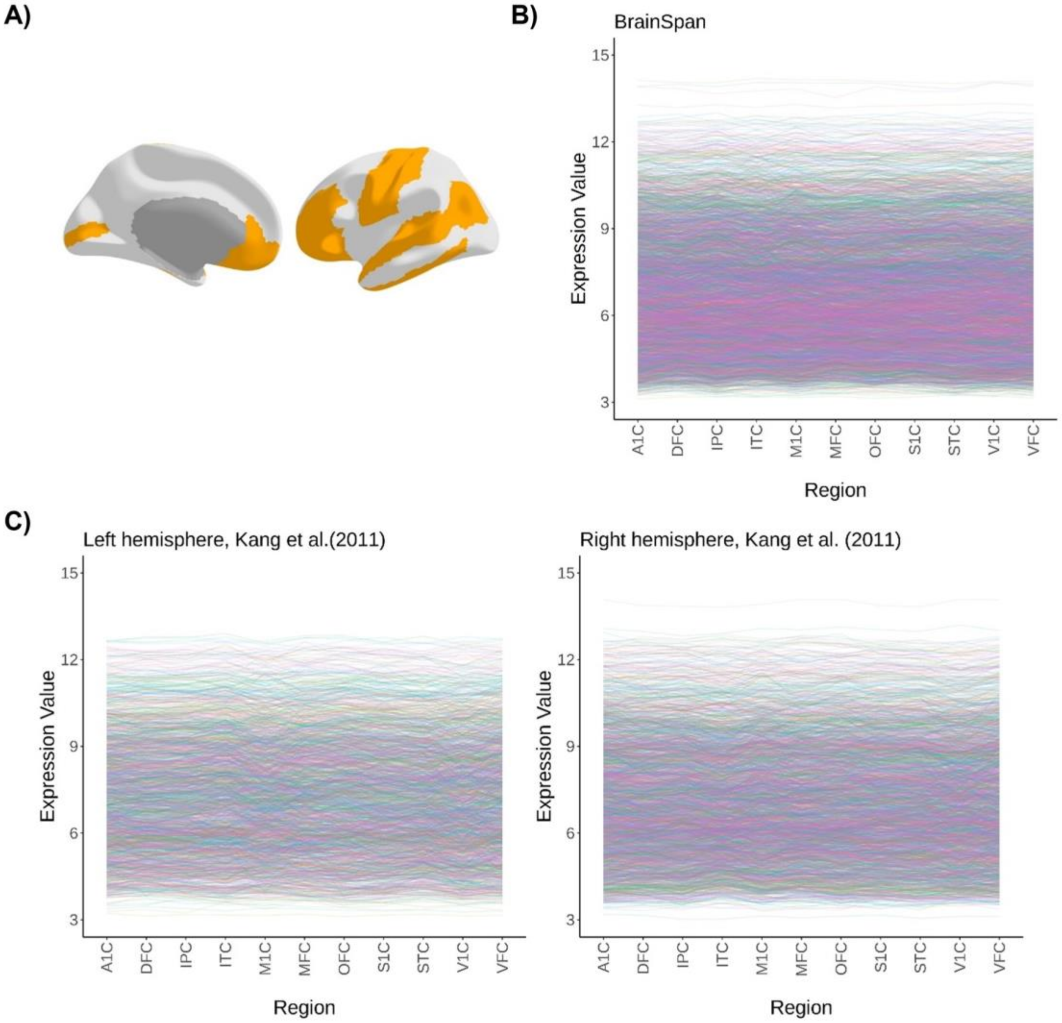


*Figure S1* Raw gene expression values for the two validation datasets, for genes that have between-donor consistency and are matched with the 8235 consistent genes in the Allen dataset. A) The 11 cortical regions of the two validation datasets roughly matched to 14 Desikan-Killiany regions, as suggested by Wong et al.[1] B) Data from the Brainspan dataset. Bilateral regions, 4410 consistent and matched genes, and C) Gene expression data from Kang et al. (2011) for the left hemisphere, 2030 consistent and matched genes and for the right hemisphere, 2778 genes.

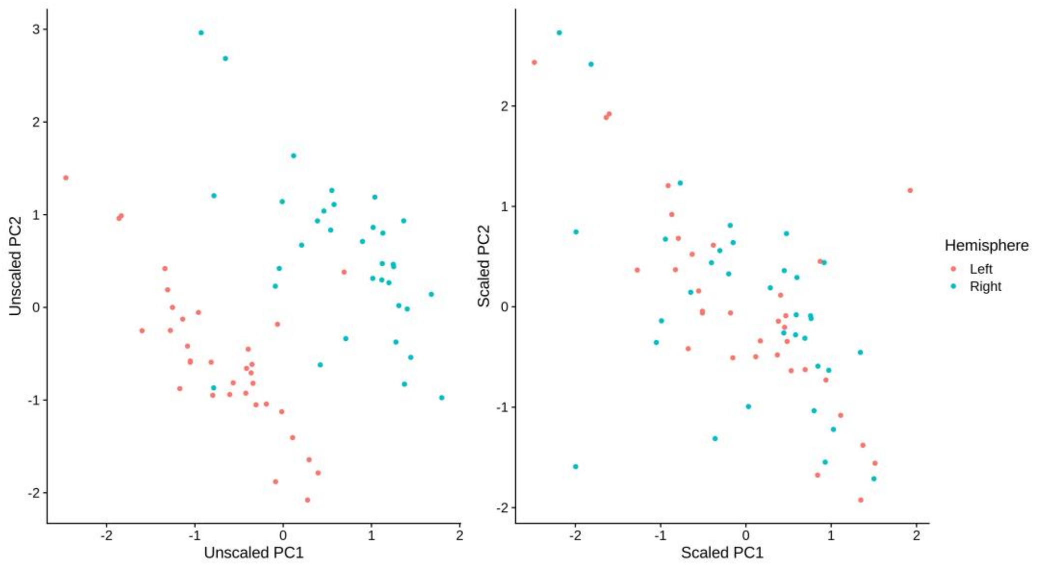


*Figure S2* Principal component scores for Component 1 and Component 2, after varimax rotation. Left: before scaling the scores in each hemisphere; Right: after scaling each hemisphere.

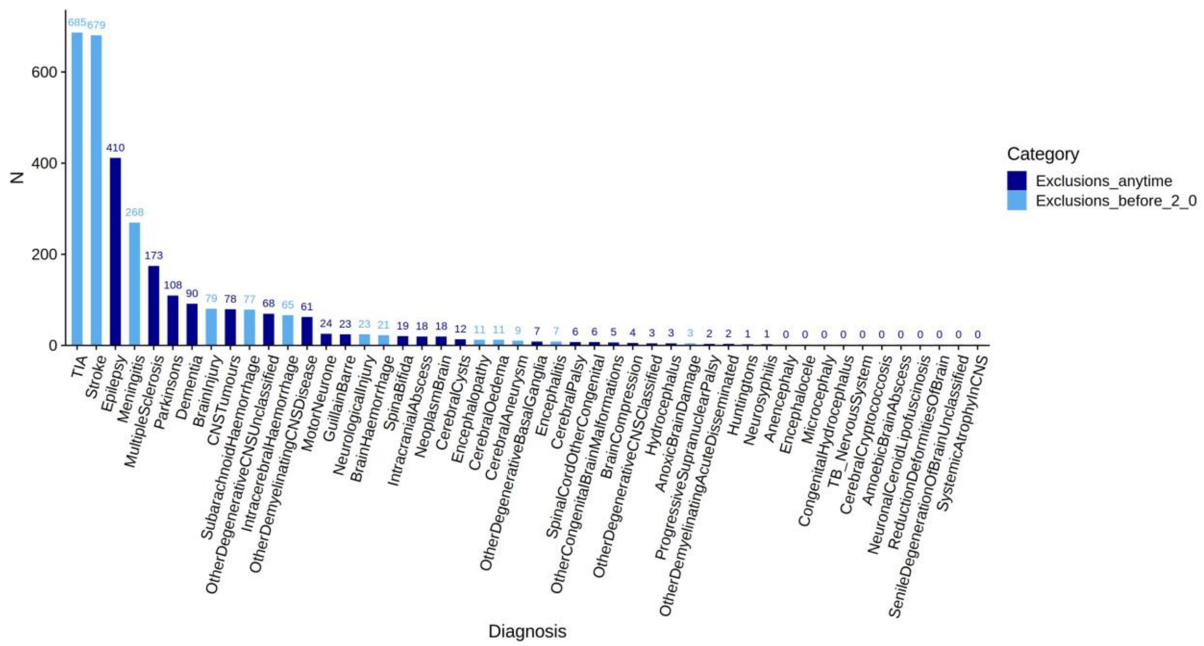


*Figure S3 N* exclusions by diagnosis in the UKB sample (total *N* of participants with any diagnosis = 2542). Initial UKB *N* (participants included in FreeSurfer bulk download) = 40,382 -> UKB *N* = 2542 excluded on medical grounds -> final UKB *N* = 37,840.

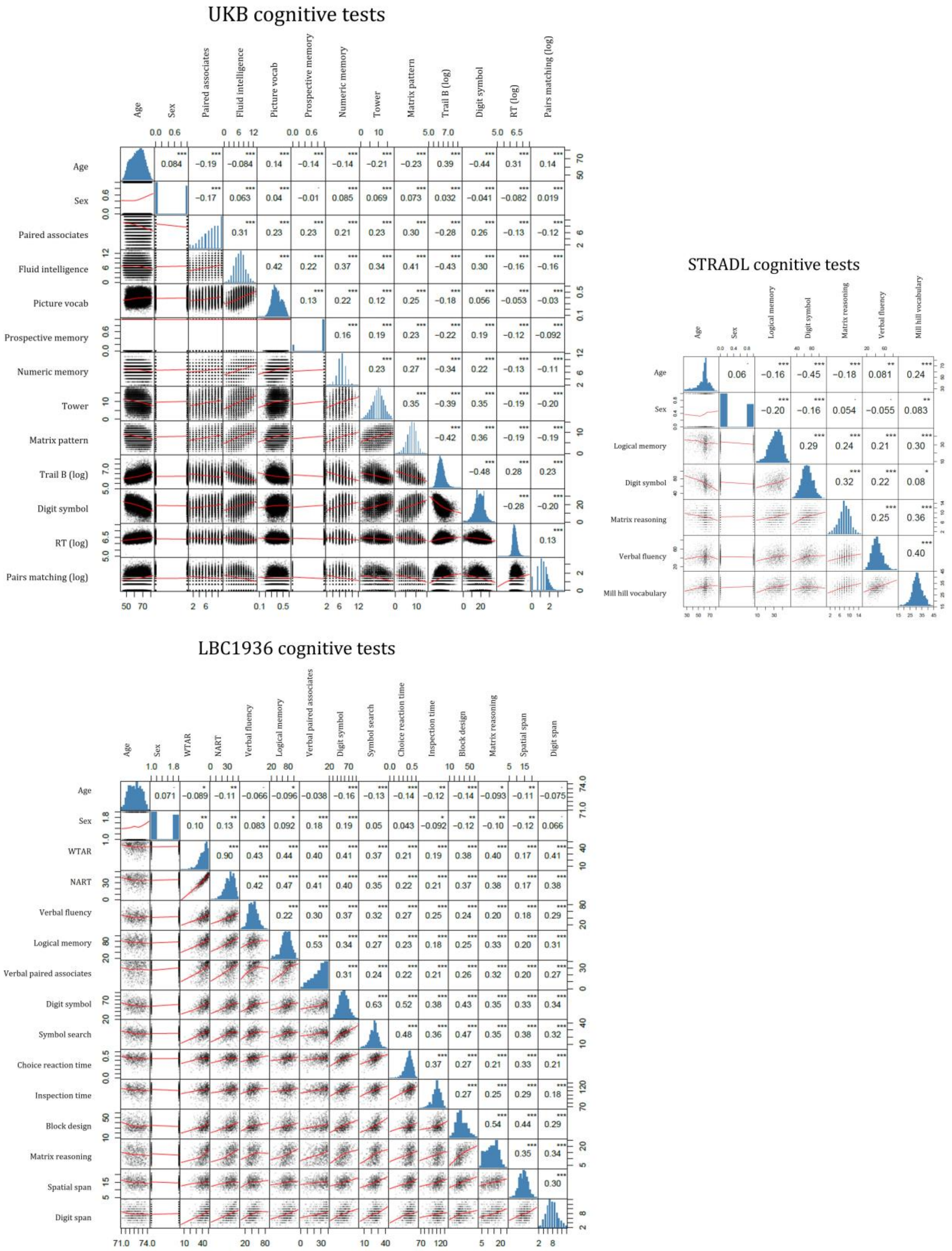


*Figure S4* Correlation plots of the individual cognitive tests within each cohort, also showing data distributions, and correlations with age and sex.

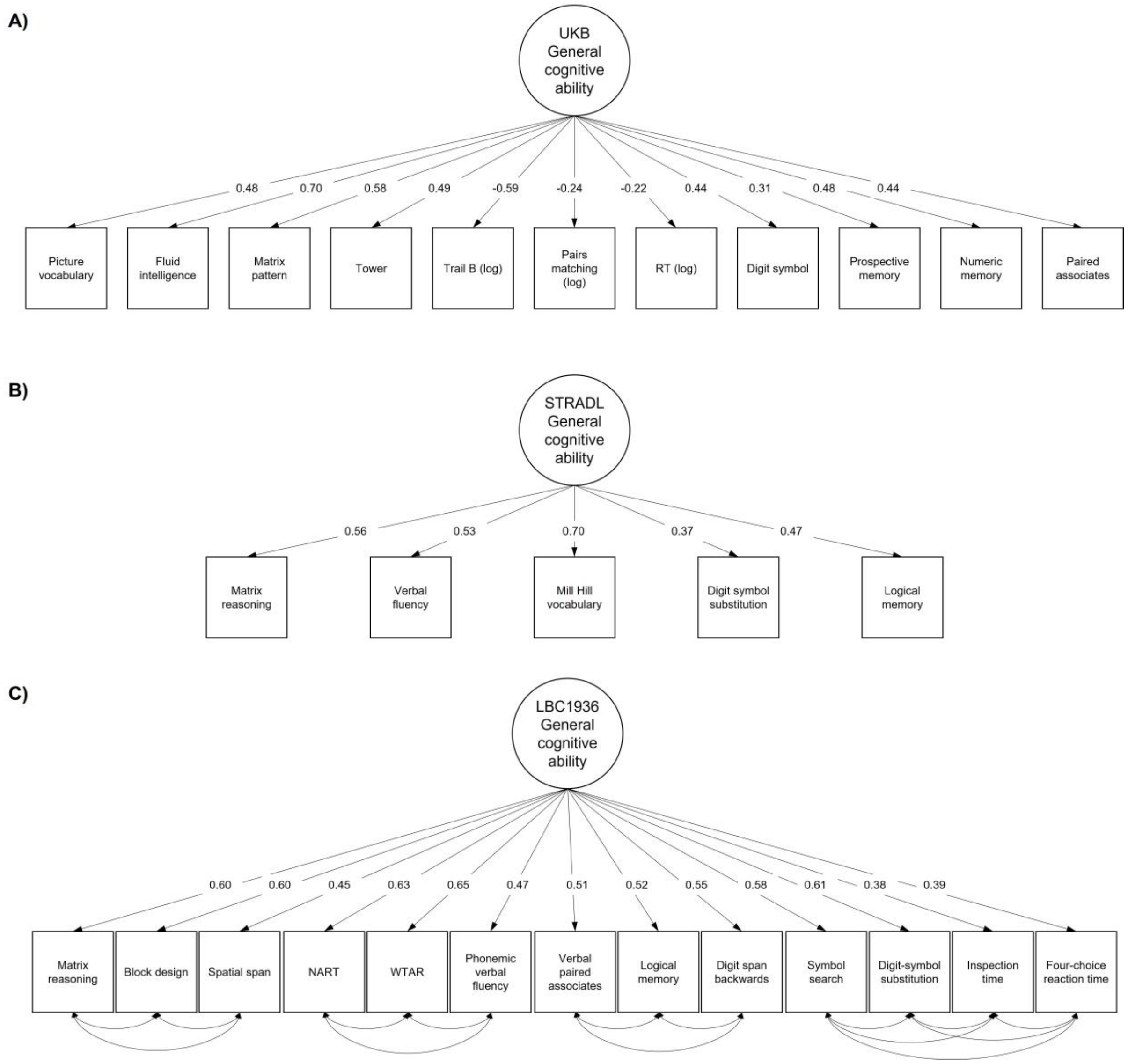


*Figure S5* Simplified path diagrams of cognitive ability latent models alongside density plots of the final cognitive ability prediction *z* scores for A) UKB, B) STRADL, and C) LBC1936. The within-domain residual variances for LBC1936 are in *Table S*17. Model fits are in *Table S18*.

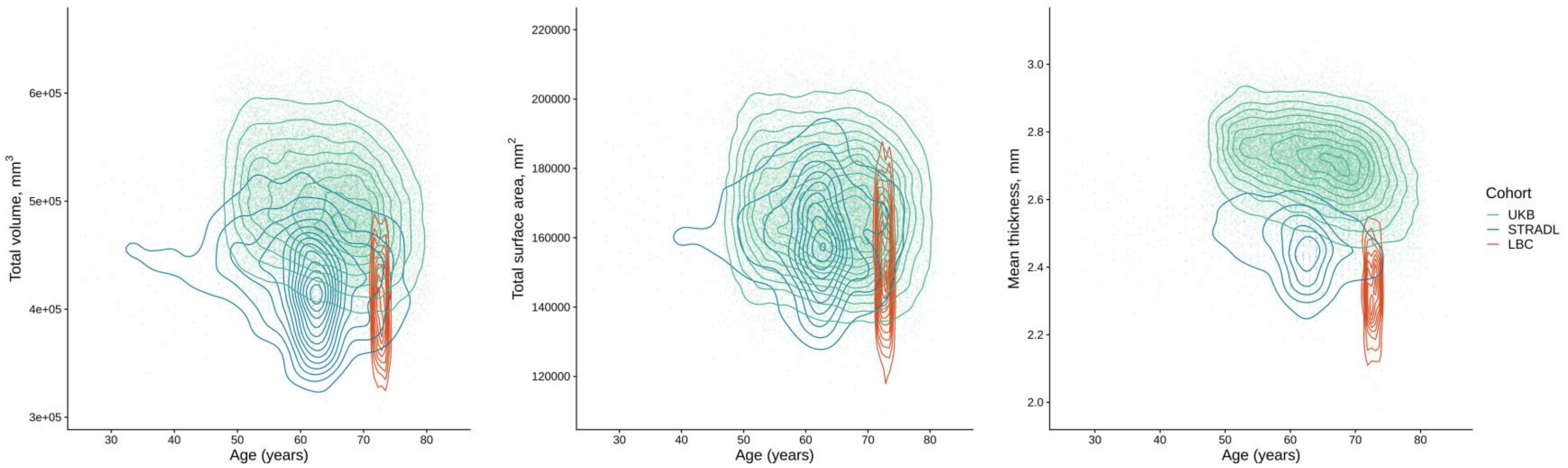


*Figure S6* Total volume (mm3), total surface area (mm2), and mean thickness plotted by age for each cohort (UKB, STRADL and LBC).

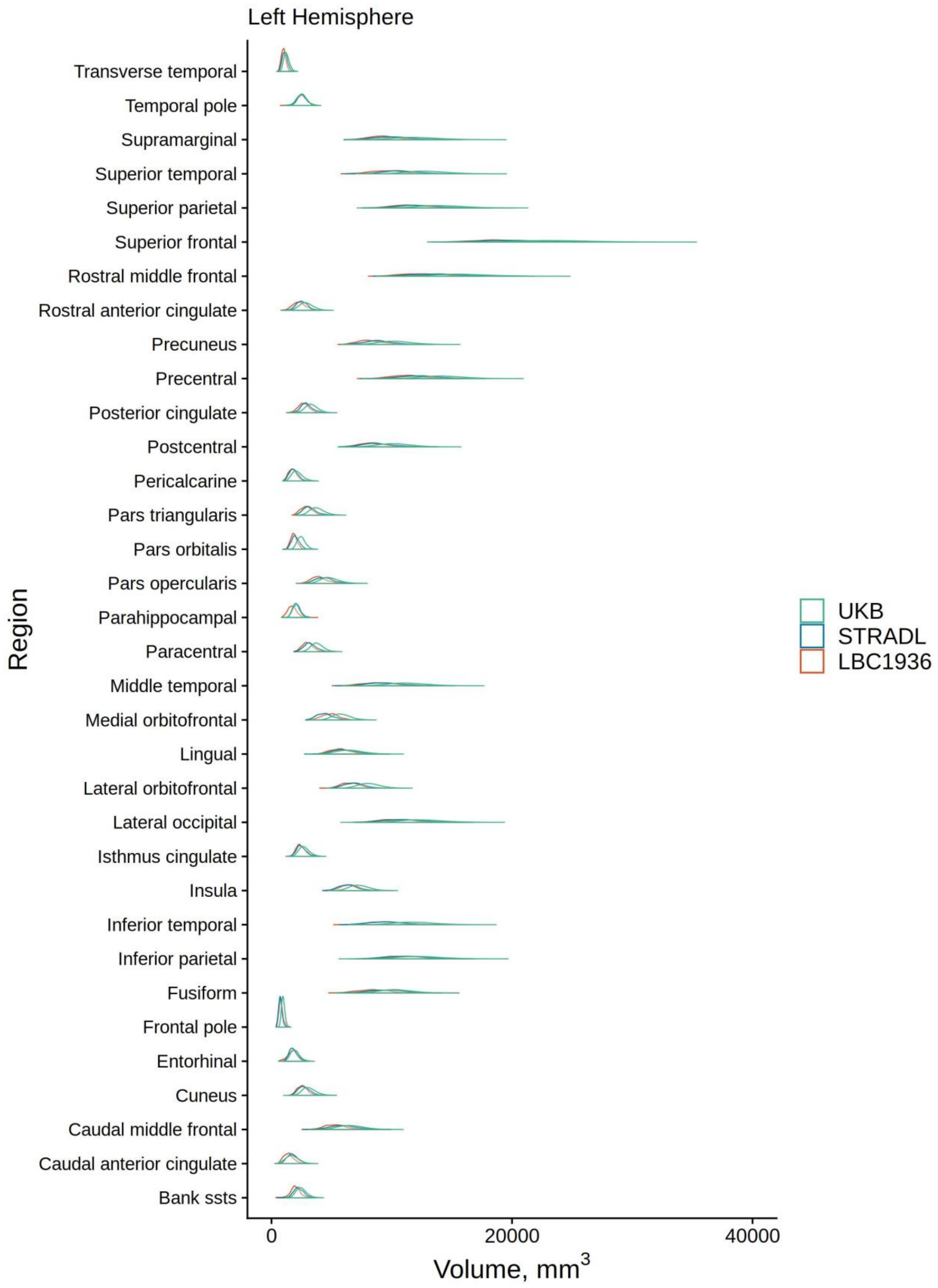


*Figure S7* Regional density plots of left hemisphere volume (mm3), for UKB, STRADL and LBC1936 .

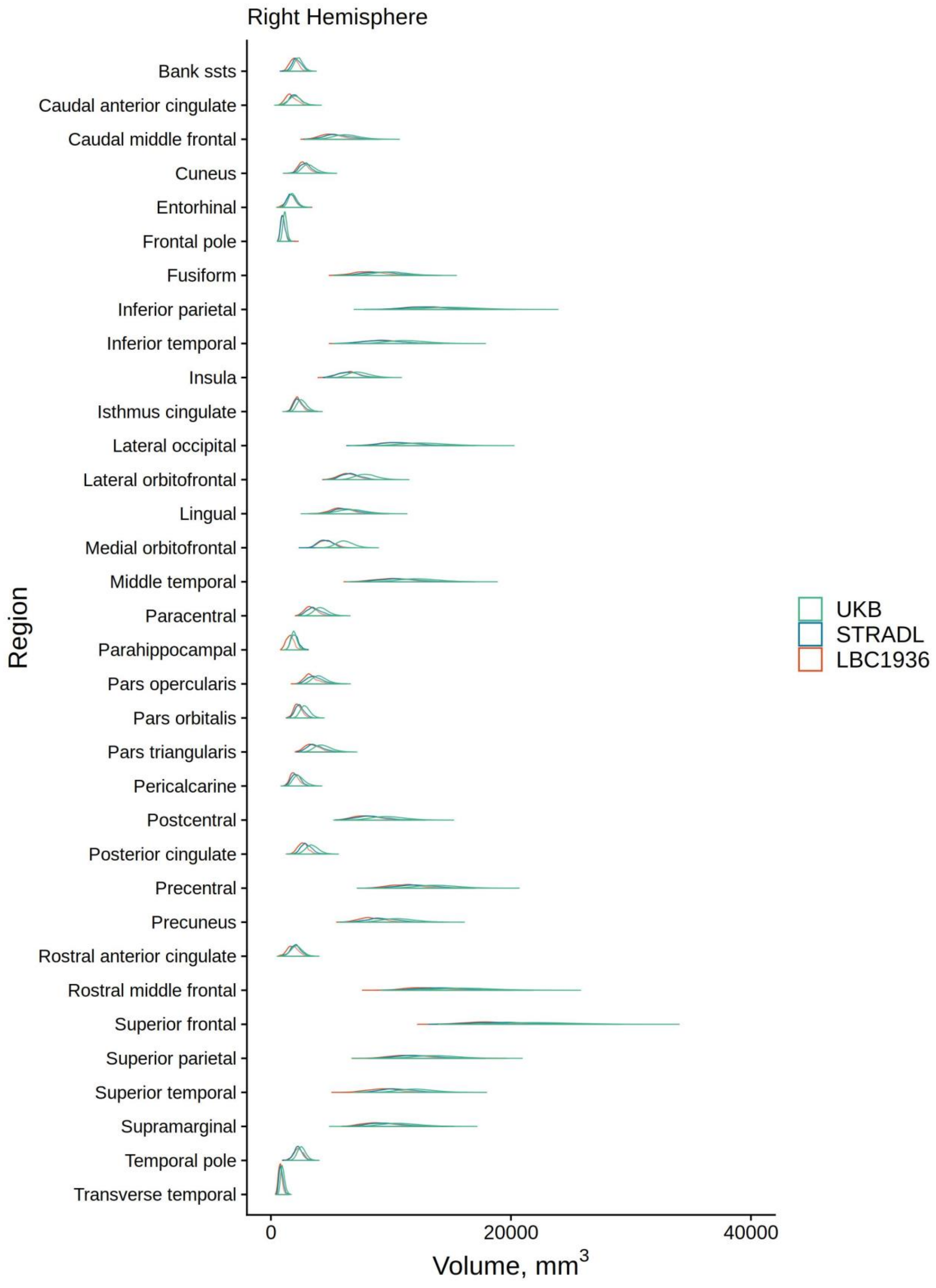


*Figure S8* Regional density plots of right hemisphere volume ($mm^3$), for UKB, STRADL and LBC1936.

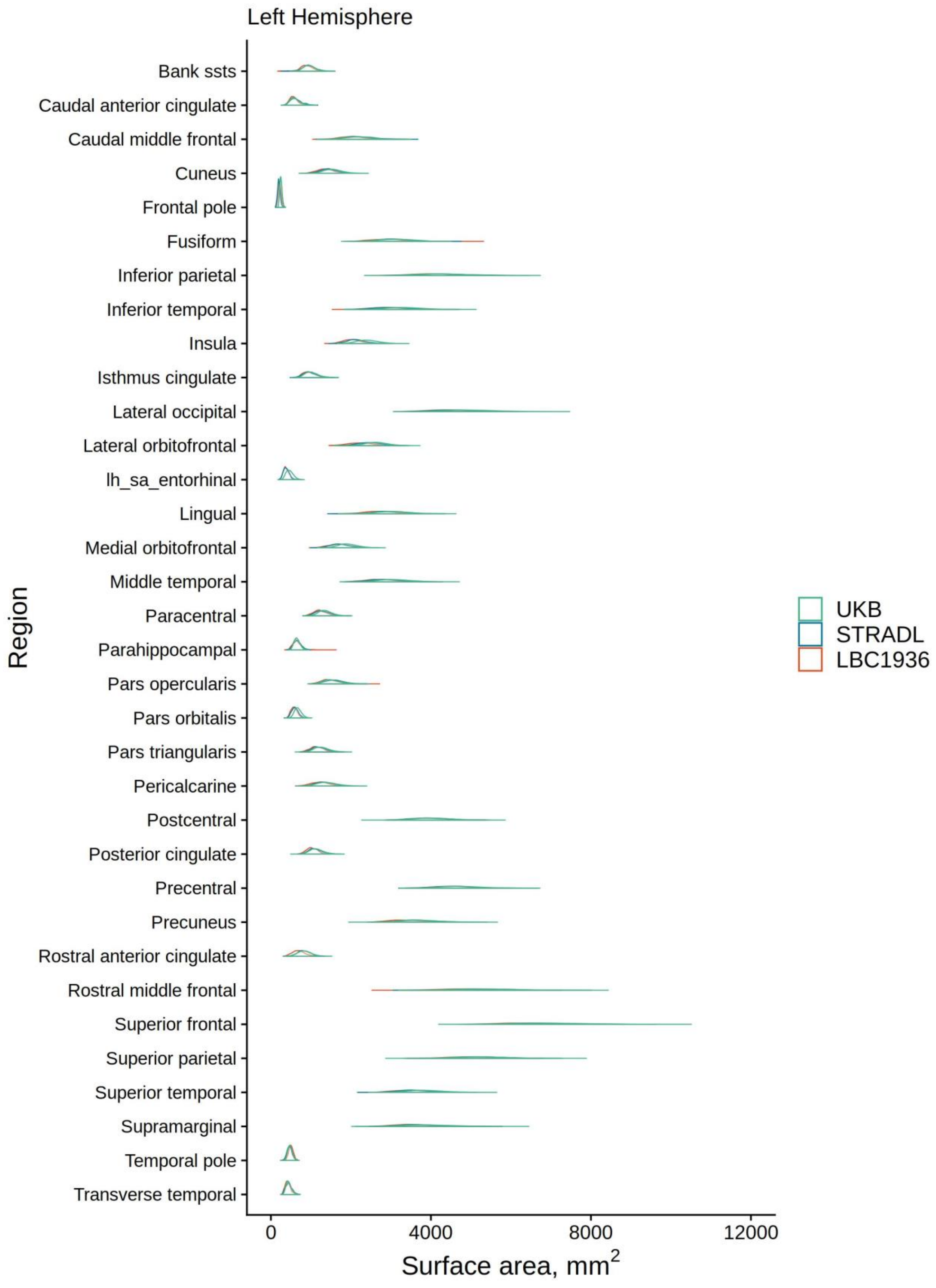


*Figure S9* Regional density plots of left hemisphere surface area (mm²), for UKB, STRADL and LBC1936.

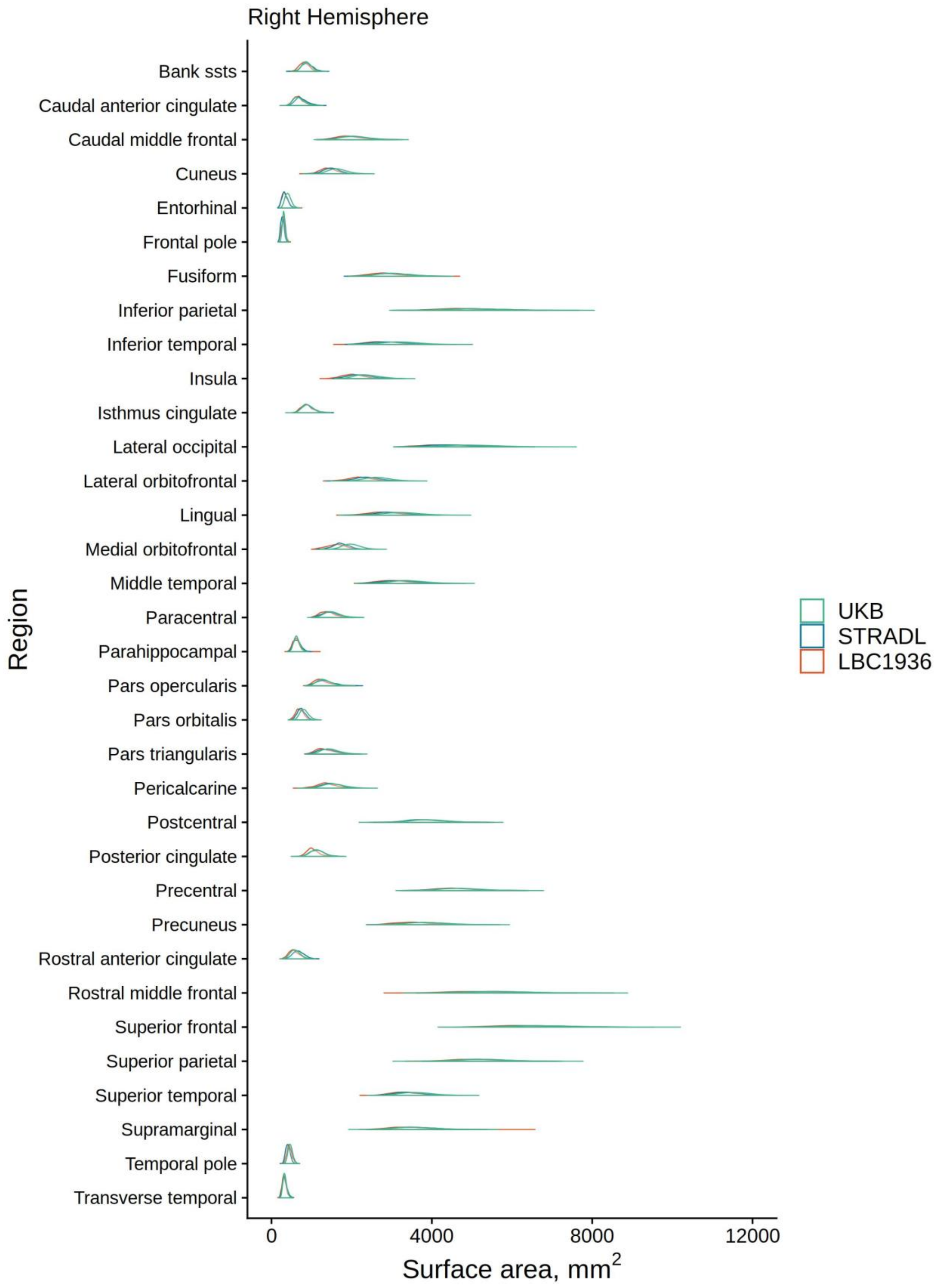


*Figure S10* Regional density plots of right hemisphere surface area (mm²), for UKB, STRADL and LBC1936.

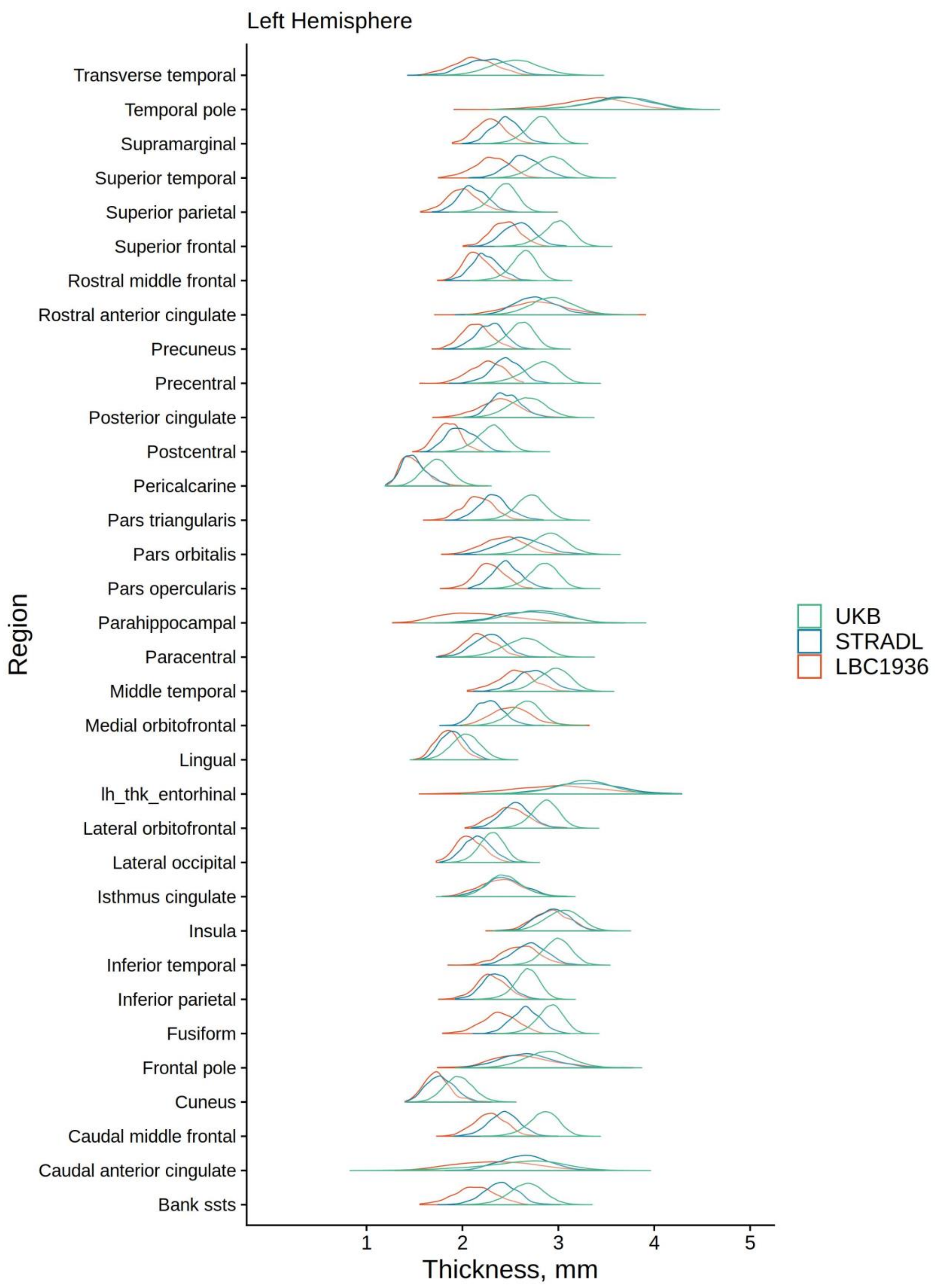


*Figure S11* Regional density plots of left hemisphere thickness (mm), for UKB, STRADL and LBC1936.

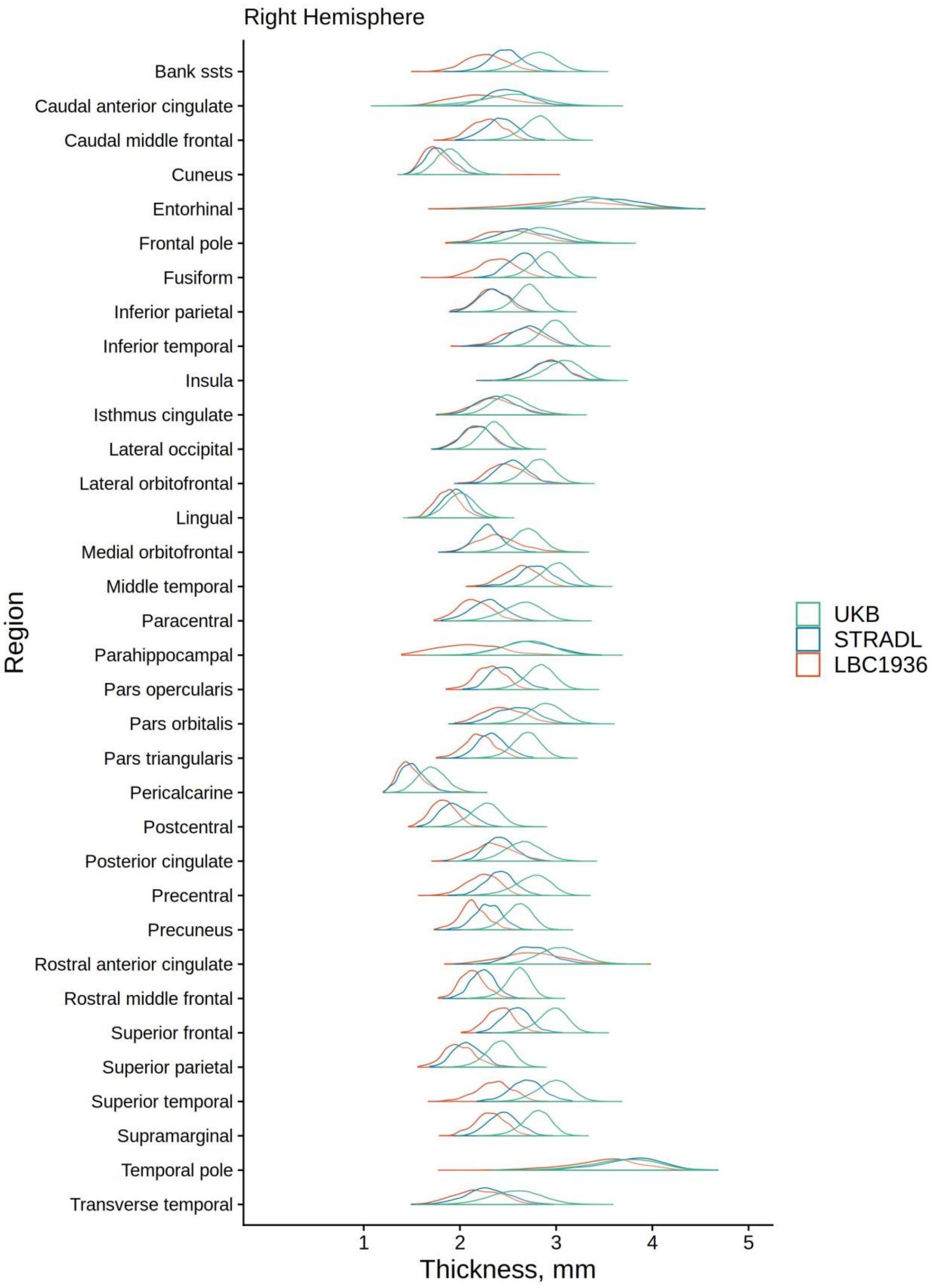


*Figure S12* Regional density plots of right hemisphere thickness (mm), for UKB, STRADL and LBC1936.

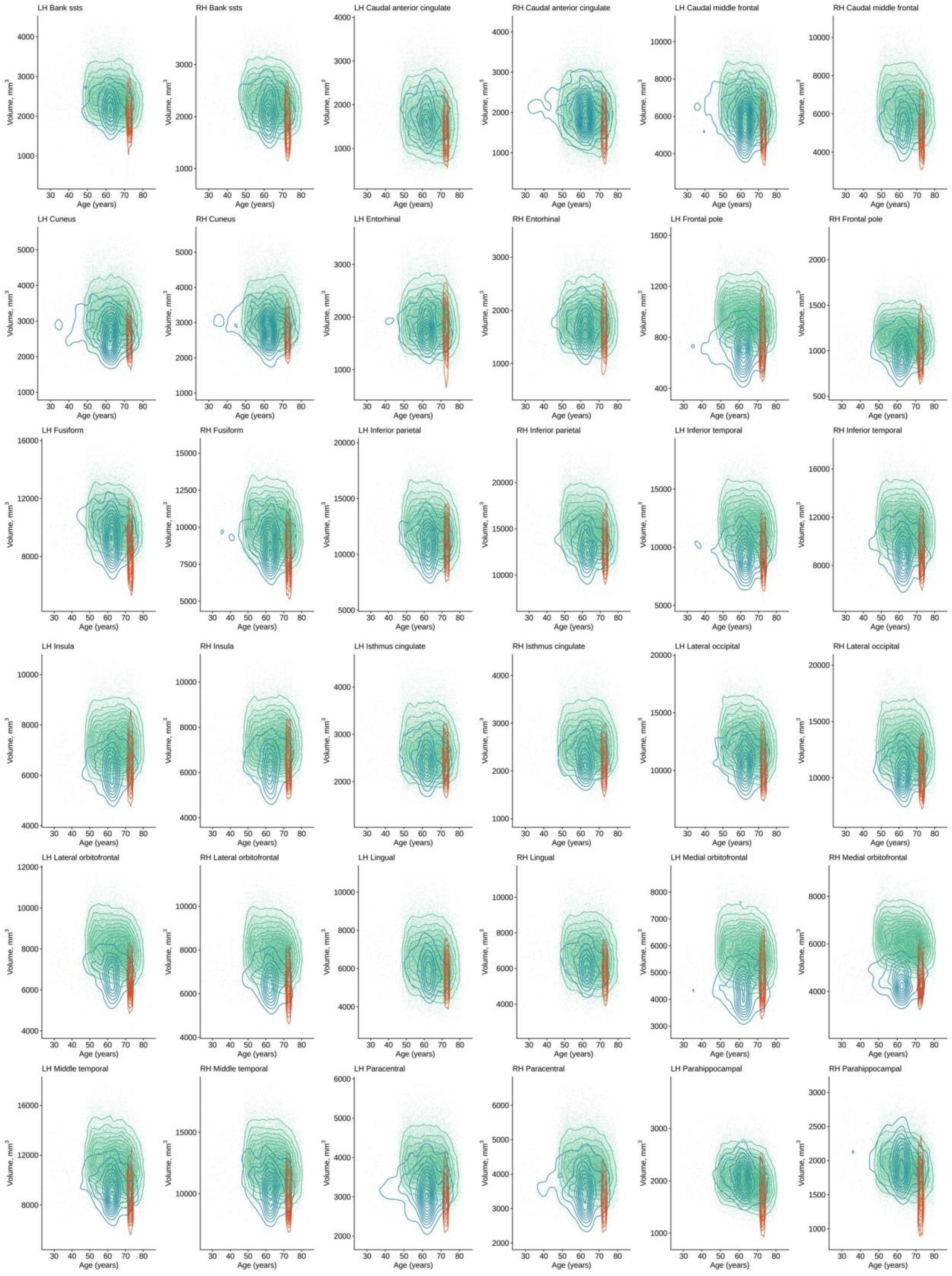

LH Bank ssts
RH Bank ssts
LH Caudal anterior cingulate
RH Caudal anterior cingulate
LH Caudal middle frontal
RH Caudal middle frontal
LH Cuneus
RH Cuneus
LH Entorhinal
RH Entorhinal
LH Frontal pole
RH Frontal pole
LH Fusiform
RH Fusiform
LH Inferior parietal
RH Inferior parietal
LH Inferior temporal
RH Inferior temporal
LH Insula
RH Insula
LH Isthmus cingulate
RH Isthmus cingulate
LH Lateral occipital
RH Lateral occipital
LH Lateral orbitofrontal
RH Lateral orbitofrontal
LH Lingual
RH Lingual
LH Medial orbitofrontal
RH Medial orbitofrontal
LH Middle temporal
RH Middle temporal
LH Paracentral
RH Paracentral
LH Parahippocampal
RH Parahippocampal
Volume, mm3
Age (years)

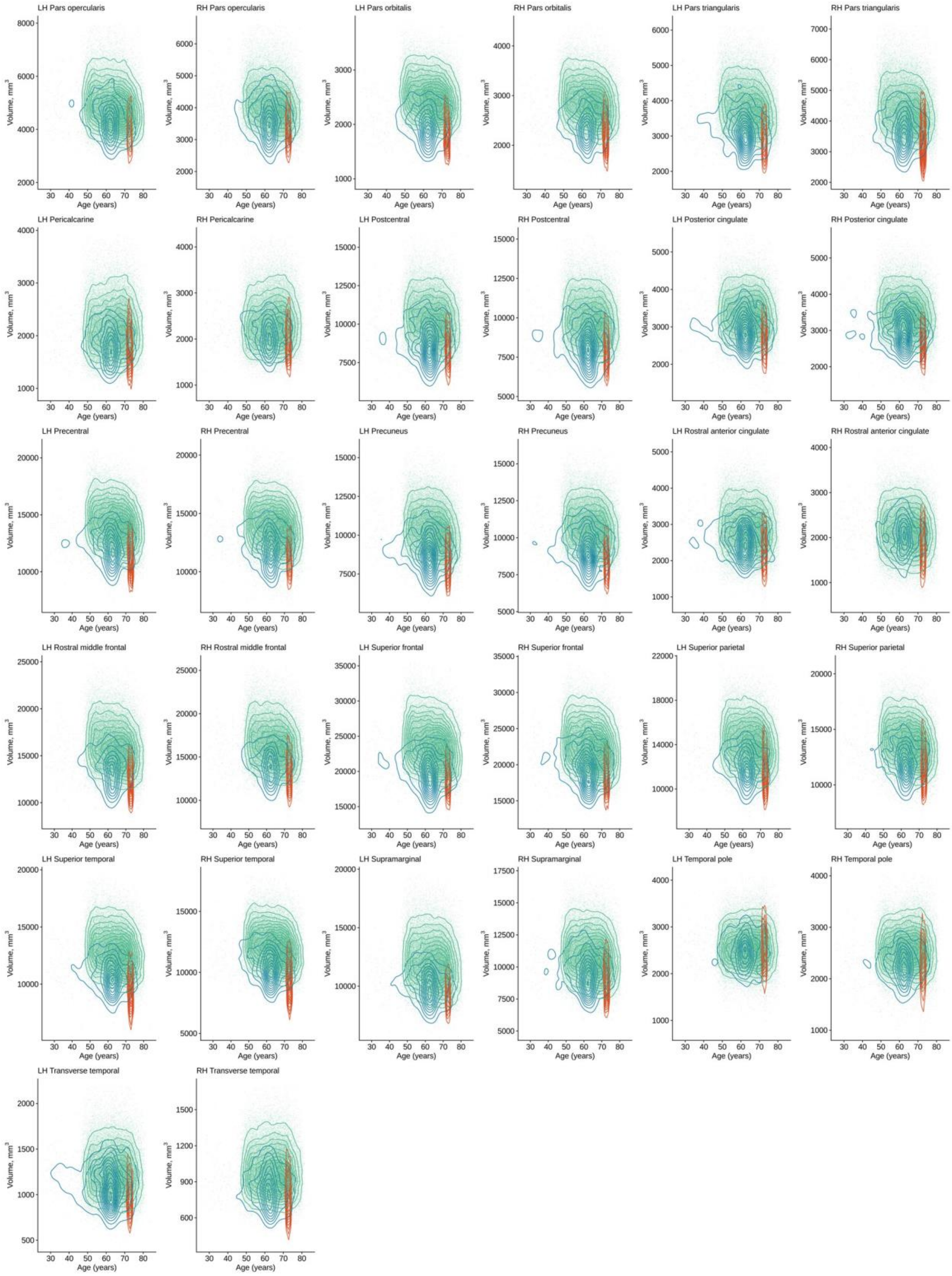
LH Pars opercularis
RH Pars opercularis
LH Pars orbitalis
RH Pars orbitalis
LH Pars triangularis
RH Pars triangularis
LH Pericalcarine
RH Pericalcarine
LH Postcentral
RH Postcentral
LH Posterior cingulate
RH Posterior cingulate
LH Precentral
RH Precentral
LH Precuneus
RH Precuneus
LH Rostral anterior cingulate
RH Rostral anterior cingulate
LH Rostral middle frontal
RH Rostral middle frontal
LH Superior frontal
RH Superior frontal
LH Superior parietal
RH Superior parietal
LH Superior temporal
RH Superior temporal
LH Supramarginal
RH Supramarginal
LH Temporal pole
RH Temporal pole
LH Transverse temporal
RH Transverse temporal
Volume, mm$^3$
Age (years)

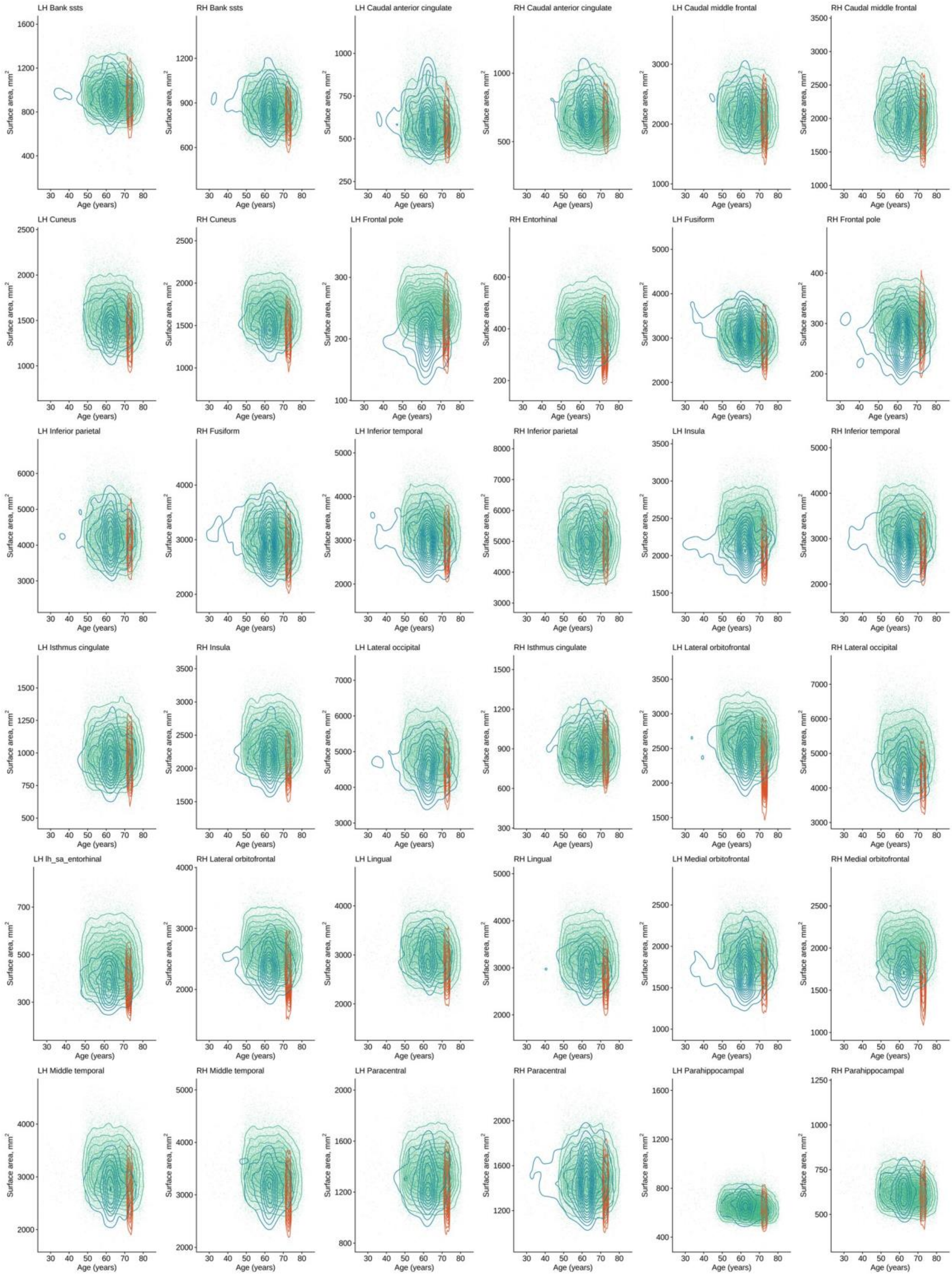
LH Bank ssts
RH Bank ssts
LH Caudal anterior cingulate
RH Caudal anterior cingulate
LH Caudal middle frontal
RH Caudal middle frontal
LH Cuneus
RH Cuneus
LH Frontal pole
RH Entorhinal
LH Fusiform
RH Frontal pole
LH Inferior parietal
RH Fusiform
LH Inferior temporal
RH Inferior parietal
LH Insula
RH Inferior temporal
LH Isthmus cingulate
RH Insula
LH Lateral occipital
RH Isthmus cingulate
LH Lateral orbitofrontal
RH Lateral occipital
LH lh_sa_entorhinal
RH Lateral orbitofrontal
LH Lingual
RH Lingual
LH Medial orbitofrontal
RH Medial orbitofrontal
LH Middle temporal
RH Middle temporal
LH Paracentral
RH Paracentral
LH Parahippocampal
RH Parahippocampal
Surface area, mm$^2$
Age (years)

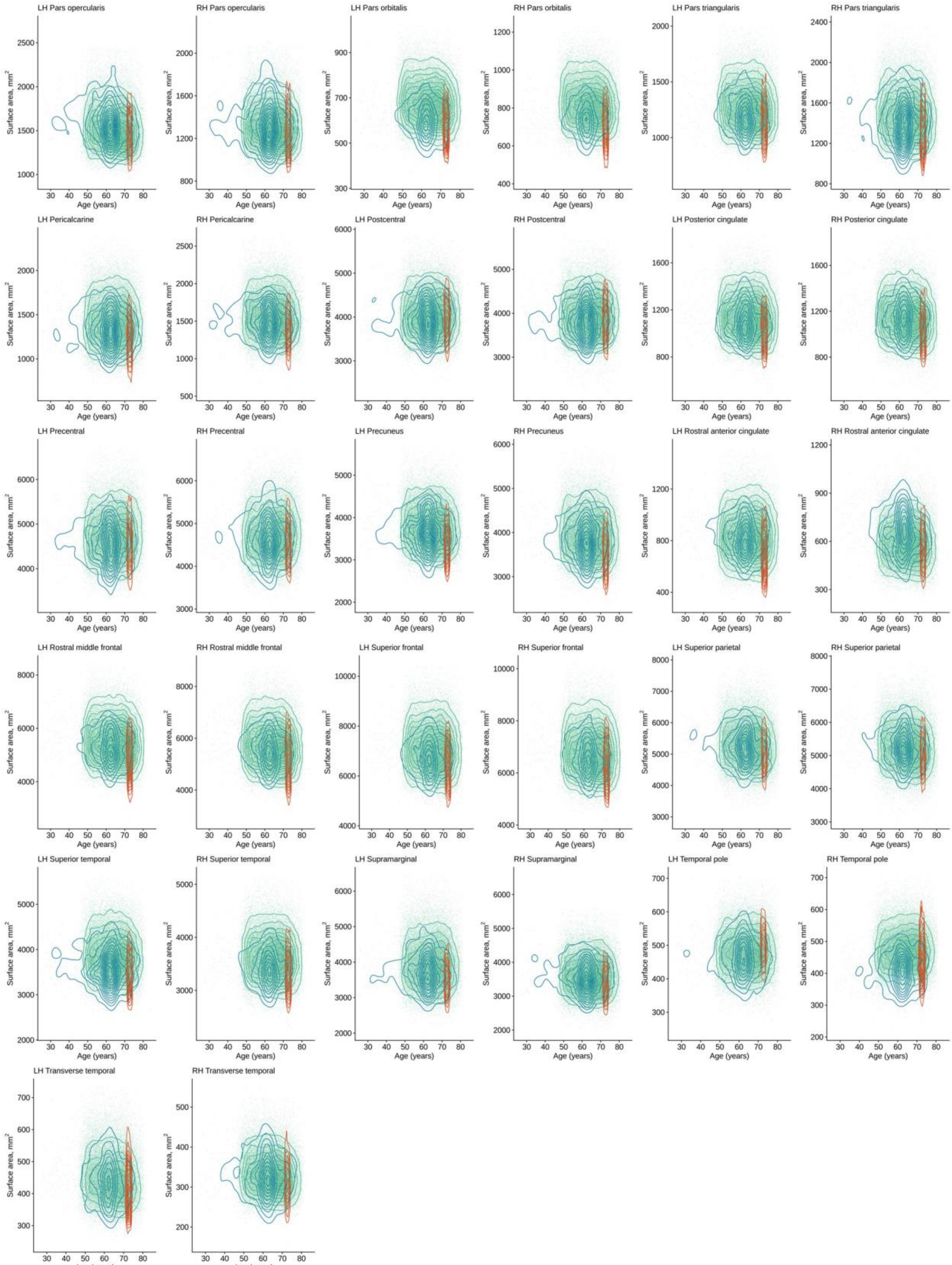

LH Pars opercularis
RH Pars opercularis
LH Pars orbitalis
RH Pars orbitalis
LH Pars triangularis
RH Pars triangularis
LH Pericalcarine
RH Pericalcarine
LH Postcentral
RH Postcentral
LH Posterior cingulate
RH Posterior cingulate
LH Precentral
RH Precentral
LH Precuneus
RH Precuneus
LH Rostral anterior cingulate
RH Rostral anterior cingulate
LH Rostral middle frontal
RH Rostral middle frontal
LH Superior frontal
RH Superior frontal
LH Superior parietal
RH Superior parietal
LH Superior temporal
RH Superior temporal
LH Supramarginal
RH Supramarginal
LH Temporal pole
RH Temporal pole
LH Transverse temporal
RH Transverse temporal
Surface area, mm$^2$
Age (years)

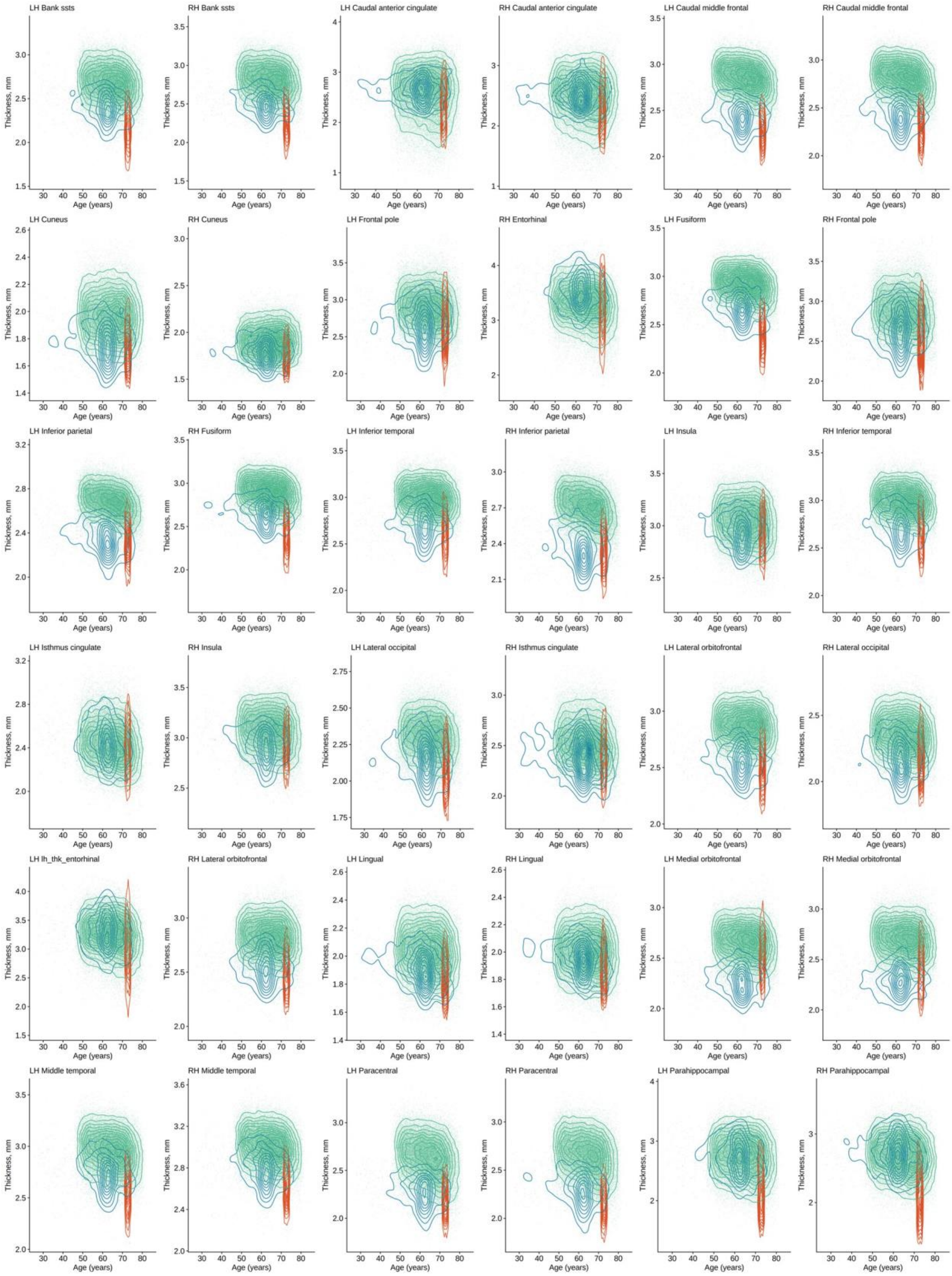
LH Bank ssts
RH Bank ssts
LH Caudal anterior cingulate
RH Caudal anterior cingulate
LH Caudal middle frontal
RH Caudal middle frontal
LH Cuneus
RH Cuneus
LH Frontal pole
RH Entorhinal
LH Fusiform
RH Frontal pole
LH Inferior parietal
RH Fusiform
LH Inferior temporal
RH Inferior parietal
LH Insula
RH Inferior temporal
LH Isthmus cingulate
RH Insula
LH Lateral occipital
RH Isthmus cingulate
LH Lateral orbitofrontal
RH Lateral occipital
LH lh_thk_entorhinal
RH Lateral orbitofrontal
LH Lingual
RH Lingual
LH Medial orbitofrontal
RH Medial orbitofrontal
LH Middle temporal
RH Middle temporal
LH Paracentral
RH Paracentral
LH Parahippocampal
RH Parahippocampal
Thickness, mm
Age (years)

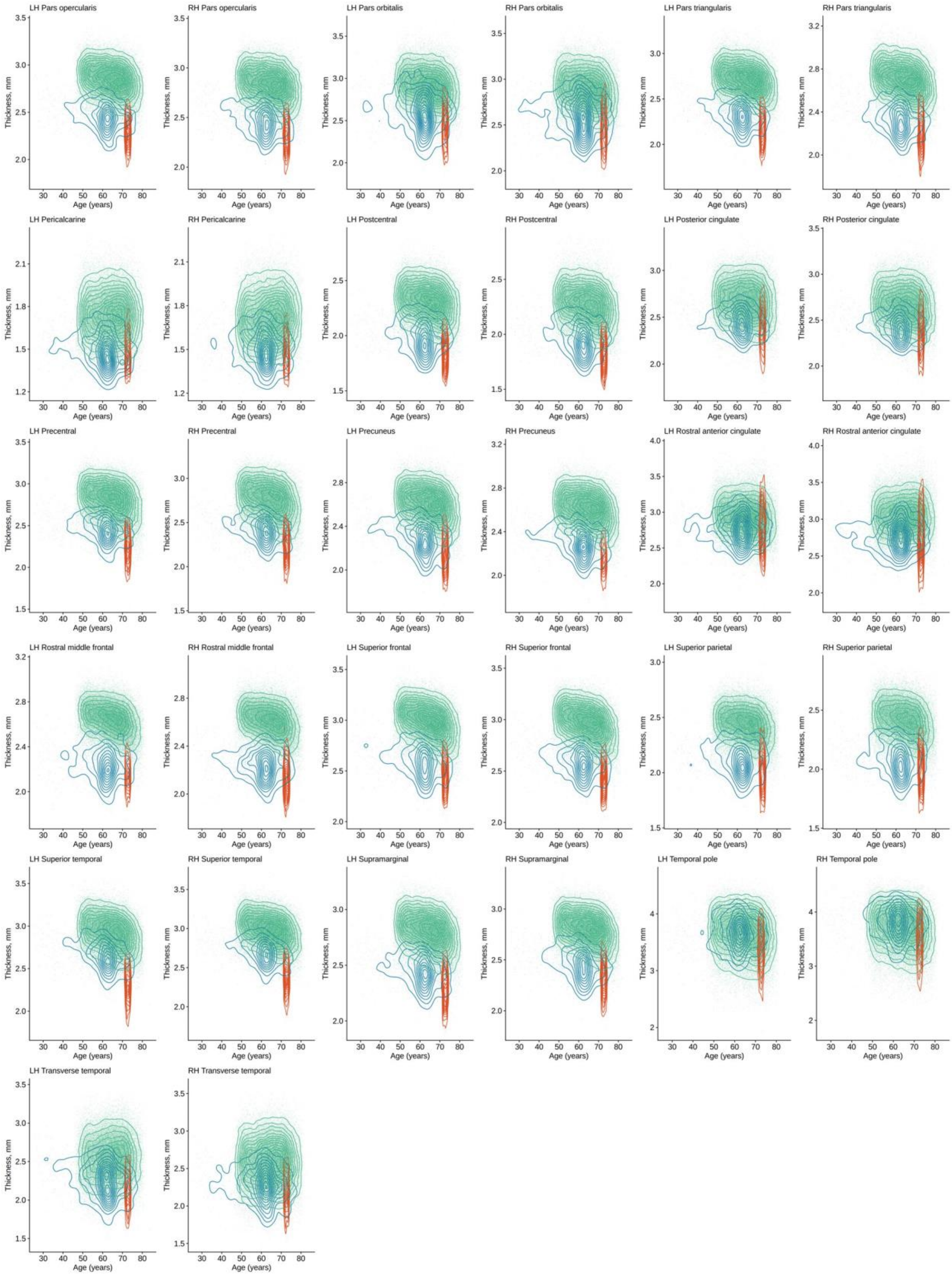


*Figure S13* Regional volume (mm³), surface area (mm²), and thickness (mm) plotted by age for each cohort (UKB, STRADL and LBC).

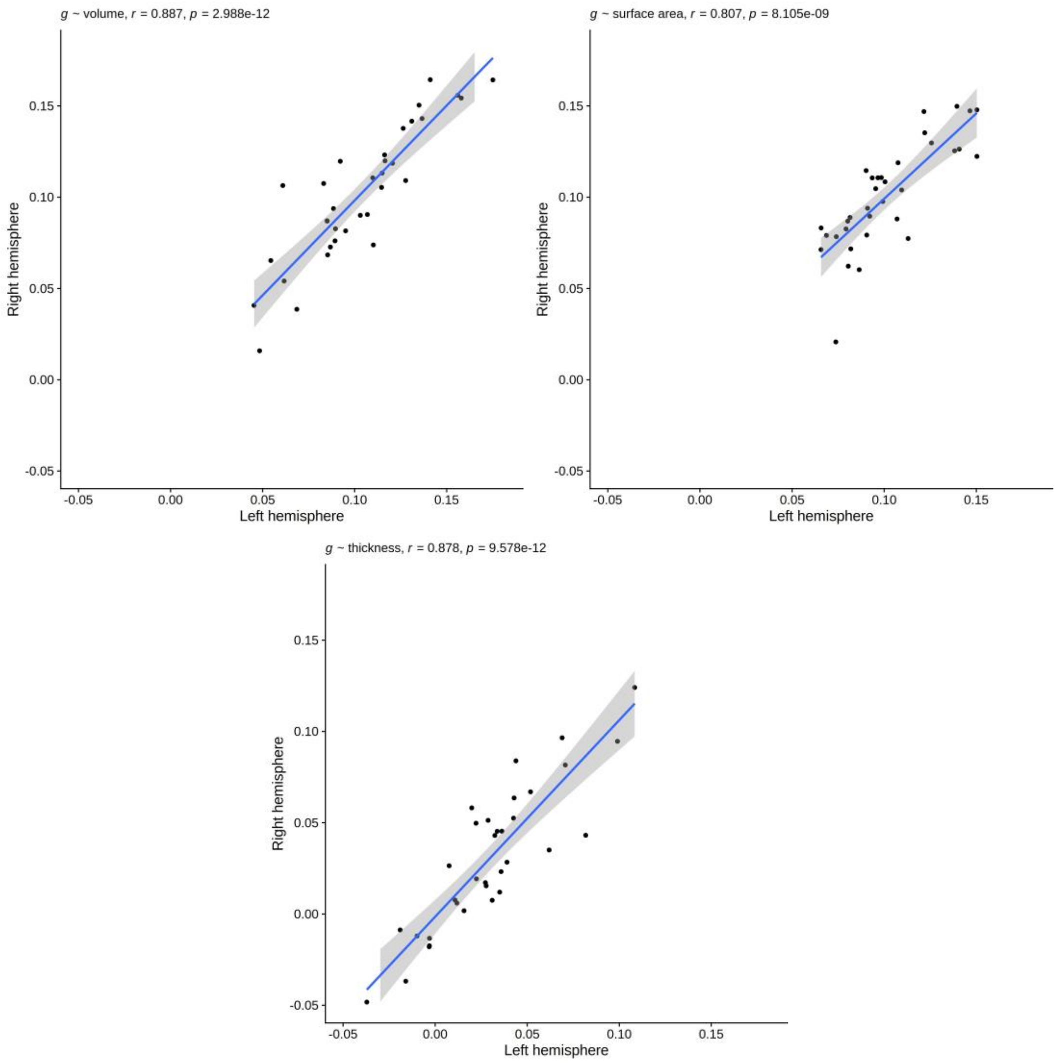


*Figure S14* Inter-hemispheric correlations for meta-analysed g-associations with volume, surface area and thickness.

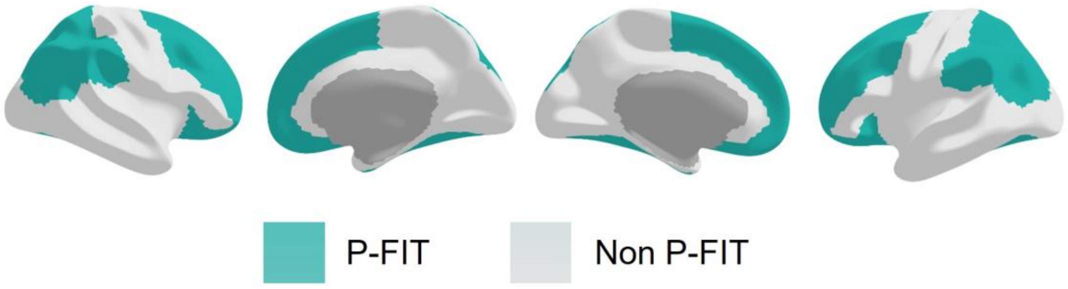


*Figure S15* P-FIT and non-P-FIT regions mapped to the cortex (as in [2]).

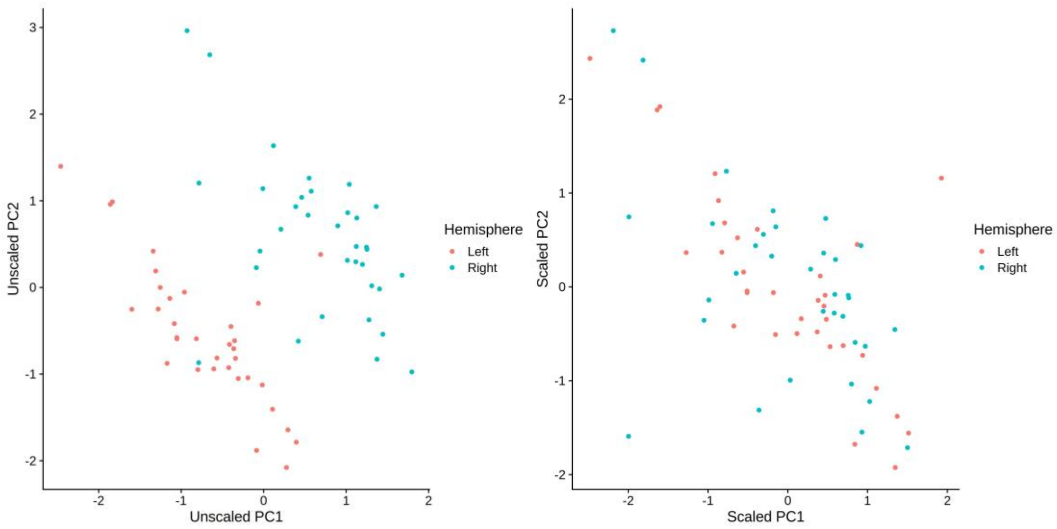


*Figure S16* Rotated component scores (PC1 and PC2), unscaled (left) and scaled by hemisphere (right).

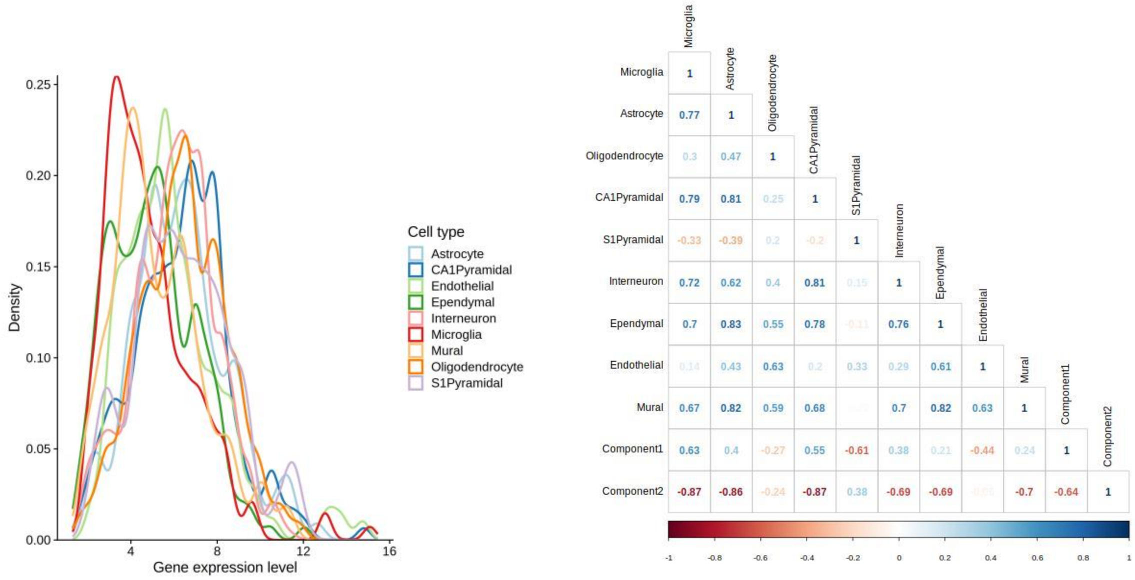


*Figure* S17 Raw expression values by the 9 cell types

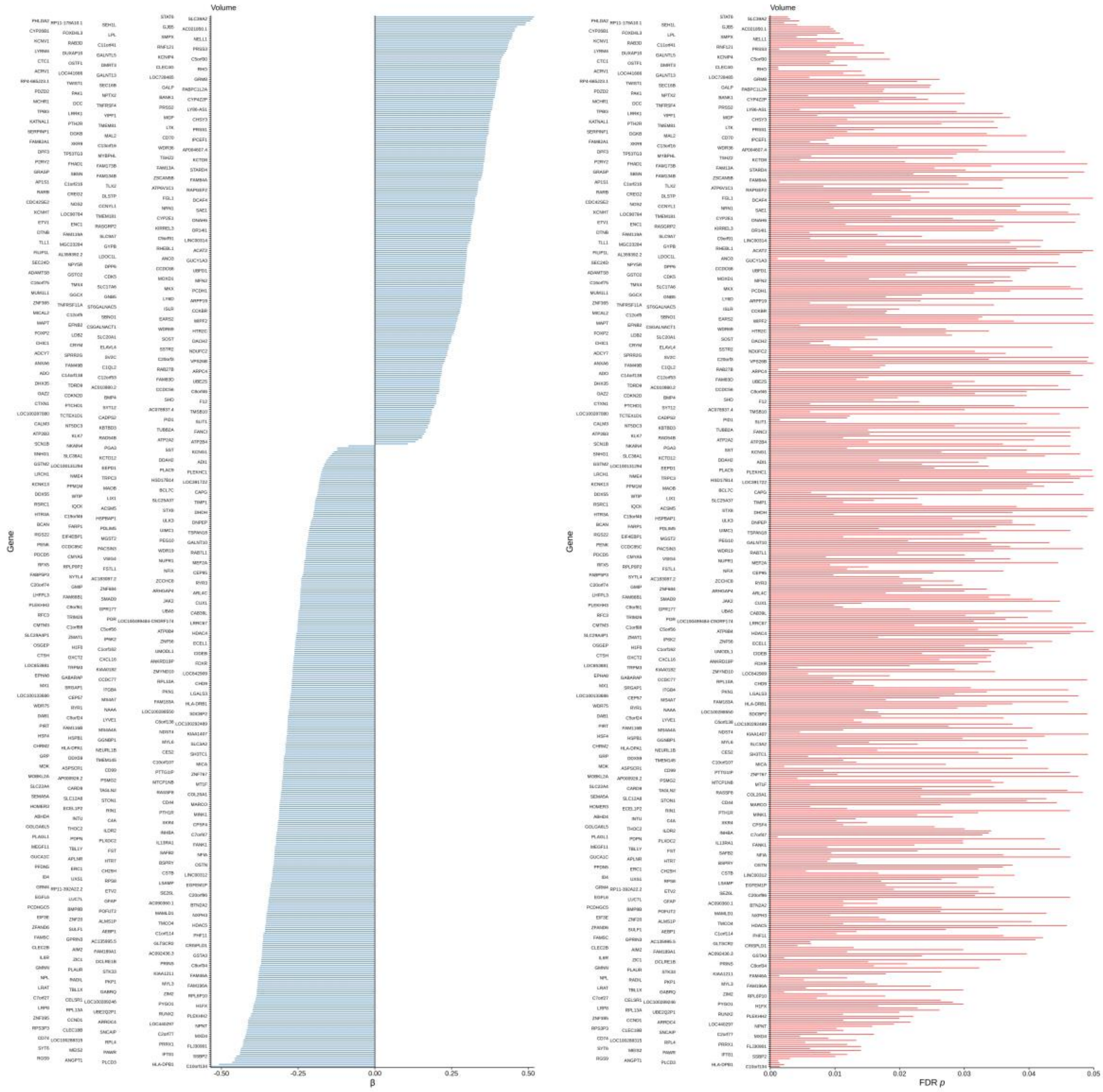


*Figure S18* Regional gene-$g$ associations that have FDR $Q$ values < .05 for volume ($N$ = 522).

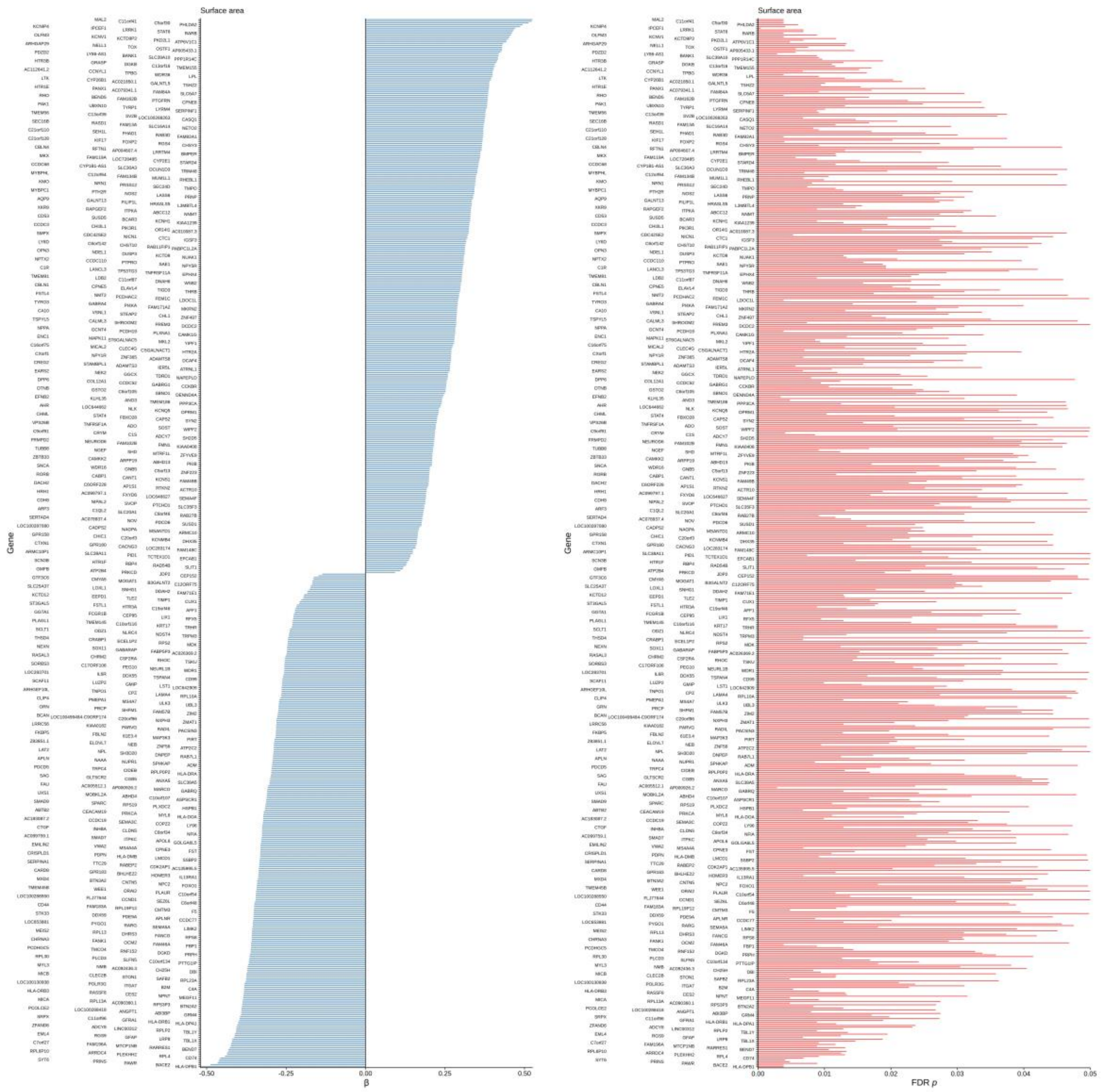


*Figure S19* Regional gene-$g$ associations that have FDR $Q$ values < .05 for surface area ($N$ = 609).

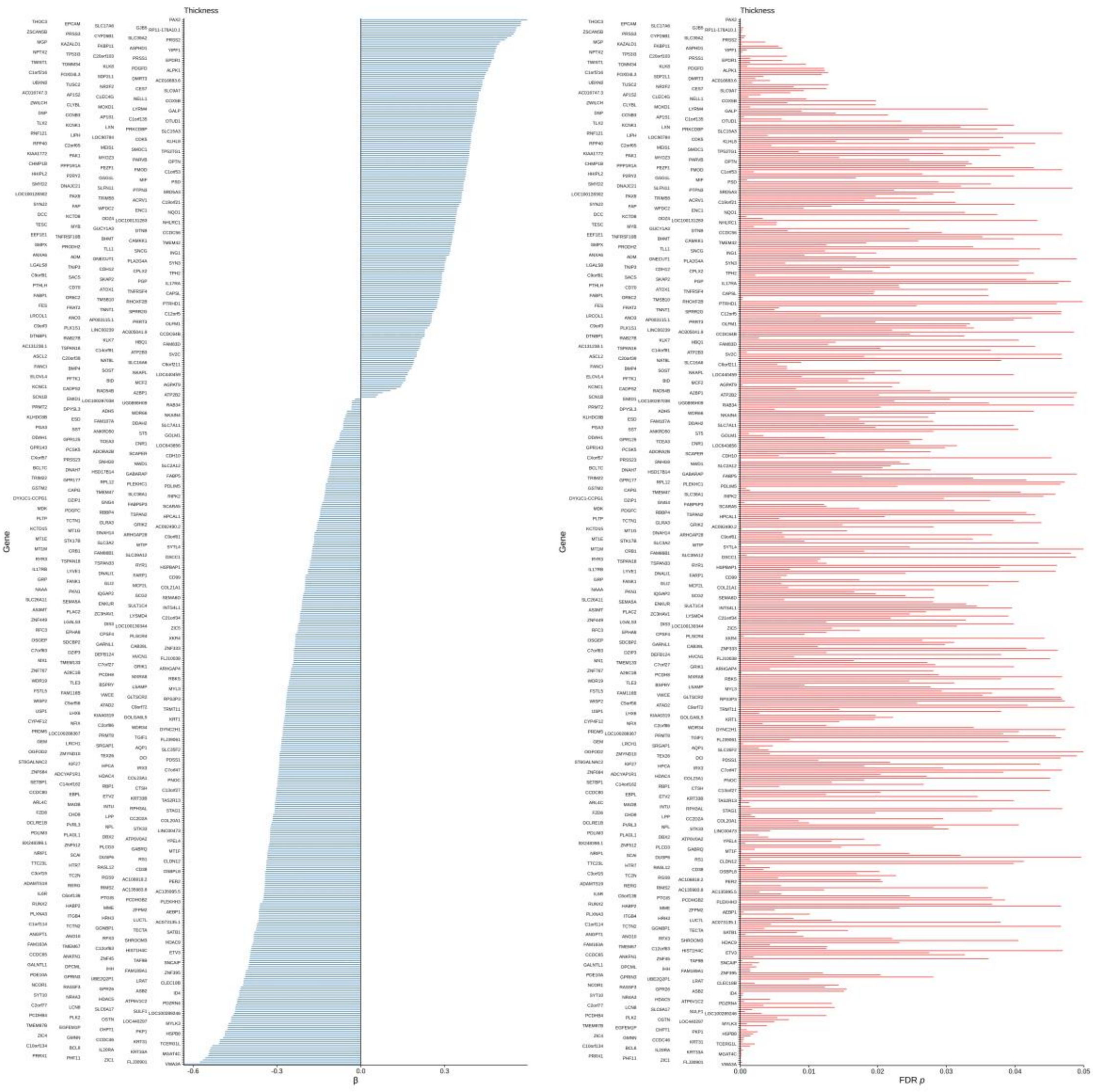


*Figure S20* Regional gene-$g$ associations that have FDR $Q$ values < .05 for thickness ($N$ = 516).

*Table S1* Descriptive statistics of donor characteristics in the subsets of the three gene expression cohorts that are included in current analyses.

| | Age (years) | | | Sex (*N*) | | Donor *N*, *M (SD)* | |
|---|---|---|---|---|---|---|---|
| Dataset | M | SD | Range | Female | Male | Left hemisphere | Right hemisphere |
| Allen Atlas | 42.50 | 13.38 | 24–57 | 1 | 5 | 5.71 (0.72) | 2.00 (0.00) |
| BrainSpan | 33.20 | 6.76 | 23–40 | 2 | 3 | 5 (0) – combined hemispheres | |
| Kang et al. | 35.91 | 8.84 | 23–55 | 4 | 7 | 9.55 (1.04) | 7.19 (0.60) |

*Table S2* Number of donors and samples for each of the 68 regions in the French and Paus (2015) expression matrix (ordered from smallest to largest number of samples in the right hemisphere region).

| | Number of donors | | Number of samples | |
|---|---|---|---|---|
| Region | Left hemisphere | Right hemisphere | Left hemisphere | Right hemisphere |
| Bank ssts | 4 | 2 | 6 | 2 |
| Frontal pole | 3 | 2 | 9 | 2 |
| Pars orbitalis | 6 | 2 | 18 | 3 |
| Entorhinal | 6 | 2 | 18 | 4 |
| Pars opercularis | 5 | 2 | 13 | 4 |
| Pericalcarine | 5 | 2 | 8 | 4 |
| Parstriangularis | 5 | 2 | 7 | 5 |
| Rostral anterior cingulate | 6 | 2 | 18 | 5 |
| Transverse temporal | 6 | 2 | 14 | 5 |
| Cuneus | 6 | 2 | 30 | 6 |
| Caudal anterior cingulate | 6 | 2 | 28 | 7 |
| Lateral occipital | 6 | 2 | 45 | 8 |
| Paracentral | 6 | 2 | 28 | 8 |
| Caudal middle frontal | 6 | 2 | 15 | 9 |
| Isthmus cingulate | 6 | 2 | 24 | 9 |
| Temporal pole | 4 | 2 | 17 | 9 |
| Inferior parietal | 6 | 2 | 37 | 10 |
| Precuneus | 6 | 2 | 40 | 10 |
| Insula | 6 | 2 | 33 | 11 |
| Lingual | 6 | 2 | 45 | 11 |
| Superior parietal | 6 | 2 | 44 | 11 |
| Lateral orbitofrontal | 6 | 2 | 27 | 12 |
| Parahippocampal | 6 | 2 | 33 | 12 |
| Posterior cingulate | 6 | 2 | 37 | 13 |
| Supramarginal | 6 | 2 | 38 | 13 |
| Medial orbitofrontal | 6 | 2 | 54 | 21 |
| Precentral | 6 | 2 | 68 | 23 |
| Fusiform | 6 | 2 | 64 | 24 |
| Rostral middle frontal | 6 | 2 | 47 | 25 |
| Superior temporal | 6 | 2 | 80 | 25 |
| Middle temporal | 6 | 2 | 77 | 27 |
| Postcentral | 6 | 2 | 64 | 27 |
| Superior frontal | 6 | 2 | 100 | 29 |
| Inferior temporal | 6 | 2 | 83 | 34 |

*Table S3* Descriptive statistics for loading distributions by cell types for both the two major components of cortical gene expression.

| | | Component 1 | | | Component 2 | | |
|---|---|---|---|---|---|---|---|
| Cell type | N | Mean *(SD)* | Skewness | Kurtosis | Mean*(SD)* | Skewness | Kurtosis |
| Astrocyte | 129 | -0.04 (0.49) | 0.375 | 1.933 | 0.31 (0.39) | -0.797 | 2.582 |
| CA1 Pyramidal | 204 | -0.03 (0.50) | 0.254 | 1.839 | 0.21 (0.46) | -0.364 | 2.057 |
| Endothelial | 127 | 0.16 (0.47) | -0.277 | 1.982 | 0.08 (0.45) | -0.016 | 1.835 |
| Ependymal | 191 | 0.09 (0.49) | 0.061 | 1.812 | 0.21 (0.45) | -0.471 | 2.229 |
| Interneuron | 181 | 0.06 (0.51) | -0.005 | 1.644 | 0.14 (0.50) | -0.187 | 1.695 |
| Microglia | 185 | -0.11 (0.46) | 0.506 | 2.279 | 0.33 (0.41) | -0.845 | 2.846 |
| Mural | 60 | 0.05 (0.47) | 0.278 | 1.938 | 0.17 (0.48) | -0.207 | 1.752 |
| Oligodendrocyte | 139 | 0.17 (0.48) | -0.228 | 1.850 | 0.18 (0.47) | -0.294 | 1.992 |
| S1 Pyramidal | 155 | 0.10 (0.49) | -0.253 | 1.920 | -0.05 (0.47) | 0.200 | 1.917 |
| Unclassified | 6864 | 0.14 (0.49) | -0.145 | 1.788 | 0.15 (0.46) | -0.145 | 1.788 |

*Table S4* Dunn pairwise comparison results for loadings on Component 1 of cortical gene expression.

| | Astrocyte | | CA1Pyramidal | | Endothelial | | Ependymal | | Interneuron | | Microglia | | Mural | | Oligodendrocyte | | S1 pyramidal | |
|---|---|---|---|---|---|---|---|---|---|---|---|---|---|---|---|---|---|---|
| Cell type | *z* | *p* | *z* | *p* | *z* | *p* | *z* | *p* | *z* | *p* | *z* | *p* | *z* | *p* | *z* | *p* | *z* | *p* |
| CA1Pyramidal | 0.032 | 0.974 | | | | | | | | | | | | | | | | |
| Endothelial | **-3.193** | **0.048** | **-3.563** | **0.014** | | | | | | | | | | | | | | |
| Ependymal | -2.318 | 0.614 | -2.660 | 0.250 | 1.179 | 1.000 | | | | | | | | | | | | |
| Interneuron | -1.866 | 1.000 | -2.141 | 0.903 | 1.590 | 1.000 | 0.474 | 1.000 | | | | | | | | | | |
| Microglia | 1.148 | 1.000 | 1.261 | 1.000 | **4.606** | **1.73e-04** | **3.837** | **0.005** | **3.316** | **0.032** | | | | | | | | |
| Mural | -1.231 | 1.000 | -1.335 | 1.000 | 1.320 | 1.000 | 0.485 | 1.000 | 0.152 | 1.000 | -2.181 | 0.846 | | | | | | |
| Oligodendrocyte | **-3.534** | **0.015** | **-3.962** | **0.003** | -0.269 | 1.000 | -1.506 | 1.000 | -1.925 | 1.000 | **-5.022** | *** | -1.552 | 1.000 | | | | |
| S1Pyramidal | -2.437 | 0.459 | -2.760 | 0.191 | 0.908 | 1.000 | -0.243 | 1.000 | -0.689 | 1.000 | **-3.876** | **0.004** | -0.645 | 1.000 | 1.212 | 1.000 | | |
| Unclassified | **-4.051** | **0.002** | **-5.118** | **1.36E-05** | 0.437 | 1.000 | -1.307 | 1.000 | -1.925 | 1.000 | **-6.599** | **1.86E-09** | -1.293 | 1.000 | 0.841 | 1.000 | -0.856 | 1.000 |

*Table S5* Dunn pairwise comparison results for loadings on Component 2 of cortical gene expression.

| | Astrocyte | | CA1 Pyramidal | | Endothelial | | Ependymal | | Interneuron | | Microglia | | Mural | | Oligodendrocyte | | S1 pyramidal | |
|---|---|---|---|---|---|---|---|---|---|---|---|---|---|---|---|---|---|---|
| Cell type | *z* | *p* | *z* | *p* | *z* | *p* | *z* | *p* | *z* | *p* | *z* | *p* | *z* | *p* | *z* | *p* | *z* | *p* |
| CA1 Pyramidal | 1.839 | 1.000 | | | | | | | | | | | | | | | | |
| Endothelial | **3.950** | **0.003** | 2.538 | 0.312 | | | | | | | | | | | | | | |
| Ependymal | 1.929 | 1.000 | 0.129 | 0.897 | -2.393 | 0.452 | | | | | | | | | | | | |
| Interneuron | 3.105 | 0.063 | 1.478 | 1.000 | -1.175 | 1.000 | 1.330 | 1.000 | | | | | | | | | | |
| Microglia | -0.499 | 1.000 | -2.602 | 0.269 | **-4.782** | ******* | -2.686 | 0.217 | **-3.970** | **0.003** | | | | | | | | |
| Mural | 1.842 | 1.000 | 0.551 | 1.000 | -1.314 | 1.000 | 0.460 | 1.000 | -0.470 | 1.000 | 2.323 | 0.525 | | | | | | |
| Oligodendrocyte | 2.185 | 0.722 | 0.548 | 1.000 | -1.846 | 1.000 | 0.424 | 1.000 | -0.804 | 1.000 | 2.890 | 0.119 | -0.134 | 1.000 | | | | |
| S1Pyramidal | **6.334** | **1.05E-08** | **5.143** | **1.14E-05** | 2.181 | 0.700 | **4.949** | **2.98E-05** | **3.628** | **9.71E-03** | **7.458** | **3.95E-12** | 3.071 | 0.068 | **4.175** | **0.001** | | |
| Unclassified | **3.894** | **0.003** | 1.959 | 1.000 | -1.650 | 1.000 | 1.721 | 1.000 | -0.156 | 1.000 | **5.413** | **2.66E-06** | 0.449 | 1.000 | 0.921 | 1.000 | **-5.033** | **1.98E-05** |

*Table S6* The initial number of retained genes, and the number of genes that were matched to French and Paus' post-consistency check genes, alongside processing choices for each of the pipelines provided by Markello et al. (2021). Highlighted cells may help to explain some of the low factor congruence coefficients for PC2, compared to the French and Paus dataset.

| Pipeline | Start *N* genes | *N* matched genes | reannotated | corrected_mni | tolerance (mm) | sample_norm | gene_norm | missing | probe_selection | donor_probes | region_agg | agg_metric | lr_mirror | ibf_threshold | norm_matched | norm_structures | sim_threshold |
|---|---|---|---|---|---|---|---|---|---|---|---|---|---|---|---|---|---|
| French & Paus (2015) | 20739 | 8108 | FALSE | FALSE | 1 | None | None | None | average | aggregate | donors | median | none | 0 | TRUE | FALSE | none |
| Anderson (2018) | 20739 | 8108 | FALSE | FALSE | 0 | None | center | None | corr_intensity | aggregate | donors | mean | none | 0 | TRUE | TRUE | none |
| Burt (2018) | 20739 | 8108 | FALSE | FALSE | 2 | zscore | zscore | interpolate | corr_variance | independent | donors | mean | none | 0 | TRUE | FALSE | 5 |
| Hawrylycz (2015) | 20739 | 8108 | FALSE | FALSE | 0 | None | zscore | cetroids | diff_stability | aggregate | donors | mean | none | 0 | TRUE | FALSE | none |
| Krienen (2016) | 20739 | 8108 | FALSE | FALSE | 0 | None | center | None | average | aggregate | donors | mean | mpme | 0 | TRUE | FALSE | none |
| Whitaker (2016) | 20739 | 8108 | FALSE | FALSE | 0 | None | zscore | centroids | average | aggregate | donors | mean | none | 0 | TRUE | FALSE | none |
| Romero-Garcia (2018) | 20233 | 6800 | TRUE | FALSE | 0 | None | zscore | interpolate | max_intensity | aggregate | samples | median | rightleft | 0 | TRUE | FALSE | none |
| Anderson (2020) | 17383 | 7800 | FALSE | FALSE | -4 | zscore | zscore | None | max_intensity | aggregate | donors | mean | none | 0.2 | TRUE | FALSE | none |
| Liu (2020) | 15631 | 6164 | TRUE | FALSE | 5 | zscore | zscore | | average | aggregate | donors | mean | none | 0.5 | FALSE | FALSE | none |
| Markello (2021) | 15634 | 6166 | TRUE | TRUE | 2 | srs | srs | None | diff_stability | aggregate | donors | mean | none | 0.5 | TRUE | FALSE | none |

*Table S7* Number of retained genes in each validation dataset after within-sample between-donor consistency measures, and the number that were matched with the 8235 consistent genes in the Allen dataset.

| Data source | Start (*N*) | Consistent genes (*N*) | Matched with Allen 8235 genes (*N*) |
|---|---|---|---|
| BrainSpan | 17604 | 4110 | 2250 |
| Kang left hemisphere | 17565 | 2030 | 1554 |
| Kang right hemisphere | 17565 | 2778 | 1784 |
| Kang both hemispheres | 17565 | 1048 | 908 |

*Table S8* The initial number of retained genes, and the number of genes that were matched to French and Paus' post-consistency check gene for each pipeline following Markello et al.'s scripts([3]).

| Pipeline | Start *N* | Matched *N* |
|---|---|---|
| Anderson (2018) | 20739 | 8108 |
| Burt (2018) | 20739 | 8108 |
| Hawrylycz (2015) | 20739 | 8108 |
| Krienen (2016) | 20739 | 8108 |
| Whitaker (2016) | 20739 | 8108 |
| Romero-Garcia (2018) | 20233 | 6800 |
| Anderson (2020) | 17383 | 7800 |
| Liu (2020) | 15631 | 6164 |
| Markello (2021) | 15634 | 6166 |

*Table S9* Regional component scores for the two major components of gene expression.

| Hemisphere | Region | Component 1 | Component 2 |
|---|---|---|---|
| Left | Bank ssts | -0.382 | 0.613 |
| | Caudal anterior cingulate | 0.841 | -1.677 |
| | Caudal middle frontal | -0.180 | -0.061 |
| | Cuneus | -1.603 | 1.921 |
| | Entorhinal | 1.347 | -1.925 |
| | Frontal pole | 1.926 | 1.159 |
| | Fusiform | 0.531 | -0.637 |
| | Inferior parietal | -0.630 | 0.523 |
| | Inferior temporal | 0.369 | -0.480 |
| | Insula | 0.694 | -0.626 |
| | Isthmus cingulate | -0.510 | -0.061 |
| | Lateral occipital | -0.912 | 1.206 |
| | Lateral orbitofrontal | 0.468 | -0.089 |
| | Lingual | -1.638 | 1.885 |
| | Medial orbitofrontal | 0.938 | -0.730 |
| | Middle temporal | 0.115 | -0.497 |
| | Paracentral | -0.554 | 0.158 |
| | Parahippocampal | 1.111 | -1.081 |
| | Pars opercularis | 0.483 | -0.346 |
| | Pars orbitalis | -0.154 | -0.508 |
| | Pars triangularis | 0.871 | 0.452 |
| | Pericalcarine | -2.480 | 2.434 |
| | Postcentral | -1.274 | 0.366 |
| | Posterior cingulate | 0.381 | -0.144 |
| | Precentral | -0.510 | -0.044 |
| | Precuneus | -0.792 | 0.681 |
| | Rostral anterior cingulate | 1.372 | -1.379 |
| | Rostral middle frontal | 0.407 | 0.116 |
| | Superior frontal | 0.454 | -0.205 |
| | Superior parietal | -0.826 | 0.369 |
| | Superior temporal | 0.169 | -0.340 |
| | Supramarginal | -0.676 | -0.417 |
| | Temporal pole | 1.514 | -1.558 |
| | Transverse temporal | -0.869 | 0.919 |
| Right | Bank ssts | -1.996 | -1.592 |
| | Caudal anterior cingulate | 0.799 | -1.035 |
| | Caudal middle frontal | 0.449 | 0.360 |
| | Cuneus | -1.815 | 2.416 |
| | Entorhinal | 1.501 | -1.713 |
| | Frontal pole | -0.361 | -1.312 |
| | Fusiform | 0.444 | -0.260 |
| | Inferior parietal | -0.647 | 0.145 |
| | Inferior temporal | 0.758 | -0.090 |
| | Insula | 0.972 | -0.632 |
| | Isthmus cingulate | -0.150 | 0.639 |
| | Lateral occipital | -0.770 | 1.232 |
| | Lateral orbitofrontal | 0.691 | -0.313 |
| | Lingual | -1.992 | 0.746 |
| | Medial orbitofrontal | 0.844 | -0.592 |
| | Middle temporal | 0.589 | -0.080 |
| | Paracentral | -0.990 | -0.139 |
| | Parahippocampal | 1.342 | -0.454 |
| | Pars opercularis | 0.030 | -0.994 |
| | Pars orbitalis | 0.475 | 0.729 |
| | Pars triangularis | 0.918 | 0.440 |
| | Pericalcarine | -2.188 | 2.730 |
| | Postcentral | -0.944 | 0.674 |
| | Posterior cingulate | 0.764 | -0.117 |
| | Precentral | -0.404 | 0.439 |
| | Precuneus | -0.306 | 0.560 |
| | Rostral anterior cingulate | 1.026 | -1.221 |
| | Rostral middle frontal | 0.286 | 0.189 |
| | Superior frontal | 0.582 | -0.279 |
| | Superior parietal | -0.184 | 0.810 |
| | Superior temporal | 0.598 | 0.291 |
| | Supramarginal | -1.050 | -0.356 |
| | Temporal pole | 0.930 | -1.547 |
| | Transverse temporal | -0.202 | 0.328 |

*Table S11* Brief descriptions of UKB cognitive tests and index codes.

| Cognitive Test | Brief description | UKB field ID |
|---|---|---|
| Reaction time (s) | Time taken to respond in snap-type computer game | 20023 |
| Number span | Number of rounds completed (the maximum length of number string recalled) | 4282 |
| Fluid intelligence | Number of 13 verbal and numerical logic questions correct | 20016 |
| Trail making B (s) [4] | Time taken to complete an alphanumeric path-making test (trail B) | 6350 |
| Matrix pattern (log) [5] | Number of matrix pattern puzzles solved | 6373 |
| Tower task | Number of tower task puzzles solved | 21003 |
| Digit-symbol substitution [6] | Number of digit-symbol pairs matched | 23324 |
| Pairs matching [7] | Number of incorrect matches in a 6-pair classic pairs game | 399 |
| Prospective memory | Test of instruction recall after a delay | 20018 |
| Paired associates | Number of novel word pairs correctly recalled after a delay | 20197 |
| Picture vocabulary | Estimate of cognitive ability based on picture vocabulary task | 26302 |

*Table S12* Brief descriptions of STRADL cognitive tests and index codes.

| Cognitive Test | Brief description | Code |
|---|---|---|
| Matrix reasoning ([5]) | Number of puzzles correct | mrtotc |
| Verbal fluency ([8]) | Number of words recalled beginning with C, F and L in 1 minute (1 minute per letter) | vftot |
| Mill Hill vocabulary ([9]) | Number of word meanings explained correctly | mhv |
| Digit symbol substitution ([10]) | Number of digit-symbol pairs matched | digsym |
| Logical memory ([10]) | Story recall score (total from immediate and delayed tests) | mema + medela |

*Table S13* Brief descriptions of LBC1936 cognitive tests and index codes.

| Cognitive test | Brief description | Code |
|---|---|---|
| Matrix reasoning ([10]) | Number of puzzles correct | matreas_w2 |
| Block design ([10]) | Number of puzzles correct | blkdes_w2 |
| Spatial span ([11]) | Number of block sequences correct (forwards + backwards) | spantot_w2 |
| National adult reading test (NART) ([12]) | Number of words from the list pronounced correctly | nart_w2 |
| Wechsler Test of Adult reading (WTAR) ([13]) | Number of words from the list pronounced correctly | wtar_w2 |
| Verbal fluency ([14]) | Number of words recalled beginning with C, F and L i in 1 minute (1 minute per letter) | vftot_w2 |
| Verbal paired associates ([11]) | Number of novel word pairs recalled after a delay (total from immediate and delayed tests) | vpatotal_w2 |
| Logical memory ([11]) | Number of story details recalled (out of a total possible of 25) - total from immediate and delayed tests | lmtotal_w2 |
| Digit span backwards ([10]) | Max length of a string of numbers recalled in reverse | digback_w2 |
| Symbol search | Number of symbols correctly detected during visual search | symsear_w2 |
| Digit-symbol substitution ([15]) | Number of digit-symbol pairs matched | digsym_w2 |
| Inspection time ([16]) | Number of correct responses at visual selection of longest line (two-alternative forced choice) | Ittotal_w2 |
| Four-choice reaction time (s) ([17]) | Time taken to press the indicated button (out of 4 buttons) | crtmean_w2 |

*Table S14* UKB cognitive test summary statistics, and latent cognitive ability model estimates (for all paths to the latent factor, $p < .001$).

| Cognitive Test | *N* | *M* (*SD*) | β | Residual variance |
|---|---|---|---|---|
| Reaction time (log) | 35138 | 6.38 (0.17) | 0.22 (0.006) | 0.84 |
| Numeric memory | 25720 | 6.76 (1.27) | -0.46 (0.006) | 0.76 |
| Fluid intelligence | 34709 | 6.60 (2.05) | -0.70 (0.004) | 0.49 |
| Trail making B (log) | 24484 | 6.28 (0.36) | 0.59 (0.005) | 0.49 |
| Matrix pattern | 25135 | 7.95 (2.14) | -0.58 (0.005) | 0.60 |
| Tower task | 24909 | 9.86 (3.23) | -0.49 (0.006) | 0.71 |
| Digit-symbol substitution | 25134 | 18.87 (5.27) | -0.44 (0.005) | 0.62 |
| Pairs matching (log) | 35365 | 1.35 (0.63) | 0.24 (0.006) | 0.92 |
| Prospective memory | 35350 | 0.84 (0.37) | 0.31 (0.006) | 0.88 |
| Paired associates | 25404 | 7.90 (02.64) | -0.44 (0.006) | 0.75 |
| Picture vocabulary | 25077 | | -0.48 (0.006) | 0.75 |

*Table S15* STRADL cognitive test summary statistics and latent cognitive ability model estimates (for all paths to the latent factor, $p < .001$).

| Cognitive Test | *N* | *M* (*SD*) | β (*SE*) | Residual variance |
|---|---|---|---|---|
| Matrix reasoning | 1043 | 8.30 (2.39) | 0.56 (0.029) | 0.65 |
| Verbal fluency | 1043 | 43.10 (11.92) | 0.53 (0.030) | 0.71 |
| Mill Hill vocabulary | 1043 | 31.64 (4.07) | 0.70 (0.028) | 0.45 |
| Digit symbol substitution | 1043 | 68.77 (15.13) | 0.37 (0.039) | 0.64 |
| Logical memory | 1043 | 31.91 (7.23) | 0.47 (0.030) | 0.72 |

*Table S16* LBC1936 cognitive test summary statistics, and cognitive ability model estimates (for all paths, besides the path between Verbal Memory and Cognitive ability which was fixed, all $p$ < .001).

| Cognitive test | *N* | *M (SD)* | β (*SE*) | Residual variance |
|---|---|---|---|---|
| Matrix reasoning | 634 | 13.52 (4.93) | 0.60 (0.03) | 0.62 |
| Block design | 634 | 34.38 (10.01) | 0.60 (0.03) | 0.60 |
| Spatial span | 634 | 14.79 (2,72) | 0.45 (0.04) | 0.77 |
| NART | 634 | 34.66 (8.10) | 0.63 (0.03) | 0.57 |
| WTAR | 634 | 41.27 (6.94) | 0.65 (0.03) | 0.56 |
| Phonemic verbal fluency | 635 | 43.55 (12.78) | 0.47 (0.04) | 0.77 |
| Verbal paired associates | 623 | 27.57 (9.48) | 0.51 (0.04) | 0.70 |
| Logical memory | 635 | 75.03 (17.84) | 0.52 (0.04) | 0.71 |
| Digit span backwards | 636 | 7.88 (2.31) | 0.55 (0.04) | 0.69 |
| Symbol search | 634 | 24.88 (6.05) | 0.58 (0.03) | 0.64 |
| Digit-symbol substitution | 634 | 56.68 (11.79) | 0.61 (0.03) | 0.57 |
| Inspection time | 634 | 111.78 (10.95) | 0.38 (0.04) | 0.84 |
| Four-choice reaction time (s) | 635 | 0.64 (0.08) | 0.39 (0.04) | 0.82 |

*Table S17 W*ithin-domain residual variances for LBC1936 general cognitive ability model.

| Cognitive test | β (*SE*) |
|---|---|
| Matrix reasoning ~~ block design | 0.25 (0.05) |
| Matrix reasoning ~~ spatial span | 0.09 (0.05) |
| Block design ~~ spatial span | 0.20 (0.05) |
| NART ~~ WTAR | 0.83 (0.02) |
| NART~~ verbal fluency | 0.16 (0.05) |
| WTAR ~~ verbal fluency | 0.16 (0.05) |
| Verbal paired associates ~~ logical memory | 0.35 (0.04) |
| Verbal paired associates ~~ digit span backward | -0.04 (0.05) |
| Logical memory ~~ digit span backward | 0.01 (0.05) |
| Symbol search ~~ digit symbol | 0.40 (0.04) |
| Symbol search ~~ inspection time | 0.17 (0.04) |
| Symbol search ~~ choice reaction time | 0.31 (0.04) |
| Digit symbol ~~ inspection time | 0.22 (0.04) |
| Digit symbol ~~ choice reaction time | 0.36 (0.04) |
| Inspection time ~~ choice reaction time | 0.25 (0.04) |

*Table S18* Model fits for the latent cognitive ability models*.*

| Cohort | $X^2$ | *df* | CFI | TLI | RMSEA | SRMR |
|---|---|---|---|---|---|---|
| UKB | 3309 | 44 | 0.949 | 0.911 | 0.044 | 0.025 |
| STRADL | 1170 | 20 | 0.972 | 0.887 | 0.079 | 0.019 |
| LBC1936 | 168 | 35 | 0.966 | 0.930 | 0.061 | 0.037 |

*Table S19* Meta-analysis results for regional *g*-**volume** associations.

| Volume | Left hemisphere | | | | | | | | Right hemisphere | | | | | | | |
|---|---|---|---|---|---|---|---|---|---|---|---|---|---|---|---|---|
| Region | B | *SE* | *p* | FDR Q | Cochrane's *Q* | *p* (*Q*) | I^2 | H^2 | B | *SE* | *p* | FDR Q | Cochrane's *Q* | *p* (*Q*) | I^2 | H^2 |
| Bank ssts | 0.110 | 0.031 | 0.000 | 0.001 | 7.785 | 0.020 | 0.002 | 5.082 | 0.111 | 0.024 | 0.000 | 0.000 | 5.993 | 0.050 | 0.001 | 3.272 |
| Caudal anterior cingulate | 0.045 | 0.005 | 0.000 | 0.000 | 0.607 | 0.738 | 0.000 | 1.000 | 0.041 | 0.005 | 0.000 | 0.000 | 0.533 | 0.766 | 0.000 | 1.000 |
| Caudal middle frontal | 0.107 | 0.025 | 0.000 | 0.000 | 5.673 | 0.059 | 0.001 | 3.425 | 0.091 | 0.009 | 0.000 | 0.000 | 2.101 | 0.350 | 0.000 | 1.125 |
| Cuneus | 0.062 | 0.005 | 0.000 | 0.000 | 0.415 | 0.812 | 0.000 | 1.000 | 0.054 | 0.033 | 0.096 | 0.098 | 7.231 | 0.027 | 0.003 | 5.842 |
| Entorhinal | 0.103 | 0.026 | 0.000 | 0.000 | 8.610 | 0.014 | 0.001 | 3.807 | 0.090 | 0.026 | 0.000 | 0.001 | 7.858 | 0.020 | 0.001 | 3.720 |
| Frontal pole | 0.048 | 0.005 | 0.000 | 0.000 | 2.338 | 0.311 | 0.000 | 1.002 | 0.016 | 0.015 | 0.301 | 0.301 | 3.014 | 0.222 | 0.000 | 1.569 |
| Fusiform | 0.158 | 0.039 | 0.000 | 0.000 | 17.520 | 0.000 | 0.004 | 9.872 | 0.154 | 0.037 | 0.000 | 0.000 | 18.194 | 0.000 | 0.003 | 8.804 |
| Inferior parietal | 0.141 | 0.031 | 0.000 | 0.000 | 11.869 | 0.003 | 0.002 | 5.418 | 0.164 | 0.041 | 0.000 | 0.000 | 22.780 | 0.000 | 0.004 | 11.305 |
| Inferior temporal | 0.137 | 0.049 | 0.005 | 0.005 | 17.043 | 0.000 | 0.006 | 14.800 | 0.143 | 0.053 | 0.007 | 0.007 | 20.177 | 0.000 | 0.008 | 18.648 |
| Insula | 0.116 | 0.004 | 0.000 | < .001 | 1.015 | 0.602 | < .001 | 1.000 | 0.123 | 0.027 | < .001 | < .001 | 8.333 | 0.016 | 0.002 | 4.978 |
| Isthmus cingulate | 0.085 | 0.012 | < .001 | < .001 | 2.575 | 0.276 | < .001 | 1.398 | 0.068 | 0.005 | < .001 | < .001 | 0.454 | 0.797 | < .001 | 1.000 |
| Lateral occipital | 0.085 | 0.004 | < .001 | < .001 | 1.009 | 0.604 | < .001 | 1.000 | 0.087 | 0.010 | < .001 | < .001 | 2.808 | 0.246 | < .001 | 1.221 |
| Lateral orbitofrontal | 0.128 | 0.017 | < .001 | < .001 | 4.094 | 0.129 | < .001 | 2.064 | 0.109 | 0.005 | < .001 | < .001 | 2.752 | 0.253 | < .001 | 1.004 |
| Lingual | 0.092 | 0.034 | 0.007 | 0.008 | 9.675 | 0.008 | 0.003 | 6.629 | 0.120 | 0.044 | 0.006 | 0.006 | 20.018 | < .001 | 0.005 | 11.091 |
| Medial orbitofrontal | 0.089 | 0.005 | < .001 | < .001 | 0.110 | 0.947 | < .001 | 1.000 | 0.094 | 0.004 | < .001 | < .001 | 0.954 | 0.621 | < .001 | 1.000 |
| Middle temporal | 0.156 | 0.045 | 0.001 | 0.001 | 17.181 | < .001 | 0.006 | 13.446 | 0.156 | 0.034 | < .001 | < .001 | 13.277 | 0.001 | 0.003 | 7.964 |
| Paracentral | 0.090 | 0.015 | < .001 | < .001 | 2.744 | 0.254 | < .001 | 1.569 | 0.083 | 0.005 | < .001 | < .001 | 0.717 | 0.699 | < .001 | 1.000 |
| Parahippocampal | 0.061 | 0.010 | < .001 | < .001 | 1.928 | 0.381 | < .001 | 1.172 | 0.106 | 0.036 | 0.003 | 0.004 | 14.066 | 0.001 | 0.003 | 7.173 |
| Pars opercularis | 0.110 | 0.025 | < .001 | < .001 | 6.645 | 0.036 | 0.001 | 3.649 | 0.074 | 0.005 | < .001 | < .001 | 1.000 | 0.607 | < .001 | 1.000 |
| Pars orbitalis | 0.089 | 0.005 | < .001 | < .001 | 0.255 | 0.880 | < .001 | 1.000 | 0.076 | 0.018 | < .001 | < .001 | 4.025 | 0.134 | 0.001 | 2.013 |
| Pars triangularis | 0.087 | 0.005 | < .001 | < .001 | 3.070 | 0.215 | < .001 | 1.015 | 0.073 | 0.005 | < .001 | < .001 | 0.571 | 0.752 | < .001 | 1.000 |
| Pericalcarine | 0.054 | 0.005 | < .001 | < .001 | 1.276 | 0.528 | < .001 | 1.000 | 0.065 | 0.017 | < .001 | < .001 | 3.249 | 0.197 | < .001 | 1.761 |
| Postcentral | 0.115 | 0.027 | < .001 | < .001 | 6.100 | 0.047 | 0.002 | 4.297 | 0.113 | 0.020 | < .001 | < .001 | 5.208 | 0.074 | 0.001 | 2.616 |
| Posterior cingulate | 0.085 | 0.005 | < .001 | < .001 | 1.359 | 0.507 | < .001 | 1.000 | 0.087 | 0.005 | < .001 | < .001 | 4.095 | 0.129 | < .001 | 1.000 |
| Precentral | 0.135 | 0.022 | < .001 | < .001 | 5.651 | 0.059 | 0.001 | 3.012 | 0.150 | 0.032 | < .001 | < .001 | 11.780 | 0.003 | 0.003 | 6.422 |
| Precuneus | 0.131 | 0.025 | < .001 | < .001 | 8.113 | 0.017 | 0.001 | 4.113 | 0.142 | 0.029 | < .001 | < .001 | 11.525 | 0.003 | 0.002 | 5.587 |
| Rostral anterior cingulate | 0.095 | 0.017 | < .001 | < .001 | 3.741 | 0.154 | < .001 | 1.816 | 0.082 | 0.016 | < .001 | < .001 | 3.020 | 0.221 | < .001 | 1.681 |
| Rostral middle frontal | 0.121 | 0.017 | < .001 | < .001 | 4.228 | 0.121 | < .001 | 2.110 | 0.119 | 0.018 | < .001 | < .001 | 4.610 | 0.100 | 0.001 | 2.289 |
| Superior frontal | 0.126 | 0.019 | < .001 | < .001 | 5.294 | 0.071 | 0.001 | 2.720 | 0.138 | 0.026 | < .001 | < .001 | 10.847 | 0.004 | 0.001 | 4.487 |
| Superior parietal | 0.116 | 0.028 | < .001 | < .001 | 7.174 | 0.028 | 0.002 | 4.684 | 0.120 | 0.032 | < .001 | < .001 | 9.309 | 0.010 | 0.002 | 6.162 |
| Superior temporal | 0.175 | 0.033 | < .001 | < .001 | 17.090 | < .001 | 0.003 | 7.183 | 0.164 | 0.033 | < .001 | < .001 | 11.702 | 0.003 | 0.003 | 6.736 |
| Supramarginal | 0.115 | 0.018 | < .001 | < .001 | 4.279 | 0.118 | 0.001 | 2.150 | 0.105 | 0.023 | < .001 | < .001 | 5.156 | 0.076 | 0.001 | 3.283 |
| Temporal pole | 0.069 | 0.021 | 0.001 | 0.001 | 5.042 | 0.080 | 0.001 | 2.424 | 0.039 | 0.005 | < .001 | < .001 | 0.133 | 0.936 | < .001 | 1.000 |
| Transverse temporal | 0.083 | 0.005 | < .001 | < .001 | 0.705 | 0.703 | < .001 | 1.000 | 0.108 | 0.022 | < .001 | < .001 | 4.992 | 0.082 | 0.001 | 2.531 |

*Table S20* Meta-analysis results for regional *g*-**surface area** associations.

| Volume | Left hemisphere | | | | | | | | Right hemisphere | | | | | | | |
|---|---|---|---|---|---|---|---|---|---|---|---|---|---|---|---|---|
| Region | B | *SE* | *p* | FDR Q | Cochrane's *Q* | *p* (*Q*) | I^2 | H^2 | B | *SE* | *p* | FDR Q | Cochrane's *Q* | *p* (*Q*) | I^2 | H^2 |
| Bank ssts | 0.090 | 0.015 | < .001 | < .001 | 3.525 | 0.172 | < .001 | 1.534 | 0.115 | 0.021 | < .001 | < .001 | 5.052 | 0.080 | 0.001 | 2.530 |
| Caudal anterior cingulate | 0.069 | 0.006 | < .001 | < .001 | 1.356 | 0.508 | < .001 | 1.015 | 0.079 | 0.013 | < .001 | < .001 | 2.704 | 0.259 | < .001 | 1.380 |
| Caudal middle frontal | 0.081 | 0.005 | < .001 | < .001 | 1.097 | 0.578 | < .001 | 1.000 | 0.089 | 0.005 | < .001 | < .001 | 1.354 | 0.508 | < .001 | 1.000 |
| Cuneus | 0.074 | 0.011 | < .001 | < .001 | 2.329 | 0.312 | < .001 | 1.244 | 0.078 | 0.005 | < .001 | < .001 | 3.242 | 0.198 | < .001 | 1.003 |
| Entorhinal | 0.080 | 0.016 | < .001 | < .001 | 3.189 | 0.203 | < .001 | 1.686 | 0.062 | 0.005 | < .001 | < .001 | 0.162 | 0.922 | < .001 | 1.000 |
| Frontal pole | 0.086 | 0.005 | < .001 | < .001 | 3.103 | 0.212 | < .001 | 1.005 | 0.060 | 0.015 | < .001 | < .001 | 3.045 | 0.218 | < .001 | 1.541 |
| Fusiform | 0.141 | 0.032 | < .001 | < .001 | 9.836 | 0.007 | 0.002 | 6.085 | 0.126 | 0.021 | < .001 | < .001 | 5.943 | 0.051 | 0.001 | 2.831 |
| Inferior parietal | 0.139 | 0.028 | < .001 | < .001 | 10.734 | 0.005 | 0.002 | 4.420 | 0.150 | 0.031 | < .001 | < .001 | 15.616 | < .001 | 0.002 | 6.053 |
| Inferior temporal | 0.147 | 0.042 | < .001 | 0.001 | 14.603 | 0.001 | 0.005 | 10.963 | 0.147 | 0.039 | < .001 | < .001 | 13.368 | 0.001 | 0.004 | 9.340 |
| Insula | 0.100 | 0.004 | < .001 | < .001 | 1.193 | 0.551 | < .001 | 1.000 | 0.108 | 0.015 | < .001 | < .001 | 3.704 | 0.157 | < .001 | 1.727 |
| Isthmus cingulate | 0.082 | 0.004 | < .001 | < .001 | 0.932 | 0.627 | < .001 | 1.000 | 0.072 | 0.005 | < .001 | < .001 | 0.500 | 0.779 | < .001 | 1.000 |
| Lateral occipital | 0.092 | 0.004 | < .001 | < .001 | 1.548 | 0.461 | < .001 | 1.000 | 0.090 | 0.004 | < .001 | < .001 | 0.755 | 0.685 | < .001 | 1.000 |
| Lateral orbitofrontal | 0.138 | 0.019 | < .001 | < .001 | 4.914 | 0.086 | 0.001 | 2.382 | 0.125 | 0.028 | < .001 | < .001 | 6.821 | 0.033 | 0.002 | 4.492 |
| Lingual | 0.098 | 0.041 | 0.016 | 0.017 | 11.350 | 0.003 | 0.004 | 9.288 | 0.111 | 0.035 | 0.002 | 0.002 | 10.039 | 0.007 | 0.003 | 6.760 |
| Medial orbitofrontal | 0.097 | 0.004 | < .001 | < .001 | 1.291 | 0.524 | < .001 | 1.000 | 0.111 | 0.004 | < .001 | < .001 | 2.534 | 0.282 | < .001 | 1.000 |
| Middle temporal | 0.150 | 0.029 | < .001 | < .001 | 9.055 | 0.011 | 0.002 | 5.046 | 0.148 | 0.024 | < .001 | < .001 | 6.937 | 0.031 | 0.001 | 3.508 |
| Paracentral | 0.079 | 0.013 | < .001 | < .001 | 2.262 | 0.323 | < .001 | 1.400 | 0.083 | 0.005 | < .001 | < .001 | 1.754 | 0.416 | < .001 | 1.001 |
| Parahippocampal | 0.066 | 0.013 | < .001 | < .001 | 3.094 | 0.213 | < .001 | 1.363 | 0.071 | 0.005 | < .001 | < .001 | 1.037 | 0.595 | < .001 | 1.000 |
| Pars opercularis | 0.113 | 0.028 | < .001 | < .001 | 8.319 | 0.016 | 0.002 | 4.180 | 0.077 | 0.005 | < .001 | < .001 | 0.524 | 0.769 | < .001 | 1.000 |
| Pars orbitalis | 0.099 | 0.005 | < .001 | < .001 | 1.983 | 0.371 | < .001 | 1.004 | 0.098 | 0.005 | < .001 | < .001 | 1.359 | 0.507 | < .001 | 1.011 |
| Pars triangularis | 0.091 | 0.005 | < .001 | < .001 | 1.593 | 0.451 | < .001 | 1.002 | 0.079 | 0.005 | < .001 | < .001 | 1.309 | 0.520 | < .001 | 1.006 |
| Pericalcarine | 0.066 | 0.010 | < .001 | < .001 | 2.629 | 0.269 | < .001 | 1.161 | 0.083 | 0.018 | < .001 | < .001 | 3.929 | 0.140 | 0.001 | 2.000 |
| Postcentral | 0.095 | 0.004 | < .001 | < .001 | 0.648 | 0.723 | < .001 | 1.000 | 0.105 | 0.015 | < .001 | < .001 | 2.970 | 0.227 | < .001 | 1.662 |
| Posterior cingulate | 0.093 | 0.005 | < .001 | < .001 | 0.564 | 0.754 | < .001 | 1.000 | 0.111 | 0.023 | < .001 | < .001 | 5.171 | 0.075 | 0.001 | 3.076 |
| Precentral | 0.091 | 0.004 | < .001 | < .001 | 0.212 | 0.899 | < .001 | 1.000 | 0.094 | 0.004 | < .001 | < .001 | 0.513 | 0.774 | < .001 | 1.000 |
| Precuneus | 0.122 | 0.022 | < .001 | < .001 | 6.262 | 0.044 | 0.001 | 3.025 | 0.135 | 0.027 | < .001 | < .001 | 10.328 | 0.006 | 0.002 | 4.604 |
| Rostral anterior cingulate | 0.109 | 0.025 | < .001 | < .001 | 6.290 | 0.043 | 0.001 | 3.466 | 0.104 | 0.019 | < .001 | < .001 | 4.261 | 0.119 | 0.001 | 2.135 |
| Rostral middle frontal | 0.126 | 0.017 | < .001 | < .001 | 3.866 | 0.145 | < .001 | 1.971 | 0.130 | 0.019 | < .001 | < .001 | 4.860 | 0.088 | 0.001 | 2.378 |
| Superior frontal | 0.122 | 0.016 | < .001 | < .001 | 3.868 | 0.145 | < .001 | 1.971 | 0.147 | 0.027 | < .001 | < .001 | 11.637 | 0.003 | 0.002 | 4.824 |
| Superior parietal | 0.108 | 0.023 | < .001 | < .001 | 5.589 | 0.061 | 0.001 | 3.125 | 0.119 | 0.027 | < .001 | < .001 | 7.744 | 0.021 | 0.002 | 4.167 |
| Superior temporal | 0.150 | 0.023 | < .001 | < .001 | 7.701 | 0.021 | 0.001 | 3.362 | 0.122 | 0.004 | < .001 | < .001 | 1.091 | 0.580 | < .001 | 1.000 |
| Supramarginal | 0.107 | 0.018 | < .001 | < .001 | 4.157 | 0.125 | 0.001 | 2.103 | 0.088 | 0.005 | < .001 | < .001 | 1.539 | 0.463 | < .001 | 1.002 |
| Temporal pole | 0.074 | 0.016 | < .001 | < .001 | 2.992 | 0.224 | < .001 | 1.673 | 0.021 | 0.023 | 0.358 | 0.358 | 5.629 | 0.060 | 0.001 | 2.634 |
| Transverse temporal | 0.080 | 0.005 | < .001 | < .001 | 0.742 | 0.690 | < .001 | 1.000 | 0.087 | 0.007 | < .001 | < .001 | 1.815 | 0.404 | < .001 | 1.068 |

*Table S21* Meta-analysis results for regional *g*-**thickness** associations.

| Volume | Left hemisphere | | | | | | | | Right hemisphere | | | | | | | |
|---|---|---|---|---|---|---|---|---|---|---|---|---|---|---|---|---|
| Region | B | *SE* | *p* | FDR Q | Cochrane's *Q* | *p* (*Q*) | I^2 | H^2 | B | *SE* | *p* | FDR Q | Cochrane's *Q* | *p* (*Q*) | I^2 | H^2 |
| Bank ssts | 0.043 | 0.036 | 0.241 | 0.299 | 8.411 | 0.015 | 0.003 | 6.538 | 0.052 | 0.044 | 0.233 | 0.299 | 13.068 | 0.001 | 0.005 | 10.048 |
| Caudal anterior cingulate | -0.037 | 0.022 | 0.087 | 0.173 | 4.902 | 0.086 | 0.001 | 2.374 | -0.048 | 0.016 | 0.002 | 0.022 | 2.644 | 0.267 | < .001 | 1.533 |
| Caudal middle frontal | 0.082 | 0.042 | 0.049 | 0.128 | 14.837 | 0.001 | 0.004 | 9.711 | 0.043 | 0.020 | 0.035 | 0.107 | 4.783 | 0.091 | 0.001 | 2.372 |
| Cuneus | 0.012 | 0.005 | 0.018 | 0.090 | 0.027 | 0.987 | < .001 | 1.000 | 0.006 | 0.005 | 0.233 | 0.299 | 3.003 | 0.223 | < .001 | 1.000 |
| Entorhinal | 0.044 | 0.025 | 0.082 | 0.173 | 5.366 | 0.068 | 0.001 | 3.071 | 0.084 | 0.054 | 0.123 | 0.197 | 25.019 | < .001 | 0.008 | 14.418 |
| Frontal pole | -0.003 | 0.005 | 0.514 | 0.546 | 0.640 | 0.726 | < .001 | 1.000 | -0.018 | 0.005 | < .001 | 0.006 | 0.203 | 0.903 | < .001 | 1.000 |
| Fusiform | 0.071 | 0.035 | 0.043 | 0.123 | 12.092 | 0.002 | 0.003 | 6.345 | 0.082 | 0.047 | 0.085 | 0.173 | 21.225 | < .001 | 0.006 | 11.839 |
| Inferior parietal | 0.029 | 0.027 | 0.280 | 0.334 | 5.814 | 0.055 | 0.002 | 4.063 | 0.051 | 0.048 | 0.285 | 0.334 | 14.151 | 0.001 | 0.006 | 14.709 |
| Inferior temporal | 0.022 | 0.029 | 0.450 | 0.486 | 6.160 | 0.046 | 0.002 | 4.677 | 0.050 | 0.052 | 0.343 | 0.382 | 16.035 | < .001 | 0.008 | 15.565 |
| Insula | 0.039 | 0.013 | 0.002 | 0.022 | 2.278 | 0.320 | < .001 | 1.367 | 0.028 | 0.021 | 0.172 | 0.249 | 4.848 | 0.089 | 0.001 | 2.364 |
| Isthmus cingulate | -0.019 | 0.005 | < .001 | 0.004 | 2.659 | 0.265 | < .001 | 1.002 | -0.009 | 0.005 | 0.085 | 0.173 | 3.710 | 0.156 | < .001 | 1.002 |
| Lateral occipital | 0.008 | 0.005 | 0.127 | 0.197 | 0.200 | 0.905 | < .001 | 1.000 | 0.026 | 0.027 | 0.324 | 0.367 | 5.861 | 0.053 | 0.002 | 4.114 |
| Lateral orbitofrontal | 0.011 | 0.005 | 0.032 | 0.107 | 0.412 | 0.814 | < .001 | 1.000 | 0.008 | 0.005 | 0.126 | 0.197 | 0.851 | 0.654 | < .001 | 1.000 |
| Lingual | 0.020 | 0.005 | < .001 | 0.002 | 1.523 | 0.467 | < .001 | 1.000 | 0.058 | 0.035 | 0.098 | 0.185 | 13.196 | 0.001 | 0.003 | 6.253 |
| Medial orbitofrontal | -0.003 | 0.005 | 0.537 | 0.561 | 1.272 | 0.529 | < .001 | 1.000 | -0.013 | 0.005 | 0.008 | 0.055 | 0.735 | 0.692 | < .001 | 1.000 |
| Middle temporal | 0.052 | 0.048 | 0.278 | 0.334 | 13.709 | 0.001 | 0.006 | 13.324 | 0.067 | 0.048 | 0.162 | 0.245 | 16.539 | < .001 | 0.006 | 13.519 |
| Paracentral | 0.036 | 0.019 | 0.061 | 0.144 | 4.310 | 0.116 | 0.001 | 2.172 | 0.023 | 0.017 | 0.182 | 0.258 | 3.801 | 0.149 | < .001 | 1.991 |
| Parahippocampal | 0.036 | 0.022 | 0.104 | 0.190 | 5.307 | 0.070 | 0.001 | 2.526 | 0.045 | 0.036 | 0.207 | 0.282 | 10.369 | 0.006 | 0.003 | 6.277 |
| Pars opercularis | 0.032 | 0.005 | < .001 | < .001 | 2.651 | 0.266 | < .001 | 1.006 | 0.043 | 0.020 | 0.032 | 0.107 | 4.642 | 0.098 | 0.001 | 2.298 |
| Pars orbitalis | 0.031 | 0.023 | 0.169 | 0.249 | 5.066 | 0.079 | 0.001 | 2.780 | 0.008 | 0.005 | 0.126 | 0.197 | 1.088 | 0.581 | < .001 | 1.000 |
| Pars triangularis | 0.035 | 0.029 | 0.220 | 0.293 | 6.337 | 0.042 | 0.002 | 4.359 | 0.012 | 0.015 | 0.417 | 0.457 | 3.092 | 0.213 | < .001 | 1.508 |
| Pericalcarine | -0.010 | 0.026 | 0.708 | 0.720 | 5.266 | 0.072 | 0.001 | 3.457 | -0.012 | 0.005 | 0.016 | 0.090 | 0.966 | 0.617 | < .001 | 1.000 |
| Postcentral | 0.062 | 0.038 | 0.106 | 0.190 | 9.713 | 0.008 | 0.004 | 9.042 | 0.035 | 0.015 | 0.018 | 0.090 | 3.605 | 0.165 | < .001 | 1.666 |
| Posterior cingulate | -0.003 | 0.032 | 0.921 | 0.921 | 6.524 | 0.038 | 0.002 | 4.945 | -0.017 | 0.009 | 0.063 | 0.144 | 2.087 | 0.352 | < .001 | 1.128 |
| Precentral | 0.099 | 0.044 | 0.024 | 0.107 | 16.601 | < .001 | 0.005 | 10.861 | 0.095 | 0.046 | 0.040 | 0.118 | 19.609 | < .001 | 0.006 | 11.679 |
| Precuneus | 0.034 | 0.016 | 0.034 | 0.107 | 3.590 | 0.166 | < .001 | 1.745 | 0.045 | 0.027 | 0.096 | 0.185 | 6.706 | 0.035 | 0.002 | 4.258 |
| Rostral anterior cingulate | -0.016 | 0.005 | 0.001 | 0.018 | 1.078 | 0.583 | < .001 | 1.000 | -0.037 | 0.013 | 0.004 | 0.030 | 2.430 | 0.297 | < .001 | 1.296 |
| Rostral middle frontal | 0.016 | 0.005 | 0.003 | 0.023 | 2.093 | 0.351 | < .001 | 1.011 | 0.002 | 0.005 | 0.709 | 0.720 | 0.867 | 0.648 | < .001 | 1.000 |
| Superior frontal | 0.027 | 0.018 | 0.127 | 0.197 | 4.104 | 0.128 | 0.001 | 2.058 | 0.017 | 0.017 | 0.319 | 0.367 | 3.633 | 0.163 | < .001 | 1.834 |
| Superior parietal | 0.028 | 0.015 | 0.059 | 0.144 | 3.187 | 0.203 | < .001 | 1.628 | 0.015 | 0.013 | 0.242 | 0.299 | 3.232 | 0.199 | < .001 | 1.460 |
| Superior temporal | 0.108 | 0.049 | 0.027 | 0.107 | 22.151 | < .001 | 0.006 | 13.875 | 0.124 | 0.067 | 0.064 | 0.144 | 36.726 | < .001 | 0.013 | 26.728 |
| Supramarginal | 0.043 | 0.034 | 0.204 | 0.282 | 7.622 | 0.022 | 0.003 | 6.520 | 0.064 | 0.032 | 0.046 | 0.126 | 9.204 | 0.010 | 0.002 | 6.020 |
| Temporal pole | 0.069 | 0.031 | 0.025 | 0.107 | 9.404 | 0.009 | 0.002 | 4.532 | 0.097 | 0.045 | 0.032 | 0.107 | 20.614 | < .001 | 0.005 | 9.896 |
| Transverse temporal | 0.022 | 0.010 | 0.018 | 0.090 | 1.643 | 0.440 | < .001 | 1.136 | 0.019 | 0.012 | 0.118 | 0.197 | 2.537 | 0.281 | < .001 | 1.286 |

*Table S22* Mean cohort **age moderation r**esults for meta-analysed ***g*-volume** associations.

| Volume | Left hemisphere | | | | | | | | Right hemisphere | | | | | | | |
|---|---|---|---|---|---|---|---|---|---|---|---|---|---|---|---|---|
| Region | B | *SE* | *p* | FDR Q | Cochrane's *Q* | *p* (*Q*) | I^2 | H^2 | B | *SE* | *p* | FDR Q | Cochrane's *Q* | *p* (*Q*) | I^2 | H^2 |
| Bank ssts | 0.008 | 0.005 | 0.126 | 0.613 | 4.094 | 0.043 | 0.001 | 4.094 | 0.006 | 0.005 | 0.275 | 0.825 | 4.372 | 0.037 | 0.001 | 4.372 |
| Caudal anterior cingulate | -0.003 | 0.004 | 0.469 | 0.825 | 0.083 | 0.773 | < .001 | 1.000 | < .001 | 0.004 | 0.979 | 0.980 | 0.533 | 0.466 | < .001 | 1.000 |
| Caudal middle frontal | 0.007 | 0.004 | 0.129 | 0.613 | 2.754 | 0.097 | 0.001 | 2.754 | 0.002 | 0.004 | 0.544 | 0.825 | 1.773 | 0.183 | < .001 | 1.773 |
| Cuneus | -0.002 | 0.004 | 0.660 | 0.859 | 0.222 | 0.637 | < .001 | 1.000 | -0.009 | 0.003 | 0.009 | 0.367 | 0.317 | 0.573 | < .001 | 1.000 |
| Entorhinal | 0.002 | 0.007 | 0.784 | 0.862 | 8.609 | 0.003 | 0.003 | 8.609 | 0.003 | 0.006 | 0.607 | 0.825 | 7.730 | 0.005 | 0.002 | 7.730 |
| Frontal pole | -0.006 | 0.004 | 0.131 | 0.613 | 0.054 | 0.817 | < .001 | 1.000 | -0.003 | 0.005 | 0.585 | 0.825 | 2.815 | 0.093 | 0.001 | 2.815 |
| Fusiform | 0.007 | 0.008 | 0.369 | 0.825 | 15.817 | < .001 | 0.005 | 15.817 | 0.005 | 0.008 | 0.527 | 0.825 | 17.784 | < .001 | 0.005 | 17.784 |
| Inferior parietal | 0.004 | 0.007 | 0.622 | 0.830 | 11.770 | 0.001 | 0.004 | 11.770 | 0.006 | 0.009 | 0.500 | 0.825 | 22.150 | < .001 | 0.006 | 22.150 |
| Inferior temporal | 0.012 | 0.006 | 0.029 | 0.561 | 6.430 | 0.011 | 0.002 | 6.430 | 0.013 | 0.006 | 0.042 | 0.561 | 9.455 | 0.002 | 0.003 | 9.455 |
| Insula | -0.002 | 0.003 | 0.571 | 0.825 | 0.695 | 0.405 | < .001 | 1.000 | -0.008 | 0.003 | 0.011 | 0.367 | 1.166 | 0.280 | < .001 | 1.166 |
| Isthmus cingulate | 0.002 | 0.004 | 0.602 | 0.825 | 2.345 | 0.126 | < .001 | 2.345 | 0.001 | 0.004 | 0.786 | 0.862 | 0.380 | 0.537 | < .001 | 1.000 |
| Lateral occipital | -0.003 | 0.003 | 0.317 | 0.825 | 0.006 | 0.939 | < .001 | 1.000 | 0.004 | 0.004 | 0.339 | 0.825 | 1.896 | 0.169 | < .001 | 1.896 |
| Lateral orbitofrontal | 0.002 | 0.005 | 0.600 | 0.825 | 3.976 | 0.046 | 0.001 | 3.976 | 0.005 | 0.003 | 0.116 | 0.613 | 0.282 | 0.595 | < .001 | 1.000 |
| Lingual | 0.008 | 0.006 | 0.135 | 0.613 | 5.634 | 0.018 | 0.002 | 5.634 | 0.008 | 0.009 | 0.374 | 0.825 | 17.998 | < .001 | 0.006 | 17.998 |
| Medial orbitofrontal | -0.001 | 0.003 | 0.822 | 0.874 | 0.059 | 0.808 | < .001 | 1.000 | 0.001 | 0.003 | 0.709 | 0.862 | 0.815 | 0.367 | < .001 | 1.000 |
| Middle temporal | 0.010 | 0.007 | 0.124 | 0.613 | 11.062 | 0.001 | 0.003 | 11.062 | 0.007 | 0.007 | 0.329 | 0.825 | 11.776 | 0.001 | 0.003 | 11.776 |
| Paracentral | -0.003 | 0.004 | 0.520 | 0.825 | 1.937 | 0.164 | < .001 | 1.937 | 0.001 | 0.004 | 0.760 | 0.862 | 0.624 | 0.430 | < .001 | 1.000 |
| Parahippocampal | 0.001 | 0.004 | 0.746 | 0.862 | 1.875 | 0.171 | < .001 | 1.875 | 0.006 | 0.008 | 0.452 | 0.825 | 13.141 | < .001 | 0.004 | 13.141 |
| Pars opercularis | 0.005 | 0.005 | 0.308 | 0.825 | 5.256 | 0.022 | 0.001 | 5.256 | 0.001 | 0.004 | 0.741 | 0.862 | 0.890 | 0.345 | < .001 | 1.000 |
| Pars orbitalis | 0.001 | 0.003 | 0.799 | 0.862 | 0.190 | 0.663 | < .001 | 1.000 | 0.006 | 0.004 | 0.134 | 0.613 | 1.310 | 0.252 | < .001 | 1.310 |
| Pars triangularis | 0.006 | 0.003 | 0.094 | 0.613 | 0.268 | 0.605 | < .001 | 1.000 | -0.003 | 0.004 | 0.450 | 0.825 | 0.001 | 0.971 | < .001 | 1.000 |
| Pericalcarine | -0.004 | 0.004 | 0.306 | 0.825 | 0.226 | 0.635 | < .001 | 1.000 | -0.002 | 0.004 | 0.590 | 0.825 | 2.520 | 0.112 | 0.001 | 2.520 |
| Postcentral | 0.008 | 0.004 | 0.048 | 0.561 | 1.720 | 0.190 | < .001 | 1.720 | 0.004 | 0.005 | 0.463 | 0.825 | 4.712 | 0.030 | 0.001 | 4.712 |
| Posterior cingulate | 0.003 | 0.004 | 0.473 | 0.825 | 0.845 | 0.358 | < .001 | 1.000 | 0.007 | 0.003 | 0.058 | 0.561 | 0.495 | 0.482 | < .001 | 1.000 |
| Precentral | 0.005 | 0.005 | 0.325 | 0.825 | 4.551 | 0.033 | 0.001 | 4.551 | 0.006 | 0.007 | 0.366 | 0.825 | 10.267 | 0.001 | 0.003 | 10.267 |
| Precuneus | 0.004 | 0.006 | 0.450 | 0.825 | 7.412 | 0.006 | 0.002 | 7.412 | 0.004 | 0.007 | 0.517 | 0.825 | 11.039 | 0.001 | 0.003 | 11.039 |
| Rostral anterior cingulate | 0.004 | 0.004 | 0.342 | 0.825 | 2.730 | 0.098 | 0.001 | 2.730 | < .001 | 0.005 | 0.980 | 0.980 | 2.962 | 0.085 | 0.001 | 2.962 |
| Rostral middle frontal | 0.003 | 0.004 | 0.480 | 0.825 | 3.868 | 0.049 | 0.001 | 3.868 | 0.002 | 0.005 | 0.669 | 0.859 | 4.547 | 0.033 | 0.001 | 4.547 |
| Superior frontal | 0.004 | 0.005 | 0.400 | 0.825 | 4.656 | 0.031 | 0.001 | 4.656 | -0.003 | 0.006 | 0.586 | 0.825 | 7.957 | 0.005 | 0.002 | 7.957 |
| Superior parietal | 0.007 | 0.005 | 0.156 | 0.662 | 4.298 | 0.038 | 0.001 | 4.298 | 0.007 | 0.006 | 0.194 | 0.778 | 6.523 | 0.011 | 0.002 | 6.523 |
| Superior temporal | 0.003 | 0.008 | 0.728 | 0.862 | 17.088 | < .001 | 0.005 | 17.088 | 0.006 | 0.006 | 0.319 | 0.825 | 9.840 | 0.002 | 0.003 | 9.840 |
| Supramarginal | 0.002 | 0.005 | 0.701 | 0.862 | 4.242 | 0.039 | 0.001 | 4.242 | 0.007 | 0.004 | 0.054 | 0.561 | 1.286 | 0.257 | < .001 | 1.286 |
| Temporal pole | -0.001 | 0.005 | 0.878 | 0.919 | 4.755 | 0.029 | 0.001 | 4.755 | -0.001 | 0.004 | 0.788 | 0.862 | 0.061 | 0.805 | < .001 | 1.000 |
| Transverse temporal | < .001 | 0.004 | 0.908 | 0.936 | 0.691 | 0.406 | < .001 | 1.000 | 0.005 | 0.005 | 0.365 | 0.825 | 3.933 | 0.047 | 0.001 | 3.933 |

*Table S23* Mean cohort **age moderation r**esults for meta-analysed ***g*-surface area** associations.

| Volume | Left hemisphere | | | | | | | | Right hemisphere | | | | | | | |
|---|---|---|---|---|---|---|---|---|---|---|---|---|---|---|---|---|
| Region | B | *SE* | *p* | FDR Q | Cochrane's *Q* | *p* (*Q*) | I^2 | H^2 | B | *SE* | *p* | FDR Q | Cochrane's *Q* | *p* (*Q*) | I^2 | H^2 |
| Bank ssts | 0.005 | 0.004 | 0.209 | 0.844 | 1.792 | 0.181 | < .001 | 1.792 | 0.004 | 0.005 | 0.446 | 0.924 | 4.382 | 0.036 | 0.001 | 4.382 |
| Caudal anterior cingulate | -0.001 | 0.004 | 0.741 | 0.954 | 1.216 | 0.270 | < .001 | 1.216 | 0.003 | 0.004 | 0.463 | 0.924 | 2.131 | 0.144 | < .001 | 2.131 |
| Caudal middle frontal | 0.003 | 0.004 | 0.400 | 0.924 | 0.390 | 0.532 | < .001 | 1.000 | 0.002 | 0.004 | 0.497 | 0.924 | 0.892 | 0.345 | < .001 | 1.000 |
| Cuneus | 0.002 | 0.004 | 0.550 | 0.924 | 2.013 | 0.156 | < .001 | 2.013 | -0.006 | 0.004 | 0.074 | 0.844 | 0.050 | 0.824 | < .001 | 1.000 |
| Entorhinal | -0.004 | 0.004 | 0.310 | 0.900 | 1.658 | 0.198 | < .001 | 1.658 | -0.001 | 0.004 | 0.700 | 0.933 | 0.013 | 0.909 | < .001 | 1.000 |
| Frontal pole | -0.006 | 0.004 | 0.120 | 0.844 | 0.691 | 0.406 | < .001 | 1.000 | -0.003 | 0.005 | 0.485 | 0.924 | 2.573 | 0.109 | 0.001 | 2.573 |
| Fusiform | 0.007 | 0.006 | 0.231 | 0.844 | 7.212 | 0.007 | 0.002 | 7.212 | 0.002 | 0.005 | 0.684 | 0.930 | 5.881 | 0.015 | 0.002 | 5.881 |
| Inferior parietal | < .001 | 0.007 | 0.992 | 0.992 | 10.393 | 0.001 | 0.003 | 10.393 | < .001 | 0.008 | 0.992 | 0.992 | 14.954 | < .001 | 0.004 | 14.954 |
| Inferior temporal | 0.010 | 0.006 | 0.080 | 0.844 | 7.277 | 0.007 | 0.002 | 7.277 | 0.009 | 0.006 | 0.146 | 0.844 | 8.416 | 0.004 | 0.002 | 8.416 |
| Insula | -0.002 | 0.003 | 0.484 | 0.924 | 0.703 | 0.402 | < .001 | 1.000 | -0.006 | 0.003 | 0.071 | 0.844 | 0.440 | 0.507 | < .001 | 1.000 |
| Isthmus cingulate | < .001 | 0.003 | 0.941 | 0.992 | 0.927 | 0.336 | < .001 | 1.000 | -0.002 | 0.003 | 0.626 | 0.930 | 0.263 | 0.608 | < .001 | 1.000 |
| Lateral occipital | -0.004 | 0.003 | 0.214 | 0.844 | 0.003 | 0.953 | < .001 | 1.000 | 0.001 | 0.003 | 0.772 | 0.955 | 0.672 | 0.413 | < .001 | 1.000 |
| Lateral orbitofrontal | < .001 | 0.005 | 0.973 | 0.992 | 4.755 | 0.029 | 0.001 | 4.755 | 0.007 | 0.004 | 0.090 | 0.844 | 2.876 | 0.090 | 0.001 | 2.876 |
| Lingual | 0.010 | 0.005 | 0.033 | 0.844 | 3.896 | 0.048 | 0.001 | 3.896 | 0.008 | 0.006 | 0.152 | 0.844 | 6.162 | 0.013 | 0.002 | 6.162 |
| Medial orbitofrontal | 0.001 | 0.004 | 0.744 | 0.954 | 1.200 | 0.273 | < .001 | 1.200 | 0.004 | 0.003 | 0.243 | 0.844 | 1.157 | 0.282 | < .001 | 1.157 |
| Middle temporal | 0.006 | 0.006 | 0.304 | 0.900 | 7.135 | 0.008 | 0.002 | 7.135 | 0.004 | 0.006 | 0.449 | 0.924 | 6.286 | 0.012 | 0.002 | 6.286 |
| Paracentral | -0.001 | 0.004 | 0.826 | 0.986 | 2.108 | 0.147 | < .001 | 2.108 | 0.003 | 0.004 | 0.344 | 0.900 | 0.858 | 0.354 | < .001 | 1.000 |
| Parahippocampal | 0.006 | 0.004 | 0.100 | 0.844 | 0.395 | 0.529 | < .001 | 1.000 | < .001 | 0.004 | 0.931 | 0.992 | 1.031 | 0.310 | < .001 | 1.031 |
| Pars opercularis | 0.005 | 0.006 | 0.430 | 0.924 | 7.283 | 0.007 | 0.002 | 7.283 | 0.002 | 0.004 | 0.571 | 0.924 | 0.203 | 0.652 | < .001 | 1.000 |
| Pars orbitalis | -0.005 | 0.003 | 0.159 | 0.844 | < .001 | 0.992 | < .001 | 1.000 | 0.004 | 0.003 | 0.244 | 0.844 | 0.003 | 0.956 | < .001 | 1.000 |
| Pars triangularis | 0.004 | 0.004 | 0.241 | 0.844 | 0.218 | 0.640 | < .001 | 1.000 | -0.004 | 0.004 | 0.264 | 0.855 | 0.062 | 0.803 | < .001 | 1.000 |
| Pericalcarine | 0.004 | 0.004 | 0.338 | 0.900 | 1.682 | 0.195 | < .001 | 1.682 | 0.002 | 0.005 | 0.658 | 0.930 | 3.804 | 0.051 | 0.001 | 3.804 |
| Postcentral | 0.002 | 0.003 | 0.562 | 0.924 | 0.311 | 0.577 | < .001 | 1.000 | < .001 | 0.004 | 0.936 | 0.992 | 2.963 | 0.085 | 0.001 | 2.963 |
| Posterior cingulate | 0.003 | 0.004 | 0.454 | 0.924 | 0.003 | 0.960 | < .001 | 1.000 | 0.007 | 0.004 | 0.062 | 0.844 | 1.401 | 0.236 | < .001 | 1.401 |
| Precentral | -0.002 | 0.003 | 0.646 | 0.930 | 0.001 | 0.972 | < .001 | 1.000 | < .001 | 0.003 | 0.990 | 0.992 | 0.513 | 0.474 | < .001 | 1.000 |
| Precuneus | 0.003 | 0.005 | 0.567 | 0.924 | 5.930 | 0.015 | 0.002 | 5.930 | 0.003 | 0.007 | 0.673 | 0.930 | 10.228 | 0.001 | 0.003 | 10.228 |
| Rostral anterior cingulate | 0.006 | 0.005 | 0.248 | 0.844 | 4.283 | 0.039 | 0.001 | 4.283 | 0.003 | 0.005 | 0.527 | 0.924 | 3.849 | 0.050 | 0.001 | 3.849 |
| Rostral middle frontal | 0.002 | 0.005 | 0.646 | 0.930 | 3.743 | 0.053 | 0.001 | 3.743 | 0.001 | 0.005 | 0.807 | 0.980 | 4.859 | 0.028 | 0.001 | 4.859 |
| Superior frontal | 0.002 | 0.004 | 0.640 | 0.930 | 3.729 | 0.053 | 0.001 | 3.729 | -0.004 | 0.006 | 0.569 | 0.924 | 8.889 | 0.003 | 0.002 | 8.889 |
| Superior parietal | 0.006 | 0.004 | 0.186 | 0.844 | 3.202 | 0.074 | 0.001 | 3.202 | 0.005 | 0.006 | 0.330 | 0.900 | 6.190 | 0.013 | 0.002 | 6.190 |
| Superior temporal | -0.001 | 0.006 | 0.909 | 0.992 | 7.326 | 0.007 | 0.002 | 7.326 | < .001 | 0.003 | 0.987 | 0.992 | 1.091 | 0.296 | < .001 | 1.091 |
| Supramarginal | -0.001 | 0.005 | 0.763 | 0.955 | 3.738 | 0.053 | 0.001 | 3.738 | 0.004 | 0.003 | 0.227 | 0.844 | 0.077 | 0.782 | < .001 | 1.000 |
| Temporal pole | -0.001 | 0.004 | 0.875 | 0.992 | 2.839 | 0.092 | 0.001 | 2.839 | 0.001 | 0.006 | 0.924 | 0.992 | 5.381 | 0.020 | 0.002 | 5.381 |
| Transverse temporal | 0.003 | 0.004 | 0.417 | 0.924 | 0.084 | 0.772 | < .001 | 1.000 | 0.002 | 0.004 | 0.627 | 0.930 | 1.604 | 0.205 | < .001 | 1.604 |

*Table S24* Mean cohort **age moderation r**esults for meta-analysed ***g*-thickness** associations.

| Volume | Left hemisphere | | | | | | | | Right hemisphere | | | | | | | |
|---|---|---|---|---|---|---|---|---|---|---|---|---|---|---|---|---|
| Region | B | *SE* | *p* | FDR Q | Cochrane's *Q* | *p* (*Q*) | I^2 | H^2 | B | *SE* | *p* | FDR Q | Cochrane's *Q* | *p* (*Q*) | I^2 | H^2 |
| Bank ssts | 0.010 | 0.005 | 0.037 | 0.277 | 2.596 | 0.107 | 0.001 | 2.596 | 0.011 | 0.006 | 0.088 | 0.438 | 6.875 | 0.009 | 0.002 | 6.875 |
| Caudal anterior cingulate | 0.001 | 0.006 | 0.909 | 0.936 | 4.674 | 0.031 | 0.002 | 4.674 | 0.002 | 0.005 | 0.700 | 0.823 | 2.285 | 0.131 | 0.001 | 2.285 |
| Caudal middle frontal | 0.009 | 0.007 | 0.246 | 0.619 | 11.898 | 0.001 | 0.004 | 11.898 | 0.003 | 0.005 | 0.615 | 0.804 | 4.673 | 0.031 | 0.001 | 4.673 |
| Cuneus | -0.001 | 0.004 | 0.874 | 0.936 | 0.001 | 0.970 | < .001 | 1.000 | -0.006 | 0.004 | 0.102 | 0.438 | 0.324 | 0.569 | < .001 | 1.000 |
| Entorhinal | 0.007 | 0.005 | 0.124 | 0.491 | 2.375 | 0.123 | 0.001 | 2.375 | 0.011 | 0.010 | 0.279 | 0.637 | 19.504 | < .001 | 0.008 | 19.504 |
| Frontal pole | 0.002 | 0.004 | 0.680 | 0.823 | 0.470 | 0.493 | < .001 | 1.000 | 0.001 | 0.004 | 0.718 | 0.823 | 0.073 | 0.787 | < .001 | 1.000 |
| Fusiform | 0.006 | 0.007 | 0.399 | 0.713 | 10.716 | 0.001 | 0.004 | 10.716 | 0.009 | 0.009 | 0.347 | 0.673 | 18.351 | < .001 | 0.007 | 18.351 |
| Inferior parietal | 0.007 | 0.004 | 0.076 | 0.438 | 2.226 | 0.136 | < .001 | 2.226 | 0.012 | 0.005 | 0.025 | 0.277 | 5.465 | 0.019 | 0.001 | 5.465 |
| Inferior temporal | 0.008 | 0.004 | 0.021 | 0.277 | 0.807 | 0.369 | < .001 | 1.000 | 0.013 | 0.005 | 0.009 | 0.277 | 4.258 | 0.039 | 0.001 | 4.258 |
| Insula | 0.001 | 0.004 | 0.901 | 0.936 | 2.271 | 0.132 | < .001 | 2.271 | < .001 | 0.005 | 0.995 | 0.995 | 4.669 | 0.031 | 0.001 | 4.669 |
| Isthmus cingulate | 0.006 | 0.004 | 0.103 | 0.438 | 0.001 | 0.978 | < .001 | 1.000 | 0.006 | 0.004 | 0.086 | 0.438 | 0.771 | 0.380 | < .001 | 1.000 |
| Lateral occipital | 0.001 | 0.004 | 0.702 | 0.823 | 0.054 | 0.816 | < .001 | 1.000 | 0.007 | 0.004 | 0.093 | 0.438 | 2.605 | 0.107 | 0.001 | 2.605 |
| Lateral orbitofrontal | 0.002 | 0.004 | 0.608 | 0.804 | 0.149 | 0.700 | < .001 | 1.000 | -0.001 | 0.004 | 0.887 | 0.936 | 0.831 | 0.362 | < .001 | 1.000 |
| Lingual | 0.001 | 0.004 | 0.726 | 0.823 | 1.439 | 0.230 | < .001 | 1.439 | 0.005 | 0.008 | 0.539 | 0.781 | 12.731 | < .001 | 0.005 | 12.731 |
| Medial orbitofrontal | -0.001 | 0.004 | 0.874 | 0.936 | 1.258 | 0.262 | < .001 | 1.258 | -0.001 | 0.004 | 0.717 | 0.823 | 0.604 | 0.437 | < .001 | 1.000 |
| Middle temporal | 0.012 | 0.005 | 0.021 | 0.277 | 4.688 | 0.030 | 0.001 | 4.688 | 0.011 | 0.007 | 0.130 | 0.491 | 11.214 | 0.001 | 0.004 | 11.214 |
| Paracentral | -0.003 | 0.005 | 0.519 | 0.781 | 2.953 | 0.086 | 0.001 | 2.953 | < .001 | 0.005 | 0.933 | 0.946 | 3.488 | 0.062 | 0.001 | 3.488 |
| Parahippocampal | -0.002 | 0.005 | 0.681 | 0.823 | 4.521 | 0.033 | 0.001 | 4.521 | 0.008 | 0.006 | 0.215 | 0.619 | 7.107 | 0.008 | 0.002 | 7.107 |
| Pars opercularis | 0.005 | 0.004 | 0.152 | 0.531 | 0.603 | 0.437 | < .001 | 1.000 | 0.002 | 0.005 | 0.761 | 0.848 | 4.642 | 0.031 | 0.001 | 4.642 |
| Pars orbitalis | 0.006 | 0.005 | 0.199 | 0.619 | 3.111 | 0.078 | 0.001 | 3.111 | 0.004 | 0.004 | 0.318 | 0.664 | 0.090 | 0.764 | < .001 | 1.000 |
| Pars triangularis | 0.008 | 0.005 | 0.103 | 0.438 | 3.004 | 0.083 | 0.001 | 3.004 | 0.003 | 0.005 | 0.442 | 0.759 | 2.556 | 0.110 | 0.001 | 2.556 |
| Pericalcarine | -0.008 | 0.004 | 0.030 | 0.277 | 0.534 | 0.465 | < .001 | 1.000 | -0.004 | 0.004 | 0.326 | 0.664 | < .001 | 0.990 | < .001 | 1.000 |
| Postcentral | 0.010 | 0.005 | 0.033 | 0.277 | 3.428 | 0.064 | 0.001 | 3.428 | 0.004 | 0.004 | 0.379 | 0.713 | 3.017 | 0.082 | 0.001 | 3.017 |
| Posterior cingulate | 0.009 | 0.004 | 0.015 | 0.277 | 0.631 | 0.427 | < .001 | 1.000 | 0.004 | 0.004 | 0.242 | 0.619 | 0.719 | 0.397 | < .001 | 1.000 |
| Precentral | 0.009 | 0.008 | 0.231 | 0.619 | 12.873 | < .001 | 0.004 | 12.873 | 0.009 | 0.009 | 0.302 | 0.662 | 16.477 | < .001 | 0.006 | 16.477 |
| Precuneus | 0.004 | 0.005 | 0.447 | 0.759 | 3.174 | 0.075 | 0.001 | 3.174 | 0.007 | 0.005 | 0.211 | 0.619 | 4.814 | 0.028 | 0.001 | 4.814 |
| Rostral anterior cingulate | 0.004 | 0.004 | 0.332 | 0.664 | 0.136 | 0.712 | < .001 | 1.000 | 0.003 | 0.004 | 0.539 | 0.781 | 2.068 | 0.150 | < .001 | 2.068 |
| Rostral middle frontal | 0.004 | 0.003 | 0.273 | 0.637 | 0.893 | 0.345 | < .001 | 1.000 | 0.002 | 0.004 | 0.593 | 0.804 | 0.582 | 0.446 | < .001 | 1.000 |
| Superior frontal | 0.003 | 0.005 | 0.554 | 0.784 | 3.958 | 0.047 | 0.001 | 3.958 | 0.003 | 0.005 | 0.540 | 0.781 | 3.380 | 0.066 | 0.001 | 3.380 |
| Superior parietal | 0.002 | 0.005 | 0.597 | 0.804 | 3.099 | 0.078 | 0.001 | 3.099 | 0.004 | 0.004 | 0.392 | 0.713 | 2.657 | 0.103 | 0.001 | 2.657 |
| Superior temporal | 0.010 | 0.009 | 0.281 | 0.637 | 18.605 | < .001 | 0.006 | 18.605 | 0.014 | 0.010 | 0.156 | 0.531 | 25.632 | < .001 | 0.008 | 25.632 |
| Supramarginal | 0.009 | 0.004 | 0.012 | 0.277 | 1.111 | 0.292 | < .001 | 1.111 | 0.007 | 0.006 | 0.238 | 0.619 | 7.237 | 0.007 | 0.002 | 7.237 |
| Temporal pole | 0.005 | 0.007 | 0.502 | 0.781 | 8.678 | 0.003 | 0.003 | 8.678 | 0.007 | 0.010 | 0.474 | 0.773 | 18.940 | < .001 | 0.007 | 18.940 |
| Transverse temporal | -0.002 | 0.004 | 0.668 | 0.823 | 1.374 | 0.241 | < .001 | 1.374 | 0.003 | 0.004 | 0.478 | 0.773 | 2.050 | 0.152 | < .001 | 2.050 |

*Table S25* Quadratic model results for $g \sim$ directional gene component scores.

| $g \sim$ | Component 1 | Component 2 |
|---|---|---|
| Volume | β = -0.432 *SE* = 0.003), *p* = .0009 | β = -0.559 (*SE* = 0.002), *p* = 3.55e-06 |
| Surface area | β = -0.356 (*SE* = 0.003), *p* = .007 | β = -0.496 (*SE* = 0.002), *p* = 5.54e-06 |
| Thickness | β = -0.168 (*SE* = 0.004), *p* = .212 | β = -0.218 (*SE* = 0.003), *p* = .089 |

*Table S26* Regression results of the association between regional mean cell type profiles and *g*-morphometry betas, after correcting for the two major components of gene expression.

| Mean | Cell type | β | *SE* | *t* | *p* | *FDR Q* |
|---|---|---|---|---|---|---|
| Volume | Astrocyte | -0.110 | 0.050 | -2.207 | 0.031 | 0.097 |
| Area | Astrocyte | -0.094 | 0.051 | -1.859 | 0.068 | 0.141 |
| Thickness | Astrocyte | -0.099 | 0.051 | -1.947 | 0.056 | 0.136 |
| Volume | CA1Pyramidal | 0.020 | 0.056 | 0.356 | 0.723 | 0.848 |
| Area | CA1Pyramidal | 0.046 | 0.056 | 0.814 | 0.419 | 0.565 |
| Thickness | CA1Pyramidal | -0.016 | 0.057 | -0.278 | 0.782 | 0.848 |
| Volume | Endothelial | -0.183 | 0.084 | -2.186 | 0.032 | 0.097 |
| Area | Endothelial | -0.148 | 0.085 | -1.741 | 0.086 | 0.146 |
| Thickness | Endothelial | -0.171 | 0.086 | -2.001 | 0.050 | 0.134 |
| **Volume** | **Ependymal** | **-0.200** | **0.054** | **-3.668** | **0.000** | **0.007** |
| Area | Ependymal | -0.143 | 0.057 | -2.492 | 0.015 | 0.069 |
| **Thickness** | **Ependymal** | **-0.244** | **0.053** | **-4.626** | **0.000** | **0.001** |
| Volume | Interneuron | -0.036 | 0.079 | -0.450 | 0.654 | 0.803 |
| Area | Interneuron | 0.115 | 0.078 | 1.469 | 0.147 | 0.233 |
| Thickness | Interneuron | -0.138 | 0.079 | -1.745 | 0.086 | 0.146 |
| **Volume** | **Microglia** | **-0.155** | **0.054** | **-2.894** | **0.005** | **0.035** |
| **Area** | **Microglia** | **-0.175** | **0.053** | **-3.322** | **0.001** | **0.013** |
| Thickness | Microglia | -0.003 | 0.058 | -0.056 | 0.956 | 0.981 |
| Volume | Mural | -0.131 | 0.073 | -1.785 | 0.079 | 0.146 |
| Area | Mural | -0.070 | 0.075 | -0.935 | 0.353 | 0.502 |
| Thickness | Mural | -0.192 | 0.072 | -2.650 | 0.010 | 0.055 |
| Volume | Oligodendrocyte | 0.107 | 0.086 | 1.245 | 0.218 | 0.327 |
| Area | Oligodendrocyte | 0.002 | 0.087 | 0.024 | 0.981 | 0.981 |
| Thickness | Oligodendrocyte | 0.164 | 0.086 | 1.912 | 0.060 | 0.136 |
| Volume | S1Pyramidal | 0.069 | 0.099 | 0.691 | 0.492 | 0.632 |
| Area | S1Pyramidal | 0.223 | 0.096 | 2.325 | 0.023 | 0.090 |
| Thickness | S1Pyramidal | -0.028 | 0.101 | -0.274 | 0.785 | 0.848 |

### Supplementary Text 1: Meta-analytic mean morphometry profile associations with general and specific patterns of human brain gene expression

While the main analysis focuses on regional *g*-associations, in this supplementary analysis, we provide a complimentary analysis on regional morphometry mean profiles. We calculate meta-analytic regional morphometry means (total $N$ = 40,292[1]) and report correlations with the major components of gene expression. Then, while correcting for these components, we identify specific cell-type-morphometry associations and individual gene-morphometry associations.

Previously, with the virtual histology approach, Paus and colleagues have found associations between regional cortical thickness profiles, and those of astrocytes ($r$ = 0.323, $p$ = 1.68e-03) and CA1 pyramidal cells ($r$ = 0.287, $p$ = 2.40e-04) in a sample of 507 young healthy men[18]. In that study, whilst the correlation for microglia was not statistically significant, it was moderate ($r$ = 0.234, $p$ = .066). Similar results were reported in another study from the same group using a different sample[19]. It should be noted that they used a more stringent two-step consistency check for retaining genes than we used in the current study (this resulted in 2511 genes, as opposed to 8235 retained in the current study). Given that it is these three cell types that have differential loading distributions to baseline on Component 1 in the current study (see supplementary data file), we might predict that controlling for this component would eliminate these apparent cell-specific associations. Taking the present approach, it is

[1] *Note* the UKB sample is slightly larger in this analysis than that used to calculate regional *g*-associations in the main text (UKB $N$ = 39,250, compared to $N$ = 37,840). This is due to slightly different exclusion criteria being used (here, there are fewer participant exclusions as criteria were only based on self-report methods at UKB assessment, and not also official records). This sampling difference is very unlikely to affect the mean results – this is supported by the high correlations in Table SA.

possible to identify cell types that are uniquely associated with regional volume, surface area and thickness, beyond their involvement in major components of cortical gene expression. As in the main analysis, the same approach is extended to the profiles of individual genes.

### Results

There were no within-cohort sex differences in the regional mean profiles (for all, $r$ = 1.00). There was also very high consistency in the regional mean profiles between the three cohorts (all $r > .91$, all $p < .0001$, see *Table SA*). The meta-analytic mean results are shown in *Figures SA-D*, and the values are reported in the supplementary data file.

*Table SA* Cross-cohort correlations for regional mean profiles in volume, surface area and thickness (all $p < .0001$).

| Cohort comparison | *r* Volume | *r* Surface area | *r* Thickness |
|---|---|---|---|
| LBC–STRADL | 0.998 | 0.999 | 0.946 |
| STRADL–UKB | 0.999 | 0.998 | 0.943 |
| UKB–LBC | 0.998 | 0.998 | 0.917 |

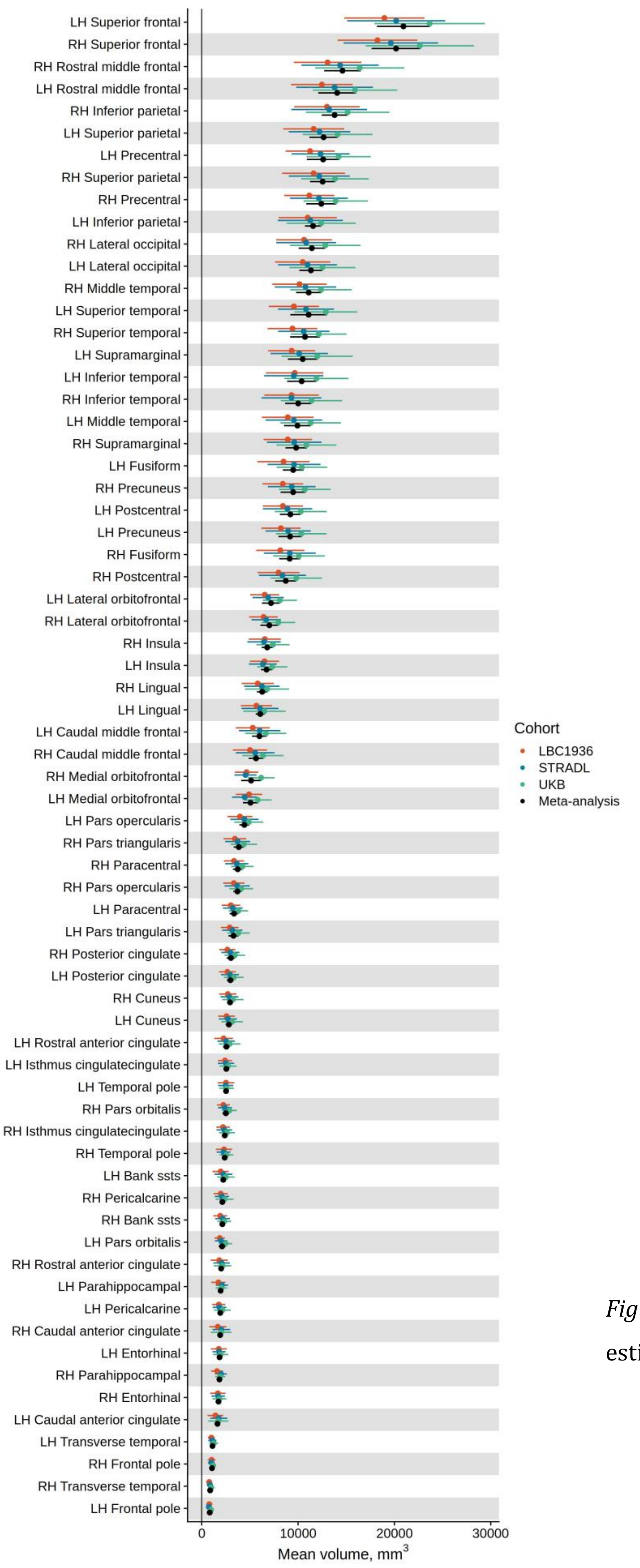


*Figure SA* Meta-analytic regional estimates of mean volume profiles

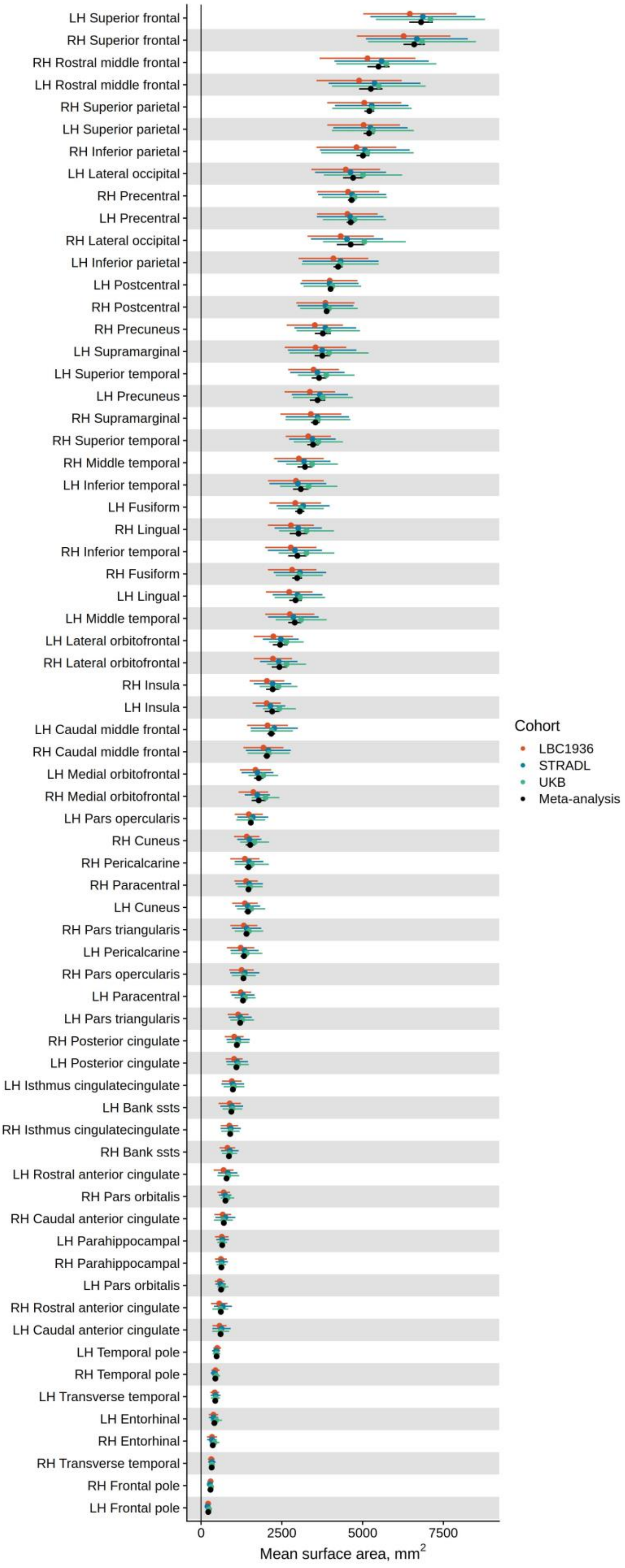


*Figure SB* Meta-analytic regional estimates of mean surface area profiles

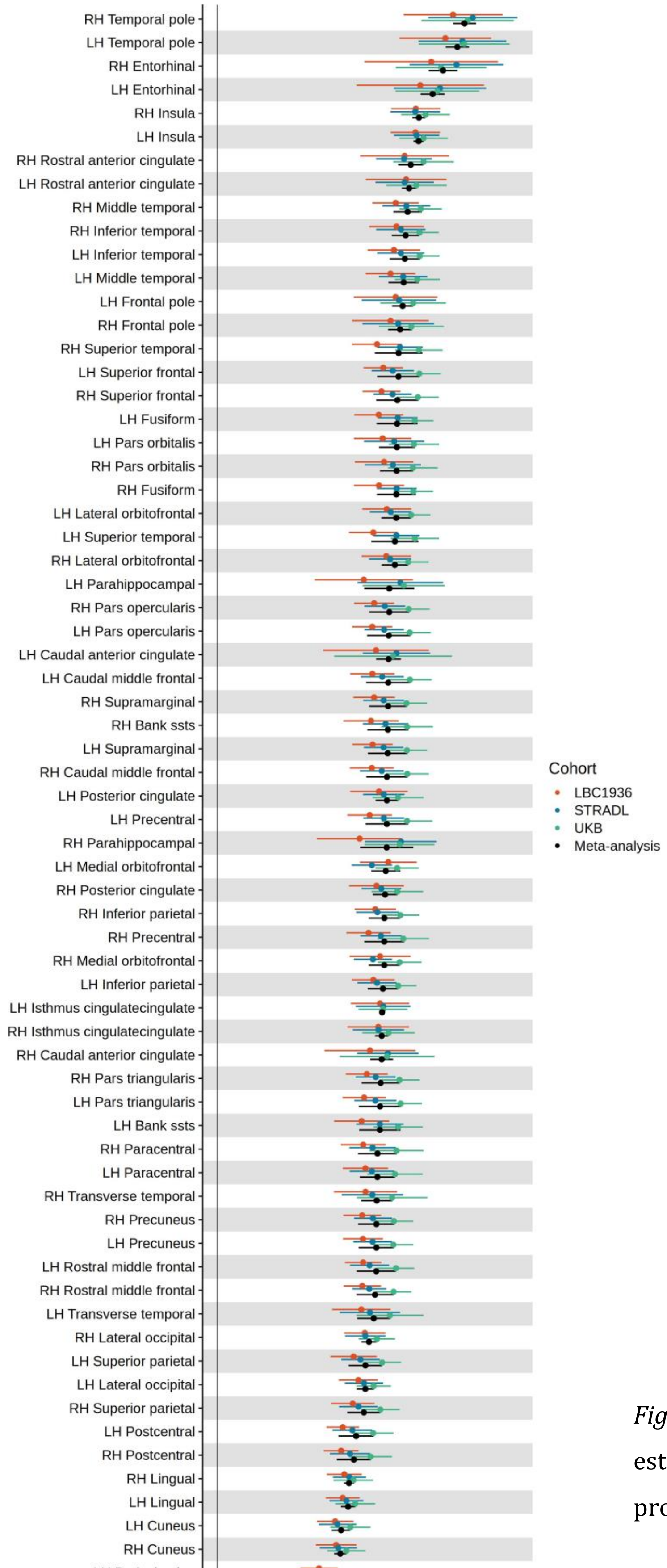


*Figure SC* Meta-analytic regional estimates of mean thickness profiles

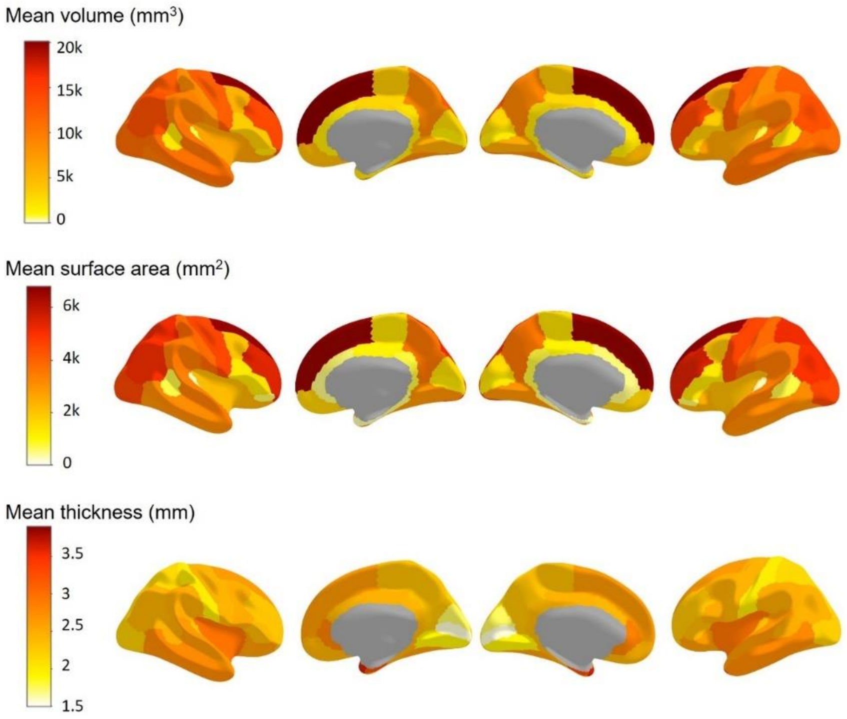


*Figure SD* Meta-analytic regional mean profiles of volume, surface area and thickness mapped to the cortex.

### Correlations between morphometry measures

The regional mean profiles of volume and surface area were closely positively correlated ($r$ = 0.976, $p$ < 2.2e-16), whilst thickness and surface area had a moderate and non-significant negative association ($r$ = -0.226, $p$ = .064), and there was no correlation between volume and thickness ($r$ = -0.055, $p$ = .658).

### Associations between regional mean morphometry profiles and *g*-associations

The *g*-associations (reported in the main analysis) were positively associated with their regional mean profiles for all three morphometry measures. The strongest correlation was for volume ($r$ = 0.709, $p$ = 1.35e-11), followed by surface area ($r$ = 0.614, $p$ = 2.58e-08), and then thickness ($r$ = 0.313, $p$ = .009). In other words, regions with stronger *g*-

associations tend to be larger in terms of volume and surface area, and also tend to be thicker.

### Associations between regional mean morphometry profiles and the non-absolute component scores or regional overall mean expression profiles

Whilst there are no particularly compelling associations between the regional mean profiles for volume and surface area and the non-absolute component scores, there are strong correlations for thickness (see *Table SB*). The thicker a region is, the more strongly it falls on the regulation side of each gene expression component – cell signalling and modification ($r$ = 0.764, $p$ = 3.67e-14); and transcription factors ($r$ = -0.799, $p$ = 3.132e-16). The regional mean thickness profiles are also moderately associated with the regional gene expression means ($r$ = 0.410, $p$ = .0005) suggesting that simply higher regional levels of gene expression tend to occur in thicker regions.

*Table SB* Correlations between the regional mean morphometry profiles and the **non-absolute** component scores and mean expression values.

| Mean profile | Component 1 | Component 2 | Mean expression |
|---|---|---|---|
| Volume | $r$ = -0.082, $p$ = .504 | $r$ = 0.111, $p$ = .368 | $r$ = -0.081, $p$ = .513 |
| Surface area | $r$ = -0.230, $p$ = .059 | $r$ = 0.245, $p$ = .044 | $r$ = -0.144, $p$ = .243 |
| Thickness | $r$ = 0.764, $p$ = 3.68e-14 | $r$ = -0.799, $p$ = 3.13e-16 | $r$ = 0.410, $p$ = .0005 |

### Associations between the regional mean morphometry profiles and the absolute component scores

Turning to the absolute component scores, the regions that are most important for defining the two major components of gene expression tend to be moderately smaller in terms of volume and surface area and slightly less thick (see *Table SC*).

*Table SC* Correlations between the regional mean morphometry measures and the **absolute** component scores and mean expression values.

| Mean profile | Component 1 | Component 2 |
|---|---|---|
| Volume | $r$ = -0.369, $p$ = .002 | $r$ = -0.471, $p$ = 5.083e-05 |
| Surface area | $r$ = -0.288, $p$ = .017 | $r$ = -0.401, $p$ = .0007 |
| Thickness | $r$ = -0.255, $p$ = .036 | $r$ = -0.162, $p$ = .187 |

### Associations between regional mean morphometry profiles and specific cell types

Then, correcting for the regional component profiles, we tested for associations between the regional morphometry means and the mean profiles for each of the 9 cell types. The only associations for which FDR $Q < .05$ were for $g$-volume and $g$-surface area with ependymal cells: $g$-volume: β = -0.210, *SE* = 0.054, $p < .001$, $g$-surface area: β = -0.207, *SE* = 0.056, $p < .001$), see *Table SD*. These associations were small-to-moderate and negative, suggesting that ependymal cells have lower expression levels in larger cortical structures.

*Table SD* Regression results of the association between regional mean cell type profiles and regional morphometry means, after correcting for the two major components of gene expression.

| Mean | Cell type | β | *SE* | *t* | *p* | *FDR Q* |
|---|---|---|---|---|---|---|
| Volume | Astrocyte | -0.620 | 0.292 | -2.125 | 0.037 | 0.253 |
| Area | Astrocyte | -0.094 | 0.052 | -1.789 | 0.078 | 0.286 |
| Thickness | Astrocyte | -0.050 | 0.103 | -0.488 | 0.627 | 0.709 |
| Volume | CA1Pyramidal | -0.027 | 0.057 | -0.479 | 0.634 | 0.709 |
| Area | CA1Pyramidal | -0.048 | 0.058 | -0.829 | 0.410 | 0.673 |
| Thickness | CA1Pyramidal | 0.157 | 0.111 | 1.414 | 0.162 | 0.318 |
| Volume | Endothelial | -0.122 | 0.086 | -1.418 | 0.161 | 0.318 |
| Area | Endothelial | -0.130 | 0.088 | -1.473 | 0.146 | 0.318 |
| Thickness | Endothelial | -0.078 | 0.173 | -0.450 | 0.654 | 0.709 |
| **Volume** | **Ependymal** | **-0.210** | **0.054** | **-3.871** | **< .001** | **0.007** |
| **Area** | **Ependymal** | **-0.207** | **0.056** | **-3.678** | **< .001** | **0.007** |
| Thickness | Ependymal | -0.292 | 0.114 | -2.565 | 0.013 | 0.114 |
| Volume | Interneuron | 0.061 | 0.080 | 0.766 | 0.446 | 0.673 |
| Area | Interneuron | 0.052 | 0.082 | 0.628 | 0.532 | 0.709 |
| Thickness | Interneuron | -0.012 | 0.159 | -0.075 | 0.940 | 0.940 |
| Volume | Microglia | -0.102 | 0.056 | -1.818 | 0.074 | 0.286 |
| Area | Microglia | -0.101 | 0.058 | -1.751 | 0.085 | 0.286 |
| Thickness | Microglia | 0.055 | 0.113 | 0.488 | 0.627 | 0.709 |
| Volume | Mural | -0.036 | 0.075 | -0.475 | 0.636 | 0.709 |
| Area | Mural | -0.021 | 0.078 | -0.266 | 0.791 | 0.822 |
| Thickness | Mural | -0.201 | 0.147 | -1.365 | 0.177 | 0.318 |
| Volume | Oligodendrocyte | 0.124 | 0.086 | 1.445 | 0.153 | 0.318 |
| Area | Oligodendrocyte | 0.130 | 0.089 | 1.468 | 0.147 | 0.318 |
| Thickness | Oligodendrocyte | -0.131 | 0.172 | -0.763 | 0.448 | 0.673 |
| Volume | S1Pyramidal | 0.171 | 0.098 | 1.752 | 0.085 | 0.286 |
| Area | S1Pyramidal | 0.142 | 0.102 | 1.398 | 0.167 | 0.318 |
| Thickness | S1Pyramidal | 0.089 | 0.198 | 0.447 | 0.656 | 0.709 |

### Associations between regional mean morphometry profiles and individual genes

There were 639 individual genes that had *FDR Q* < .05 for associations with volume mean profiles, 604 with surface area and 153 with thickness. The full FDR-corrected results of this analysis are in the supplementary data file. 561 genes had *Q* < .05 for both volume and surface area, 6 for both volume and thickness and 3 for surface area and thickness. There were three genes that were significant for all three morphometry measure mean profiles: *CCDC19*, *FSTL4* and *NPTX2* (see *Table SE*).

*Table SE* Standardised β values for the three individual genes for which *FDR Q* < .05 for all three morphometry measure mean profiles, after correcting for the two major components of gene expression.

| Mean profile | *CCDC19* | *FSTL4* | *NPTX2* |
|---|---|---|---|
| Volume | β = -0.412 | β = 0.289 | β = 0.305 |
| Surface area | β = -0.436 | β = 0.264 | β = 0.288 |
| Thickness | β = 0.715 | β = 0.510 | β = 0.561 |

### Brief discussion

In this supplementary analysis, we provide a robust meta-analysis of regional cortical means for three morphometry measures: volume, surface area and thickness. There was very high consistency (all *r* > 0.91) in these regional profiles between three large cohorts, demonstrating their reliability.

Regions that are larger in terms of volume and surface area, and thicker tend positively associated with *g*-associations and tend not to be those that define the two components of gene expression.

We found that regional thickness is moderately positively associated with regional mean expression levels, and that thicker regions fall strongly on the regulation side of both the cell-signalling and modification and transcription factors axes.

We then tested for associations between individual cell type profiles and regional morphometric means, after controlling for the two major components of gene expression. We identified ependymal cells as being associated with the mean volume and surface area beyond the two major components. These associations were moderately negative, suggesting that ependymal cells have lower expression levels in larger cortical structures. As predicted, the previously-reported associations between cortical thickness and astrocytes, microglia and CA1 pyramidal cells were not significant, whilst correcting for the two major components of gene expression. This is likely because thickness has strong correlations with both components (Component 1 $r$ = 0.764, Component 2 $r$ = -0.799), and microglia, astrocytes and CA1 pyramidal cells load differentially onto Component 1 than other genes (see supplementary data file). Therefore, such associations are likely to be best-explained in the context of the two components of gene expression, instead of by the mean expression values of specific cell types alone.

Finally, we calculated unique individual gene-morphometry associations, beyond the two major components. Three genes had significant associations with all three morphometry measures: *CCDC19*, *FSTL4* and *NPTX2*. *CCDC19* has previously been found to suppress cell growth[20]. It has a negative association with both volume and surface area, potentially suggesting that its cell growth suppression is less desirable in regions with that are larger in terms of volume and surface area. Conversely, it has a positive association with regional thickness profiles, indicating that it might suppress cell growth in thicker regions. *FSTL4* is thought to play an important role in regulating neurotrophic factors which regulate morphological plasticity[21] - this could explain its positive associations with all three morphometry measures – larger structures generally require more morphological plasticity. Finally, *NPTX2* appears to be an especially important gene for regional brain organisation – it is positively associated with all three morphometry measures, and it is positively associated with all three *g*-associations from the main body of the current paper. It is thought to play a critical role in synaptic plasticity and protection, as well as being a key gene in epilepsy, Parkinson's Disease and Alzheimer's Disease[22].
